# The rich evolutionary history of the ROS metabolic arsenal shapes its mechanistic plasticity at the onset of metazoan regeneration

**DOI:** 10.1101/2024.07.25.605162

**Authors:** Aurore Vullien, Aldine Amiel, Loeiza Baduel, Dilara Diken, Cécile Renaud, Michel Vervoort, Eric Röttinger, Eve Gazave

## Abstract

Regeneration, the ability to restore body parts after injury, is widespread in metazoans; however, the underlying molecular and cellular mechanisms involved in this process remain largely unknown, and its evolutionary history is consequently unresolved. In the last decade, ROS have emerged as shared actors that trigger apoptosis and cell proliferation to drive regenerative success in a few metazoan models. However, it is not known whether the contribution of ROS to regeneration relies on conserved mechanisms in animals.

Here we performed a comparative genomic analysis of ROS metabolism actors across metazoans, and carried out a comparative study for the deployment and roles of ROS during regeneration in two different research models: the annelid Platynereis dumerilii and the cnidarian Nematostella vectensis. We established that the vast majority of metazoans possess a core redox kit allowing for the production and detoxification of ROS, and overall regulation of ROS levels. However, the precise composition of the redox arsenal can vary drastically from species to species, suggesting that evolutionary constraints apply to ROS metabolism functions rather than precise actors. We found that ROS are produced during and are necessary for regeneration in both Platynereis and Nematostella. However, we also uncovered different enzymatic activities underlying ROS dynamics, as well as distinct effects of ROS signalling on injury-induced apoptosis and cell proliferation in the two species. We conclude that, while ROS are a robust feature of metazoan regeneration, their production and contribution to this phenomenon may depend on plastic molecular mechanisms.

## Introduction

Reactive Oxygen Species (ROS) have been considered for a long time as deleterious components to the cellular environment, a reputation owed to the fact that these very reactive molecules, in particular hydroxyl radicals (HO^-^), can cause the alteration of all sorts of molecules (proteins, lipids, nucleic acids…) (Di Meo et al. 2016; Sies et al. 2017). As a result, oxidative stress, *i.e.* the excess of reactive oxidants, has long been identified as a deleterious factor in various diseases (cancer, age-related afflictions …) (Halliwell and Gutteridge 1984).

However, this rather negative vision was revised as a growing number of studies revealed an unsuspected role of ROS as signalling actors (Suzuki et al. 1997), with hydrogen peroxide (H_2_O_2_) in particular emerging as a second messenger. Hence, their physiological role has since been established in processes as varied as cell proliferation, cell death or survival, differentiation, migration and metabolic regulation (Covarrubias et al. 2008; Salas-Vidal et al. 2020). ROS perform these functions mainly through reversible thiol oxidation, modifying the conformation, activity and localization of diverse targets such as kinases and phosphatases, receptors, transcription factors, either directly or through protein relays (Winterbourn 2013). In light of these findings, a new vision emerged, in which excessive amounts of ROS do cause cellular damage (oxidative distress), but innocuous concentrations of ROS allow for physiological signalling (oxidative eustress) (Sies et al. 2017), with sensing and signalling systems involving ROS described particularly in mammalians, yeasts and bacteria (Brigelius-Flohé and Flohé 2011).

Consequently, ROS are subjected to a complex metabolism to maintain cellular redox homeostasis, balancing ROS-producing and antioxidant systems, that has been extensively characterised in vertebrates (de Almeida et al. 2022). The different ROS molecules are produced sequentially (Figure 1). The “original” ROS, the superoxide anion (O ^-^), results from the reduction of dioxygen. It is produced either by complexes of the mitochondrial electron transport chain or by membrane Nox/Duox enzymes, and is dismutated by SOD enzymes into H_2_O_2_ in all compartments (mitochondria, reticulum, extracellular) (Lennicke and Cochemé 2021). The multiple locations of O_2_ and H_2_O_2_-producing systems constitute an asset for precise and specific ROS signalling (Herb et al. 2021).

**Figure 1:**
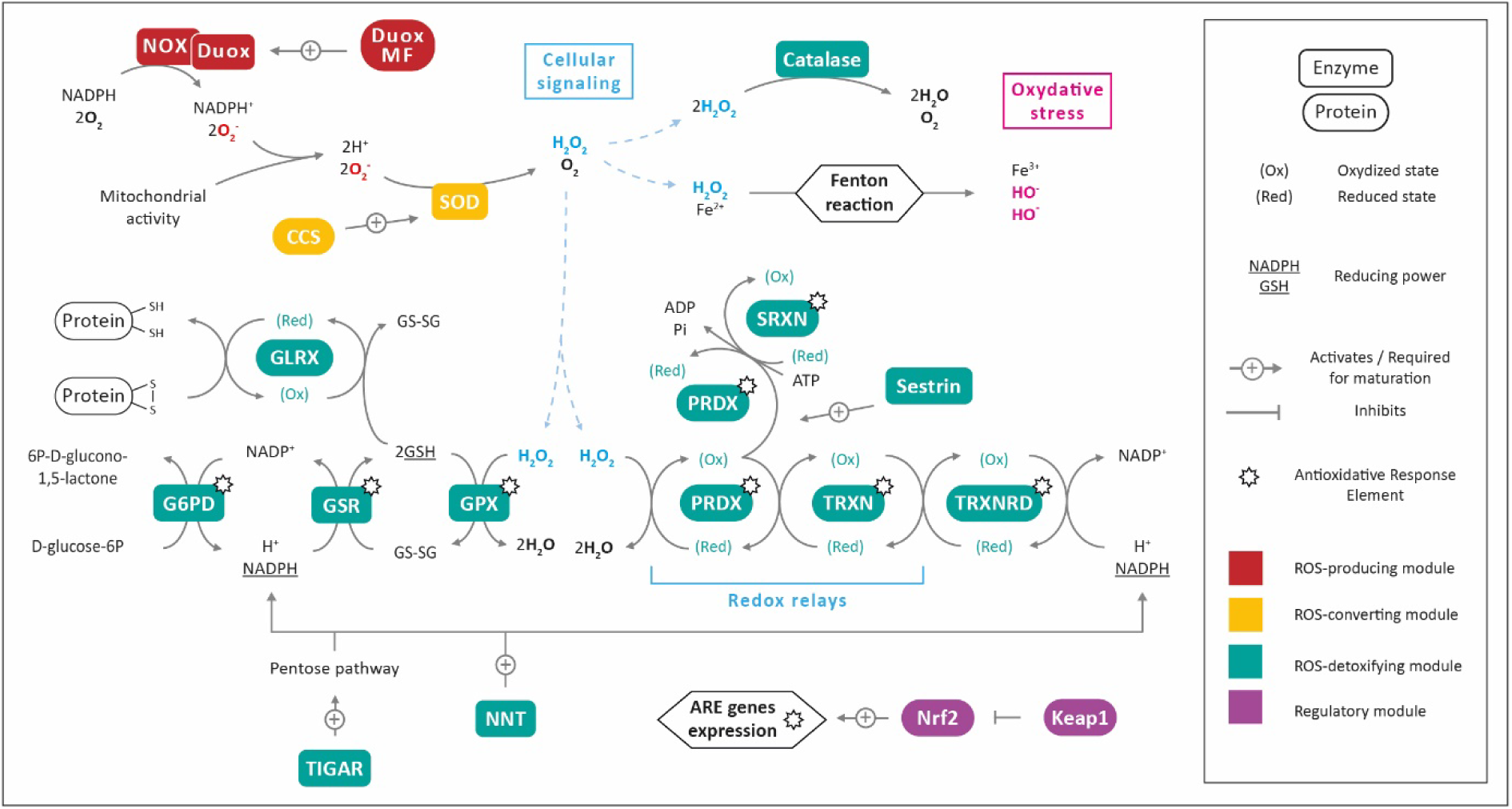
ROS metabolism. Primary ROS (superoxide, O2-) are produced by Nox/Duox enzymes or by the mitochondrial respiratory chain. It is converted to H2O2 by superoxide dismutases (SOD). H2O2 reaction with metal cations (Fe2+ or Cu+) causes the formation of HO-radicals, a source of molecular damage (DNA, proteins, lipids). On the other hand, H2O2 contributes to cellular signalling by affecting the thiol oxidation status of target proteins, acting in some cases through protein redox relay such as peroxiredoxins (PRDX) and thioxiredoxin (TXRN). It is detoxified to water by catalase and peroxidase (GPX) enzymes. ROS detoxification involves enzymes responsible for the production of reducing power in the form of reduced glutathione (GSH) and NADPH, such as Glucose-6-phosphate dehydrogenase (G6PD), glutathione-disulfide reductase (GSR), NAD(P) transhydrogenase (NNT), TP53 Induced Glycolysis Regulatory Phosphatase (TIGAR), or for restoring the reduced status of other key proteins, such as thioxiredoxin reductase (TXRNRD), sulfiredoxin (SRXN), glutaredoxin (GLRX). The global antioxidant response is under the control of the transcription factor Nrf2, targeting genes with an Antioxidant Response Element (ARE). Under homeostatic conditions, Nrf2 is constitutively associated with and targeted to degradation by Keap1 in the cytoplasm, an association disrupted by oxidative stress. Other abbreviations: copper chaperone for SOD (CCS), Duox maturation factor (Duox MF).

H_2_O_2_ is the main actor of this signalling, however its accumulation can cause the formation of the highly reactive HO^-^ (through the Fenton reaction). To prevent this, the redox balance is kept in check by antioxidant enzymes (catalase; glutathione peroxidase or GPX), metabolites providing reducing power (glutathione, NADPH), and numerous proteins acting as redox relays (thioredoxins or TRX; peroxiredoxins or PRDX etc), allowing the detoxification of H_2_O_2_ into H_2_O and O_2_. The expression of several aforementioned antioxidant actors is under the control of the transcription factor Nrf2. In physiological conditions, its activity is suppressed by its inhibitor, Keap1, which targets Nrf2 for degradation. Under oxidative stress, their association is disrupted, allowing for the transcription of genes with an Antioxidant Response Element (Figure 1). However, this global overview of ROS metabolism results mostly from studies carried on mammalian models or cell cultures. Some species are known to lack potential key players, such as *C. elegans* that lack the *Keap1* gene and regulates its Nrf2 ortholog SKN-1 through an alternate mechanism (Choe et al. 2009), suggesting a variability of the redox arsenal across species, maintaining the core functions of ROS metabolism.

While extensively studied in vertebrates, ROS metabolism and signalling systems are reported in many metazoan lineages, as well as plants, yeasts and bacteria, suggesting that H_2_O_2_ as a cell signalling molecule is an ancient and conserved feature (D’Autréaux and Toledano 2007). Indeed, it appears that key components of the ROS metabolism (Nox/Duox, Sod, catalase, GPX) originated as early as 4 billion years ago (Inupakutika et al. 2016), meaning that the potential of ROS as signalling molecules might have been harnessed very early during life history (Taverne et al. 2018). Additional protective mechanisms against oxidative stress might still have emerged following the Great Oxidation Event caused by the emergence of photosynthesis about 2,5-2,2 billion years ago (Lyons et al. 2014). In particular, photosynthetic but also photosymbiotic organisms, such as acoels and cnidarians, need to cope with high intra-cellular levels of ROS and might thus have evolved specific antioxidative strategies (Richier et al. 2005). Hence, ROS have certainly played an extensive role in the evolution of living organisms.

Consistent with this evolutionary aspect, ROS have been shown to significantly contribute to numerous processes such as embryonic development and regeneration-independent wound healing (Rampon et al. 2018). During embryonic and larval development in various model species, ROS production is responsible for driving axons projections (Gauron et al. 2016), organ formation (e.g. zebrafish thyroid) (Chopra et al. 2023), tissue specification (Han et al. 2018) and triggering stem cells differentiation (Bigarella et al. 2014). In an injury context, ROS are best known to perform bactericidal functions and immune cells chemotaxis (Cordeiro and Jacinto 2013).

More recently, studies uncovered a key role of ROS in another developmental trajectory: restorative regeneration, *i.e.* the reformation of a body part lost to injury (Poss 2010; Bideau et al. 2021). Although regenerative abilities vary greatly among metazoan lineages, the regeneration process can be divided into three stereotypical modules in all species: (1) wound healing, (2) induction (mobilization of precursor cells taking part in regeneration), (3) morphogenesis (differentiation and patterning) (Galliot and Ghila 2010). An ever-growing number of studies report ROS production during the first module of regeneration of varied structures in a diversity of species: such as caudal fin in zebrafish (Gauron et al. 2013), *Xenopus* tadpole tail (Love et al. 2013), whole-body in *Hydra* (Suknovic et al. 2022) and flatworms (Pirotte et al. 2015) and *Drosophila* imaginal discs (Fogarty et al. 2016) (for review, see Vriz et al. 2014; Rampon et al. 2018).

In addition to their well-documented involvement during regeneration-independent wound healing (Cordeiro and Jacinto 2013), functional approaches suggest that 1) the implication of ROS during regeneration goes far beyond their bactericidal and chemotactic role and 2) they are mandatory for regeneration success in various model species. Indeed, ROS can trigger cell death and mitogenic events, and are later required for cell fate determination (Rampon et al. 2018; Vullien et al. 2021). These observations suggest that ROS could be conserved actors of regeneration initiation in association with apoptosis and compensatory cell proliferation, and a putative ROS -> apoptosis -> cell proliferation cascade, involving various signalling pathways (JNK, Wnt, MAPK, FGF) has been hypothesised (Diwanji and Bergmann 2018; Guerin et al. 2021). However, the available data remain fragmented and it is still unclear whether the role of ROS in regeneration initiation implies the same actors and relies on conserved mechanisms in metazoans or in contrast involves specific ROS metabolism members and cellular events in different lineages.

In this study, we chose to address these key questions by combining a fine investigation of the evolutionary history of ROS metabolism genes in metazoans with comparative functional studies in key regeneration research models of different phylogenetic positions within animals: the annelid *Platynereis dumerilii* (Lophotrochozoa, Errantia) (Özpolat et al. 2021; Schenkelaars and Gazave 2021) and the sea anemone *Nematostella vectensis* (Cnidaria, Anthozoa) (Layden et al. 2016; Röttinger 2021). This original approach revealed that the ROS machinery can vary drastically from one species to another at the protein level, but is constrained at the modules level. Furthermore, ROS signalling during regeneration in the two studied species uses different enzymatic players. Phenotypic analysis of ROS inhibition during regeneration in *Platynereis* and *Nematostella* highlighted that the previously proposed “ancestral” relationship between ROS, apoptosis, cell proliferation and regeneration is not confirmed. Instead, it revealed a mechanistic plasticity of ROS metabolism required for metazoan regeneration.

## Results

### Evolutionary history of the ROS metabolism arsenal

To establish how the main gene families involved in ROS metabolism (Figure 1) have evolved in metazoans, we explored genomic data at an unprecedented level, encompassing 85 species representing the main metazoan lineages, as well as four non-metazoan opisthokont species (Sup Table 1). We specifically searched for 9 multigenic families of the major modules of the redox arsenal: the ROS-producing module (Nox/Duox enzymes and the maturation factors DuoxMF), the ROS-converting module (Sod enzymes and the chaperone CCS), the detoxifying module (catalase and GPX) and the regulatory module (Nrf and Keap1). Using phylogenetic analyses, we established the presence and number of copies for all these proteins in all assessed species and inferred their orthology relationships (Figure 2; Sup Fig 1-9). By performing synteny and domain-composition analyses, prediction of 3D structures and protein interactions as well as ancestral state reconstructions (Figure 3; Sup Fig 10; Sup Table 2), we then explored in depth the evolutionary histories of each module.

**Figure 2:**
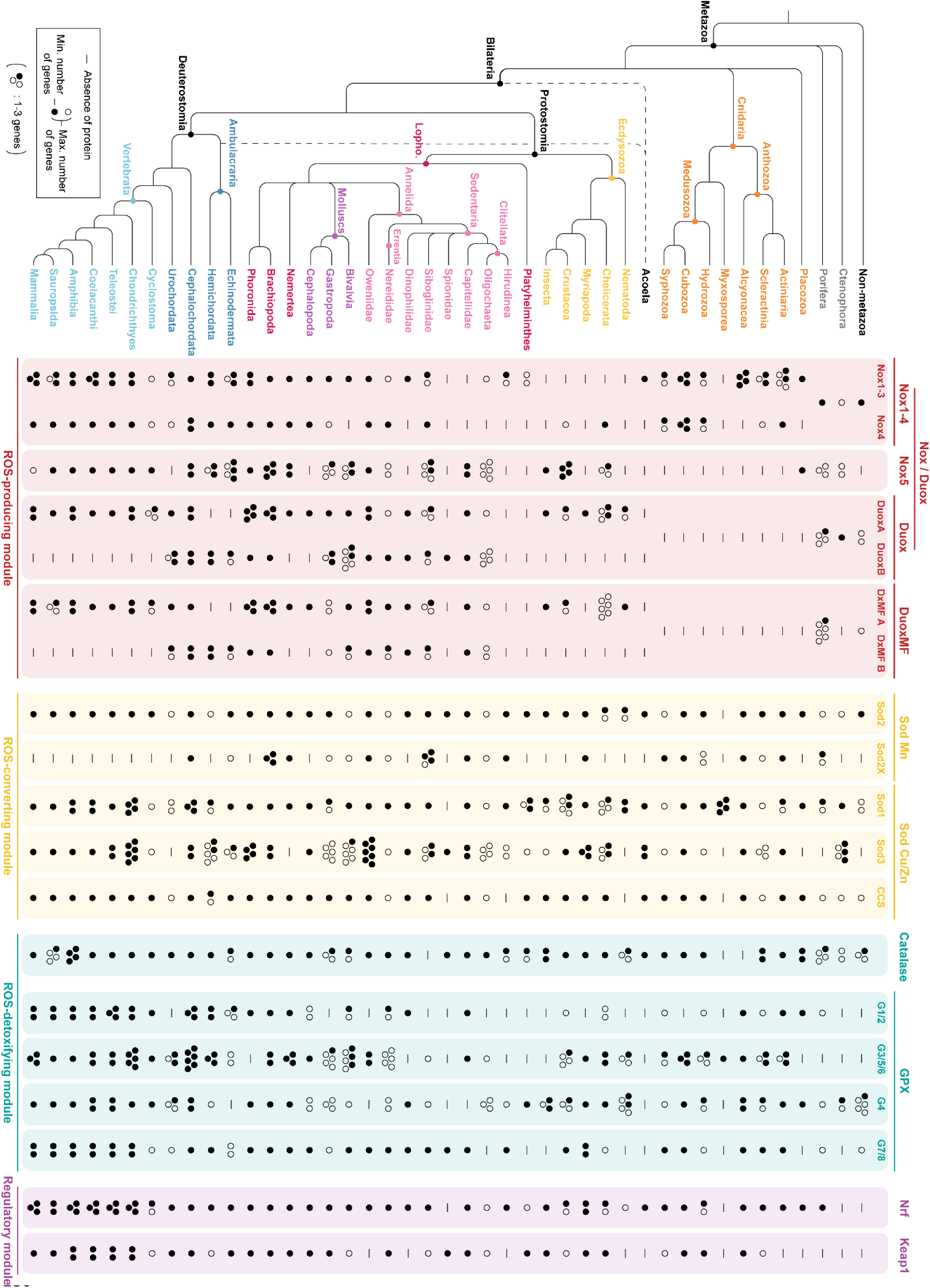
Overview of the ROS metabolism toolkit in metazoan lineages. A simplified metazoan phylogeny is displayed on the left, indicating the relationships between the main lineages to which the 89 species in the dataset belong. Controversial positions of acoels are indicated with dotted lines, and polytomies are used where uncertainties remain about the relationships between non-bilaterian groups. Major phylogenetic groups’ names are indicated at each corresponding node. Presence/absence and number of genes of the main ROS metabolism actors (columns, arranged by ROS metabolism functional modules) are indicated for each lineage (lines). Plain circles indicate the minimal number of genes found in all species of a lineage, plain and empty circles taken together indicate the maximum number of genes found (for example, non-metazoan species possess between 1 to 3 catalase genes). A dash indicates the absence of the gene. Sequences of the identified proteins and phylogenetic trees can be found in Sup Files 1-8 and Sup Fig 1-9. The protein names are colour coded according to the ROS-module to which they belong.

#### ROS-producing module: Nox/Duox and DuoxMF families

The Nox/Duox enzymes are a multigenic family of ROS-producing membrane proteins composed of Nox1-3, Nox4, Nox5 and Duox members (Kawahara et al. 2007). All of them are characterized by the presence of a Nox domain composed of a transmembrane ferric reductase element, a FAD-binding segment and a NADPH-binding region (Moghadam et al. 2021). Nox5 and Duox contain in addition two EF-hand motifs and Duox also possess an extra region homologous to a heme-containing peroxidase (schematic representations in Sup Fig 1A).

We identified 432 *Nox* and *Duox* genes in our dataset (list of protein sequences listed in Sup File 1). Phylogenetic analyses confirm the separation of *Nox* and *Duox* genes in four clades: Nox1-3, Nox4, Nox5 and Duox (Sup Fig 1A) (Kawahara et al. 2007). The number of Nox/Duox members *per* species varies largely ranging from 0 to 12 (Figure 2). We nevertheless observed some interesting points, such as the absence of Nox1-3 in ecdysozoans, Nox4 in about half of the protostome lineages and both Nox5 and Duox in cnidarians (Figure 2). We also determined several duplication events. A specific phylogeny of Nox1-3 and Nox4 sequences confirms that Nox4 result from the duplication of an ancestral Nox1-4 gene, while the Nox1, Nox2 and Nox3 proteins found in mammalians result from two rounds of duplication in vertebrates, possibly linked to documented whole-genome duplications in this clade (Dehal and Boore 2005) (Sup Fig 1B). Additional duplications of Nox1-3 are found in anthozoans and echinoderms, and of Nox4 in both cubozoan and scyphozoan cnidarians (Sup Fig 1B).

Other than the above-mentioned loss of Nox5 in cnidarians, the specific phylogeny of Nox5 revealed ancestral duplications in Ambulacraria, as well as some molluscs (mussels and scallops) (Sup Fig 1C). These results indicate that while ancestral *Nox1-4* and *Nox5* precede the emergence of metazoan, the duplication responsible for the acquisition of *Nox4* took place in the Last Common Ancestor (LCA) of eumetazoans (FIG 3 A).

A phylogenetic analysis focusing on Duox shows the presence of two clades – that we named Duox A and Duox B, both containing sequences of molluscs, annelids, cephalochordates and urochordates, suggesting an ancient duplication of *Duox* genes in bilaterians, while vertebrate and ecdysozoan sequences do not follow the same pattern (Sup Fig 1D). A further duplication of *Duox A* is observed in mammals, and of *Duox B* in molluscs.

An additional protein, named Duox Maturation Factor (or DuoxMF) is required for ROS-production by Duox, more specifically for its delivery to the cell membrane (Morand et al. 2009). We identified 121 *DuoxMF* sequences in our dataset that encode for proteins consisting of a Duox Maturation factor domain (list of protein sequences in Sup File 2, Sup Fig 2). Strikingly, their presence/absence pattern within Metazoa matches that of *Duox* proteins in the vast majority of the studied species (*i.e.* absence in Cnidaria, Ecdysozoa and Vertebrata), with the noticeable exception of Ctenophores. DuoxMF phylogenetic analyses indicate that *DuoxMF* was ancestrally duplicated in bilaterian and form two clades named DuoxMF A and DuoxMF B (Sup Fig 2).

Taken together, these results suggest that the LCA of metazoans possessed *Duox* and *DuoxMF* genes that were lost in cnidarian and duplicated in the LCA of bilaterians. This resulted in two copies of *Duox* (*Duox A* and *Duox B*) and two copies of *DuoxMF* (*DuoxMF A* and *DuoxMF B*). *Duox B/DuoxMF B* were then lost in ecdysozoan and vertebrates, while *Duox A/DuoxMF A* were lost in Ambulacraria and duplicated in mammals, giving rise to the set of genes known in humans (*Duox1*, *Duox2*, *Doxa1*, *Doxa2*) (Figure 3A, B; Figure 4). This has implications for the study of Duox/DuoxMF in non-mammalian models, as it implies that the *Duox A/DuoxMF A* and *Duox B/DuoxMF B* genes present in multiple bilaterian lineages are not respectively orthologous to the mammalian *Duox1/Doxa1* and *Duox2/Doxa2* genes, which have received the most attention so far.

**Figure 3:**
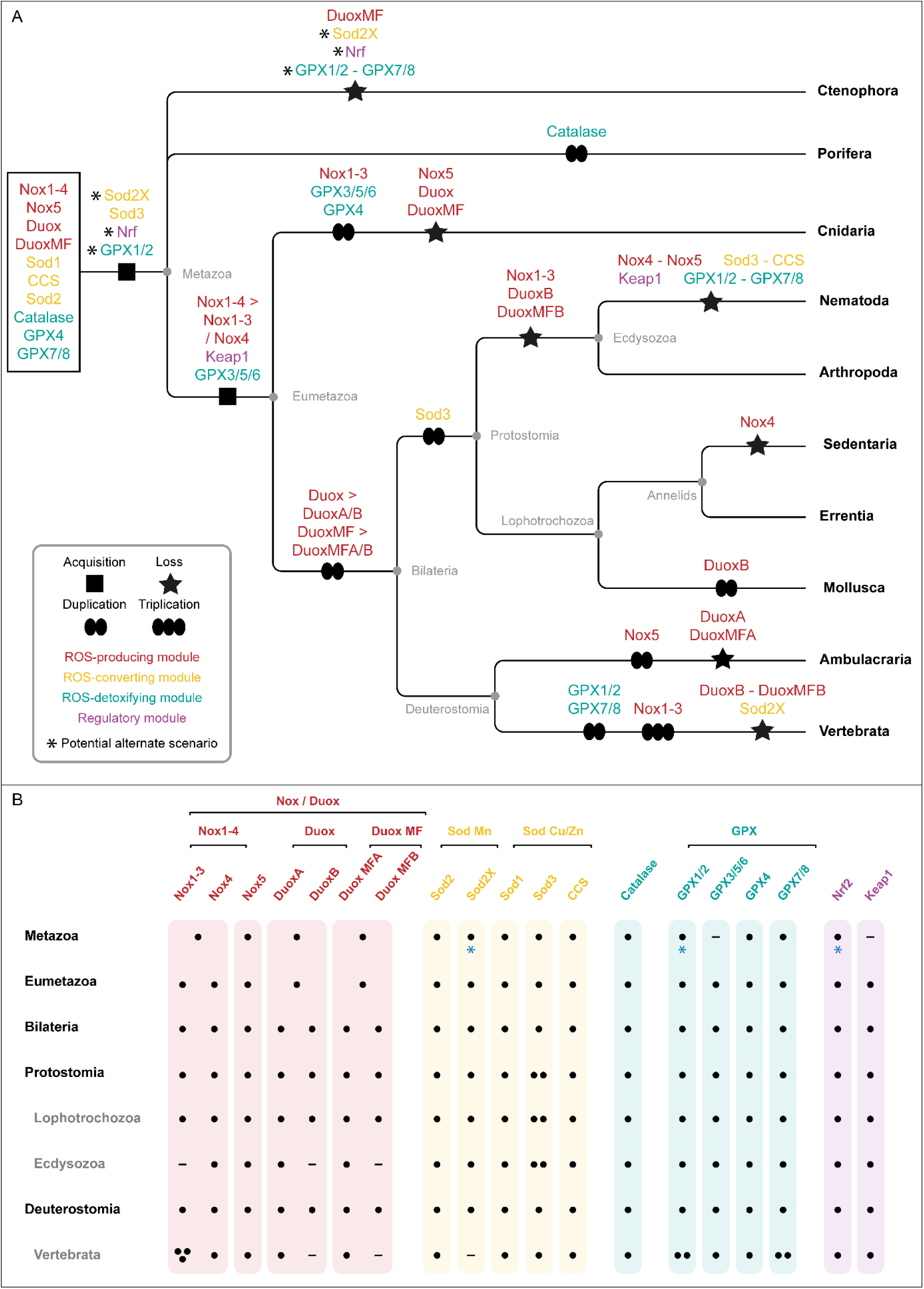
Evolution of ROS metabolism gene families in metazoan. (A) Simplified phylogeny of the major metazoan lineages with the main evolutionary events of the ROS metabolism gene families mapped (acquisition – black square, loss – black star, duplication and triplication – black ovals). Asterisks (*) indicate potential alternative scenario: either the gene was acquired in metazoan and lost in ctenophores (as depicted), or the gene was acquired after ctenophores divergence, in the hypothesis that ctenophores are the sister-lineage to all other metazoan lineages (Schultz et al. 2023). Genes framed on the left preceded the emergence of metazoan. The gene names are colour coded according to the ROS-module to which they belong. (B) Putative ancestral sets of gene families in the main phylogenetic metazoan lineages. The number of black dots indicate the number of genes present in the Last Common Ancestor (LCA) of each lineage, while a dash indicates its absence. A blue asterisk indicates the corresponding gene might be absent in the metazoan LCA if ctenophores are the sister-group of all other metazoan lineages.

**Figure 4:**
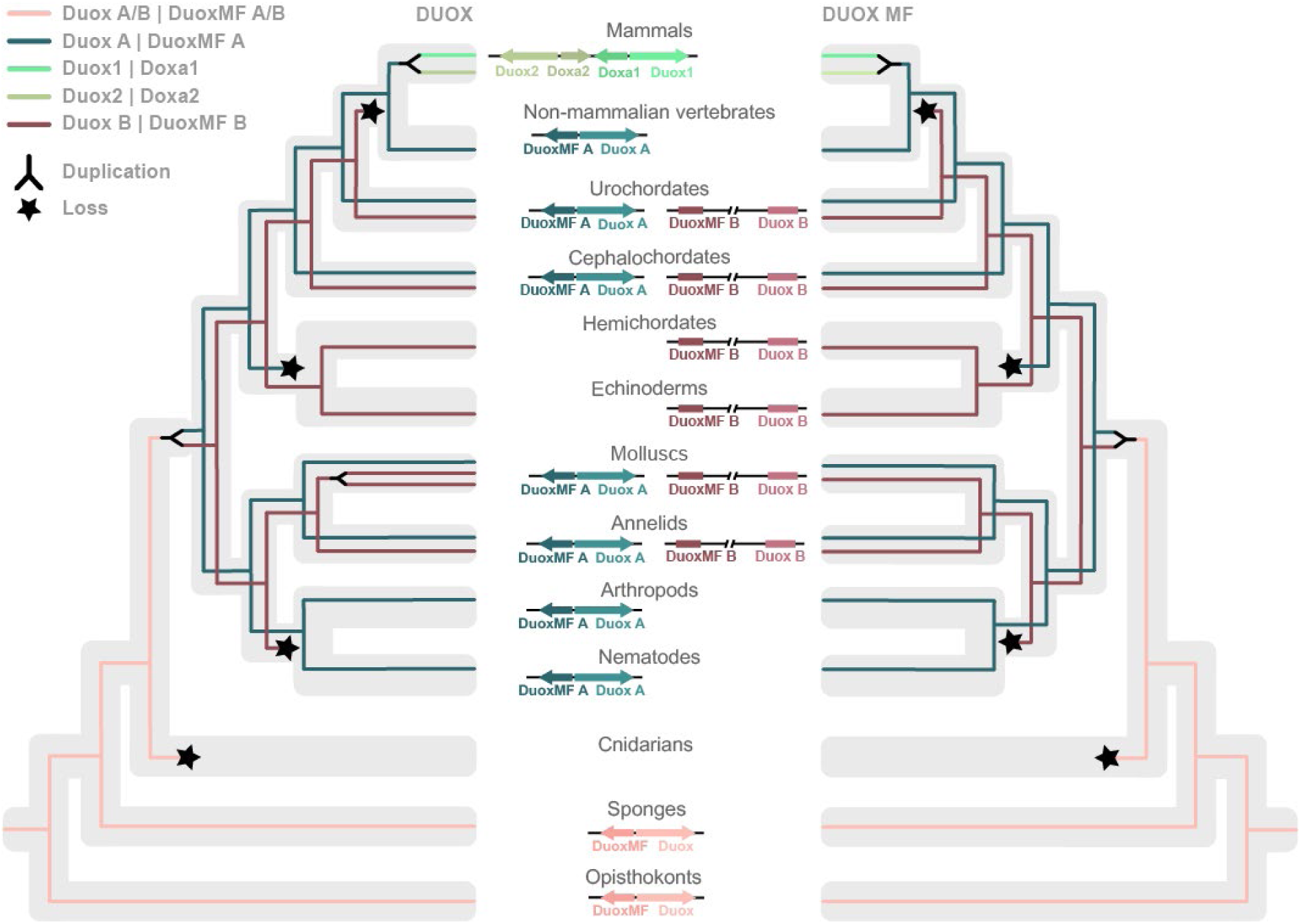
Evolutionary history of the Duox and Duox Maturation Factor families. Schematic representation of the parallel evolutionary scenarios for the Duox and Duox Maturation Factor gene families. The genes trajectories and main duplication (fork) or loss (star) events are mapped over a simplified phylogenetic tree of metazoans (grey background). The schematic in the middle represents the microsynteny of Duox and DuoxMF genes in the major lineages.

Such a congruence of *Duox* and *DuoxMF* evolutionary histories strongly suggests a coevolution of the two gene families. To test this hypothesis, we investigated the potential synteny of *Duox* and *DuoxMF* genes by browsing available genomes, and found that *Duox* and *DuoxMF* genes are arranged head-to-head in sponges and non-metazoans (opisthokonts). This organization is maintained at the *Duox A/DuoxMF A* locus in bilaterian including mammals, in which the additional duplication of *Duox/DuoxMF A* results in a double head-to-head conformation (named Duox 1 and ,2 and Doxa1 and 2, from the mammalian nomenclature). Conversely, this organization is lost for *Duox B/DuoxMF B*, with the two genes being located either far apart or on different chromosomes entirely (Figure 4). Taken together these phylogenetic and syntenic analyses reveal an ancient coevolution of the *Duox* and *DuoxMF* gene families.

#### ROS-converting module: superoxide dismutases

The term “superoxide dismutase” (SOD) refers to enzymes performing the reaction 2O_2_ ^•−^+2H^+^ → H O + O_2_, however it aggregates two groups of proteins with very different domain composition. SOD enzymes that use copper as an electron acceptor (such as Sod1 and Sod3 in humans) are characterized by a Sod Cu/Zn binding domain. This domain is also found in CCS (Copper Chaperone for Sod), a protein required to deliver copper to the catalytic site of human Sod1. Conversely, SOD enzymes that use manganese as an electron acceptor (such as Sod2 in humans) contain two Mn/Fe SOD domains, with no resemblance to the Sod Cu/Zn binding domain. The two families are hypothesized to have acquired their O_2_ ^•−^ dismutation functions convergently (Fink and Scandalios 2002). We thus investigated them separately.

We identified 348 *Sod Cu/Zn* genes (including 70 *CCS*) and 143 *Sod Mn* genes in our dataset (list of protein sequences in Sup File 3 and Sup File 4). A phylogenetic analysis of Sod Cu/Zn sequences reveals three main groups we labelled Sod1, Sod3 and CCS according to human sequences positions in the phylogeny (Sup Fig 3). Domain analyses on all the sequences shows that Sod1 contain solely the Sod Cu/Zn binding domain, while CCS sequences possess an additional Heavy-Metal Associated domain (consistent with their role as copper-delivering chaperones) and Sod3 sequences contain an additional signal peptide targeting them to the endoplasmic reticulum and ultimately to an extracellular location. Species in our dataset possess 1 to 5 copies of Sod1; 1 to 2 copies of CCS; 1 to 7 copies of Sod3 (Figure 2). Gene losses are found in all three families: Sod1 is absent in a lineage of annelids (Spionidae); CCS in the Myxosporea cnidarians, nematodes and several annelid groups; Sod3 in sponges, placozoans, Myxosporea, nematodes, platyhelminths, nemerteans and urochordates (Figure 2).

While the phylogeny of Sod3 does not recapitulates the metazoan phylogeny, the presence of annelid, mollusc and arthropod sequences from similar species at different positions in the tree suggests an ancient duplication of Sod3 in protostomes (Figure 3A). The presence of Sod1 and CCS non-metazoan sequences suggests that the two corresponding genes precede the emergence of metazoans, while Sod3 was acquired in the LCA of metazoans (Figure 3A, B).

A phylogenetic analysis of Sod Mn sequences revealed two clades, labelled Sod2 and Sod2X (nomenclature from Hewitt and Degnan 2022) (Sup Fig 4). Domain analysis shows that in addition to the two Mn/Fe SOD domains, Sod2 possesses a mitochondrial transfer peptide, and Sod2X a signal peptide. In species from our dataset, Sod2 is found in 1 to 2 copies, and Sod2X in 1 to 4 copies (Figure 2). While Sod2 is lost only in Myxosporea, Sod2X is lost in numerous lineages, including ctenophores, placozoans, Myxosporea, multiple ecdysozoan lineages, several annelid ones, cephalopods, hemichordates, urochordates and vertebrates. In addition, Sod2X is duplicated twice in Siboglinidae (annelids) (Fig Sup 4). While Sod2 sequences are found in non-metazoans, Sod2X was acquired in the LCA of metazoans (Figure 3A). Sod2 and CCS evolutionary histories appear rather straightforward, with both genes being found in one copy in most metazoan lineages. On the other hand, the evolution of Sod Cu/Zn and Sod2X genes is more dynamic, with notable variations in the number of Sod Cu/Zn copies (especially for Sod3), and numerous losses of Sod2X in various lineages. This raises the question whether the functional redundancy of Sod Cu/Zn and Sod2X genes (all encoding non-mitochondrial SODs) might have alleviated the evolutionary pressure, allowing for such dynamic history. Conversely, the only mitochondrial SOD-encoding gene and the only SOD chaperone-encoding gene might have been more strictly conserved.

#### ROS-detoxifying module

##### Catalase

We identified 160 *catalase* sequences in our dataset that encode for proteins containing a catalase domain (list of protein sequences in Sup File 5, Sup Fig 5). The number of *catalase* genes *per* species ranges from 1 to 4, with only 1 copy found in most animal species. No catalase gene was found in two lineages within Cnidaria (Alcyonacea and Myxosporea) and one within Annelida (Siboglinidae) (Figure 2). Phylogenetic analysis of catalase sequences does not recapitulate metazoan phylogeny, but allows for the identification of punctual ancestral duplications within Mollusca (the scallops species *Pecten maximus* and *Patinopecten yessoensis*) and Cnidaria (Scleractinia) (Sup Fig 5, black dots). Given the aforementioned results, we infer that an ancestral catalase gene predates the emergence of metazoan species (Figure 3A) and is found in a single copy in the LCA of metazoans, eumetazoans, bilaterians, protostomes and deuterostomes (Figure 3B).

##### The GPX multigenic family

Glutathione peroxidases (GPX or Peroxidases) constitute a multigenic family, of which eight members are known in humans (GPX1-8) (Brigelius-Flohé and Maiorino 2013). We identified 403 *GPX* sequences in our dataset that encode for proteins consisting of a glutathione peroxidase domain (list of protein sequences in Sup File 6). The number of GPX genes *per* species varies a lot, ranging from 1 to 12, and Acoela seems to be the only lineage devoid of any GPX (Figure 2).

Phylogenetic analysis indicates that GXP sequences fall into four main groups that we labelled GPX1/2, GPX3/5/6, GPX 4 and GPX7/8 (similar to Margis et al. 2008, based on the position of human sequences in the phylogenetic tree) (Sup Fig 6). GPX4 and GPX7/8 seem more closely related, as do GPX 1/2 and GPX3/5/6.

These results are consistent with previous studies that identified the same four ancestral groups in vertebrates (Margis et al. 2008) and to a greater extent in metazoans (Trenz et al. 2021). The number of GPX1/2 and GPX7/8 *per* species is rather modest, from 1 to 3 and 1 to 2, respectively. By contrast a more variable number of copies is found for GPX3/5/6 and GPX4, from1 to 6 and 1 to 5 respectively. Mapping the presence/absence and number of copies for all four ancestral groups of sequences in the studied species reveal scattered but frequent events of loss and duplications. GPX1/2 is duplicated in vertebrates and lost in insects, myriapods, nematodes, urochordates and some annelid and cnidarian lineages. GPX3/5/6 is ancestrally duplicated in Cnidaria, in Mollusca, and twice in mammals, but lost in insects and some lophotrochozoan lineages. GPX4 is duplicated in Cnidaria and lost in echinoderms and many annelid lineages. GPX7/8 is duplicated in vertebrates and lost in both insects and nematodes as well as few other scattered lineages. Ancestral state reconstruction suggests that GPX4 and GPX7/8 precede metazoan emergence, while GPX1/2 was acquired in the metazoan LCA, and GPX3/5/6 in the eumetazoan one (Figure 3), however duplications responsible for their respective emergences are not established. All four genes are present in a single copy in the LCA of bilaterians, protostomes and deuterostomes.

#### Antioxidative response module: Nrf and Keap1

The functional association of Nrf2 and Keap1 is well described in vertebrates (Suzuki and Yamamoto 2015). It relies on the presence of two domains in Nrf2, DLG and ETGE, with which the Kelch domains of Keap1 interact, and on three specific Keap1 cysteines (C151, C273 and C288), the oxidation of which causes a change in conformation and the release of Nrf2 (Figure 5A). Nrf1 and Nrf3 are two other transcription factors from the Nrf gene family found in vertebrates. They possess a DNA-binding bZIP domain like Nrf2, and Nrf1 also has the DLG / ETGE motifs. However, they are not reported to participate in the antioxidative response or interact with Keap1.

**Figure 5:**
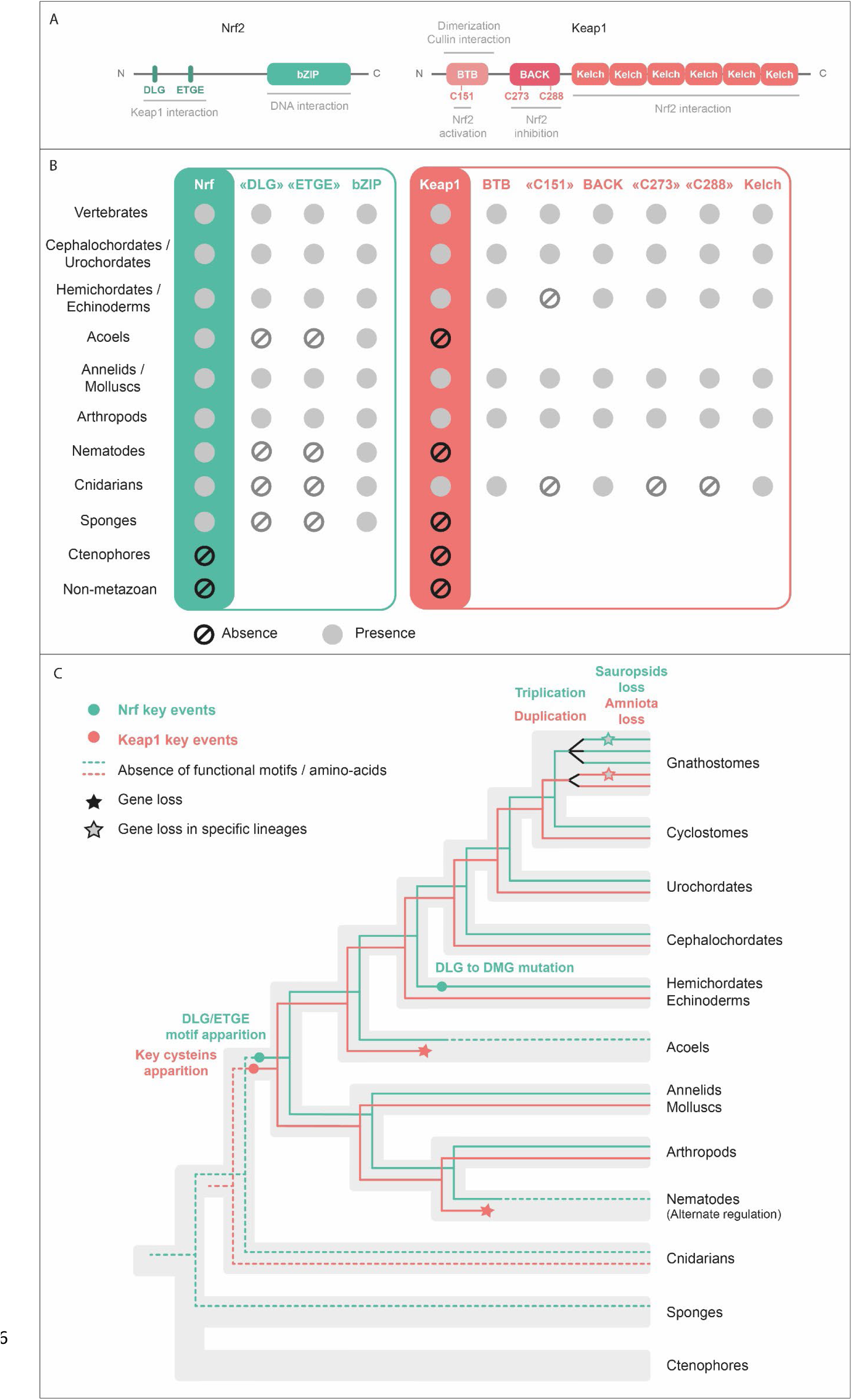
Structure and evolutionary history of Nrf and Keap1 gene families. (A) Schematic representation of Nrf and Keap1 structures. Domains, motifs, key cysteines and their associated functions (as characterized in vertebrates) are indicated. (B) Summary of the presence/absence of domains, motifs and key cysteines of Nrf and Keap1 in the main metazoan lineages. (C) Evolutionary scenario for Nrf and Keap1. Dotted lines are used when DLG/ETGE motifs and key cysteines are absent in Nrf and Keap1 respectively. Major evolutionary events are represented by a star (loss) or a dot (other events).

In our dataset, we identified 101 and 63 protein sequences for NRF and KEAP1 respectively (list of protein sequences in Sup File 7 and Sup File 8). One or two copies of Nrf were found in most species, except for vertebrates that possess up to three copies, and only few metazoan lineages lack Nrf orthologs: ctenophores, Myxosporea (cnidarians) and Spionidae (annelids). Keap1 was also found in a low number of copies *per* species (1 or 2) but is absent in much more metazoan lineages than Nrf: ctenophores, sponges, multiple cnidarian and annelid groups, nematodes and acoels (Figure 2).

A phylogenetic analysis of Keap1 sequences recapitulated the phylogeny of species and revealed ancestral duplications within molluscs (the gastropod species *Biomphalaria glabrata* and *Aplysia californica*) and gnathostomes (Sup Fig 7). A phylogenetic analysis of Nrf sequences, while less consistent with species phylogeny, indicated two successive duplications in gnathostomes, resulting in the Nrf1, 2 and 3 proteins known in humans, of which Nrf3 is lost in sauropsids (Sup Fig 8). Overall, the higher number of Nrf and Keap1 copies found in vertebrates probably results from the well-documented whole-genome duplications in this clade.

Consistent with the fact that those two proteins are supposed to act jointly, we noted that all species lacking a Nrf protein lack Keap1 as well, and that only a subset of cnidarian species possessed a putative Keap1 sequence. These species belong to various cnidarian families and do not constitute a monophyletic group, suggesting numerous scattered losses of Keap1 in cnidarians. To establish whether these cnidarian sequences were indeed Keap1 orthologs, we performed a phylogenetic analysis including our 63 Keap1 sequences as well as a selection of metazoan Kelch-like proteins (Kelp), which domain composition and overall sequences resemble Keap1. Cnidarian sequences of interest group together, and are closer to bilaterian Keap1 sequences (although a little divergent from them) than to any other Kelp group, confirming their orthology (Sup Fig 9).

To assess whether the association of Nrf and Keap1 might be retained in other non-vertebrate lineages, we performed domain analysis on all Nrf and Keap1 sequences and searched for DLG / ETGE motifs and C151, C273 and C288 equivalents (results in Figure 5B, complete analysis in Sup Table 2). From this analysis, we found that most species whose Nrf sequences lack DLG and ETGE motifs (sponges, acoels, nematodes) do not possess a Keap1 sequence. The only exception are cnidarians, in which all species lack DLG and ETGE motifs in Nrf, but some species, as we mentioned earlier, possess a Keap1 sequence lacking the specific cysteines involved in Nrf2-Keap1 dissociation.

To further investigate the possibility of a Nrf / Keap1 interaction in non-bilaterians, we obtained PSOPIA (Prediction Server Of Protein-protein InterActions) scores (Murakami and Mizuguchi 2014) for Nrf / Keap1 interactions based on their sequences, and retrieved high scores for cnidarian Nrf and Keap1 proteins (Sup Fig 10A). However, tests pairing human Nrf2 with another Kelp protein and human Keap1 with Nrf3 also produced high interaction scores, suggesting Kelch-domain containing proteins are susceptible to interaction with members of the Nrf gene family, regardless of DLG / ETGE presence. To look further into the conformation of Nrf / Keap1 association, we used AlphaFold to generate structure predictions for Keap1 dimers and Nrf in several species (Sup Fig 10B). Consistent with the literature, all predictions had the Keap1 proteins dimerizing through their BTB domain, and interacting through their Kelch-repeat domains with two Nrf segments (DLG and ETGE motifs in the case of *H. sapiens* and *Platynereis*, other residues entirely for *M. capitata* and other cnidarian species). This suggests that based on their sequences alone, cnidarian Nrf and Keap1 seem likely to interact. However, whether cnidarian Keap1 proteins can act as regulators of Nrf activity in the absence of the key cysteine residues responsible for their conformation shift in vertebrates remains unknown. Overall it seems that the functional association of Nrf and Keap1 proteins might be conserved in the vast majority of bilaterian lineages.

Taken together, these results suggest that ancestral Nrf and Keap1 genes were acquired in the LCA of metazoans and eumetazoans respectively (Figure 3A). However, these genes lacked the specific motives (Nrf) and amino-acids (Keap1) required for their functional association. These features were acquired for both genes in the LCA of bilaterians (Figure 5B). Other key events of Nrf and Keap1 evolutionary history include the loss of Keap1 in acoels and nematodes (along with Nrf DLG and ETGE motifs), and the mutation of the DLG motif into DMG in ambulacrarians (possibly without consequences given the similar chemical properties of leucine and methionine).

In sum of the genomic assessment of the ROS metabolism machinery, our investigation revealed a dynamic evolutionary history for several genes of the ROS metabolism, with numerous duplications and losses across metazoan lineages. However, most metazoan groups ancestrally possess at least one copy from each gene family of the ROS metabolism arsenal, defining a sort of “core ROS management kit” (1 Nox or Duox, 1 Sod Mn, 1 Sod CuZn and associated chaperone, 1 catalase, 1 GPX, 1 Nrf and 1 Keap1) (Figure 3B).

### ROS involvement in regeneration

Our comparative genomic analyses provided evidence for an evolutionary complex and versatile redox arsenal, involved in ROS production, conversion, and subsequent detoxification during many biological processes such as development, homeostasis and regeneration in animals (Rampon et al. 2018). However, how this arsenal is deployed during these processes, in particular following injury and during the regeneration process remains largely unknown. Indeed, studies report expression variation of ROS metabolism genes in several regenerating models (Han et al. 2014; Wenger et al. 2014; Khan et al. 2017; Carbonell et al. 2021), but rarely explore the spatio-temporal expression dynamic of the entire set of genes over the course of regeneration or their transcriptional control via ROS signalling. In addition, ROS production and its requirement for successful regeneration have been established in axolotl, zebrafish, *Xenopus,* planarian and *Hydra* but remain unknown for a majority of regenerating model systems (Gauron et al. 2013; Love et al. 2013; Pirotte et al. 2015; Al Haj Baddar et al. 2019; Suknovic et al. 2022).

We thus focused on assessing how animals with various degrees of regenerative capacities, possessing different redox arsenal members, respond to an injury signal and re-establish a ROS balance during regeneration.

#### ROS metabolism genes are dynamically expressed during regeneration in metazoans

To get a broad overview of ROS metabolism genes expression dynamics during regeneration, we took advantage of available transcriptomes for a number of highly regenerative species spanning metazoan diversity (2 deuterostomes, 2 protostomes and 2 non-bilaterians). We retrieved temporal expression data for differentially expressed ROS metabolism genes during: tadpole tail regeneration in *Xenopus tropicalis* (0, 6, 15 and 24 hours post-amputation, hpa; non-amputated, NA), caudal fin regeneration in *Danio rerio* (0, 2, 3, 10 days post-amputation, dpa), posterior regeneration in *Platynereis dumerilii* (0, 1, 2, 3, 5dpa), leg regeneration in *Parhyale hawaiensis* (11 timepoints between 0 and 120hpa, NA), apical regeneration in *Hydra vulgaris* (10 timepoints between 0 and 48hpa, NA), oral regeneration in *Nematostella vectensis* (15 timepoints between 0 and 144hpa, NA) (Figure 6; Sup Table 3).

**Figure 6:**
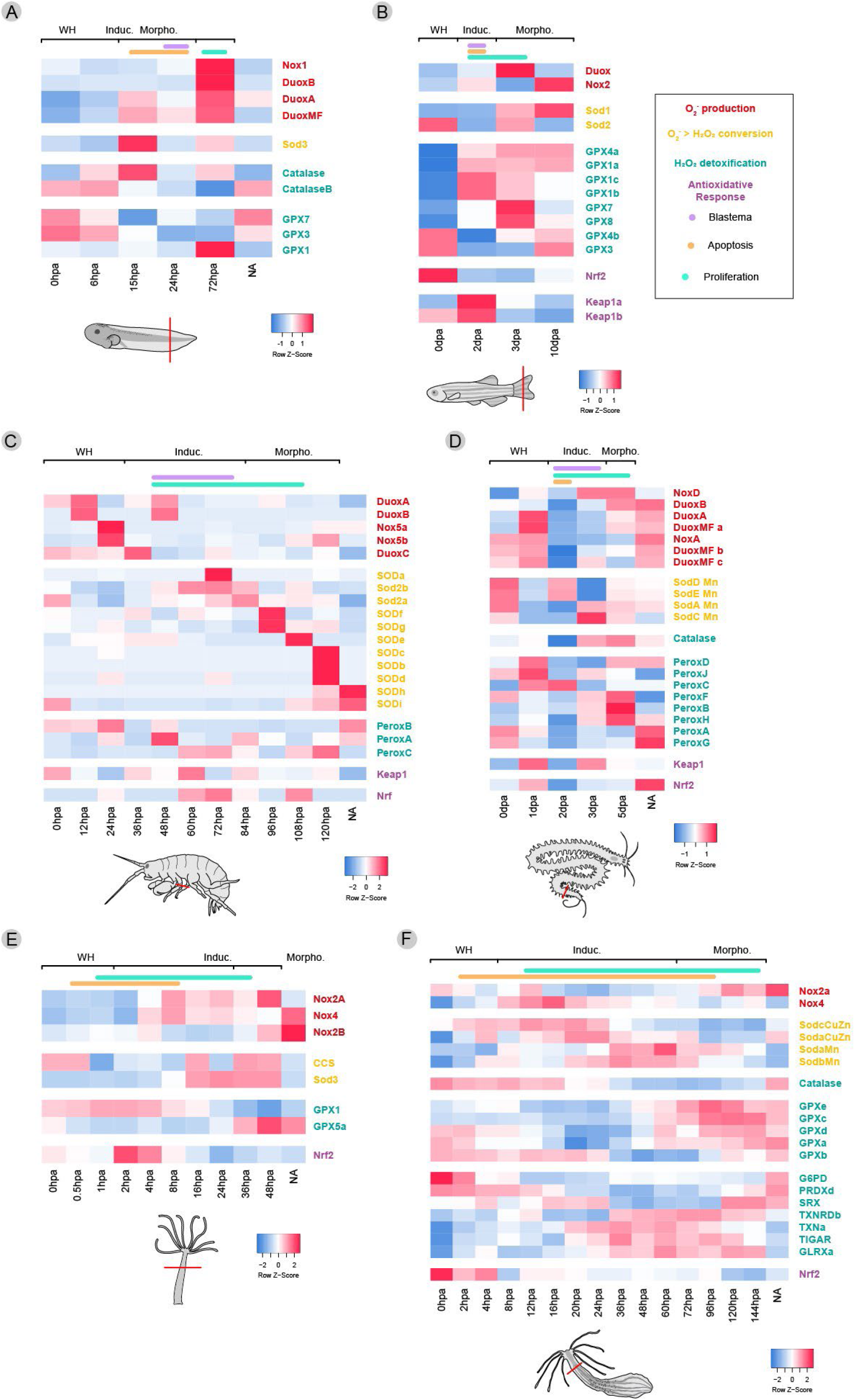
Dynamic expressions of ROS metabolism genes during regeneration in metazoans. *Heatmap representations of the expression levels of differentially expressed genes (Fold Change ≥ 2) of ROS metabolism during (A)* Xenopus tropicalis *tadpole tail regeneration (Chang et al. 2017), (B)* Danio rerio *caudal fin regeneration (Nauroy et al. 2019), (C)* Parhyale hawaiiensis *leg regeneration (Sinigaglia et al. 2022), (D)* Platynereis dumerilii *posterior regeneration (Paré et al. 2023), (E)* Hydra vulgaris *apical regeneration (Wenger et al. 2019) and (F)* Nematostella vectensis *oral regeneration (Warner et al. 2018)*.

Overall, up-regulation of members of the different gene families is found scattered over the course of regeneration, but combinatorial expression patterns emerge for some species. Nox / Duox are upregulated early in *Parhyale* (12-26hpa) and *Nematostella* (8-20hpa), but later in *Hydra* (48hpa), *Xenopus* (72hpa) and *Danio* (3-10dpa). Catalase, GPX and SOD genes expression peaks are scattered, except for the later up-regulation of SOD genes in *Parhyale* (72-120hpa). Finally, Nrf and Keap1 expression peaks are observed either early (*Nematostella*, *Danio*, *Hydra*), late (*Platynereis* Nrf) or at multiple timepoints during regeneration (*Platynereis* Keap1, *Parhyale*).

Altogether, genes of the production, conversion and detoxification modules appear sequentially expressed in a couple of species during regeneration (*Parhyale* and *Nematostella*). In other models, no real trend is observed, suggesting a more complex or post-transcriptional regulation of pro- and anti-oxidant actors is at play. Nevertheless, genes from all three modules are differentially expressed over the course of regeneration in all models, indicating a key role of ROS metabolism regulation during this process. Hence, we propose to extend the data available so far by focusing on two highly regenerative species: the annelid *Platynereis dumerilii* and the cnidarian *Nematostella vectensis*. *Platynereis* possesses a complete ROS arsenal (Sup Table 4) and belongs to the Lophotrochozoa, in which little to no data exist regarding ROS involvement in regeneration. Although this subject has been investigated in another cnidarian, the hydrozoan *Hydra* (Suknovic et al. 2022), *Nematostella* provides an interesting comparison to it, as it belongs to a different cnidarian lineage (Anthozoa) and lacks Duox and Duox MF (as all cnidarians do), but also Keap1 (contrary to *Hydra*) (Sup Table 4).

We performed whole-mount *in situ* hybridization (WMISH) at different stages of *Platynereis* posterior regeneration (0-5 dpa, plus 15dpa as a proxy for non-amputated worms) and *Nematostella* oral regeneration (uncut polyps, 2, 4 and 20hpa), focusing on a set of genes from the different modules of the ROS metabolism (Figure 7; Sup Fig 11). *Nematostella* time points were selected based on the analysis of the temporal expression profile of the genes of interest accessible on the NvERTx database (https://nvertx.ircan.org/ER/ER_plotter/home) (Table Sup 4). Figure 7 shows a selection of expression patterns, while Sup Fig 11 shows the entire panel and associated counts for *Nematostella* expression patterns.

**Figure 7:**
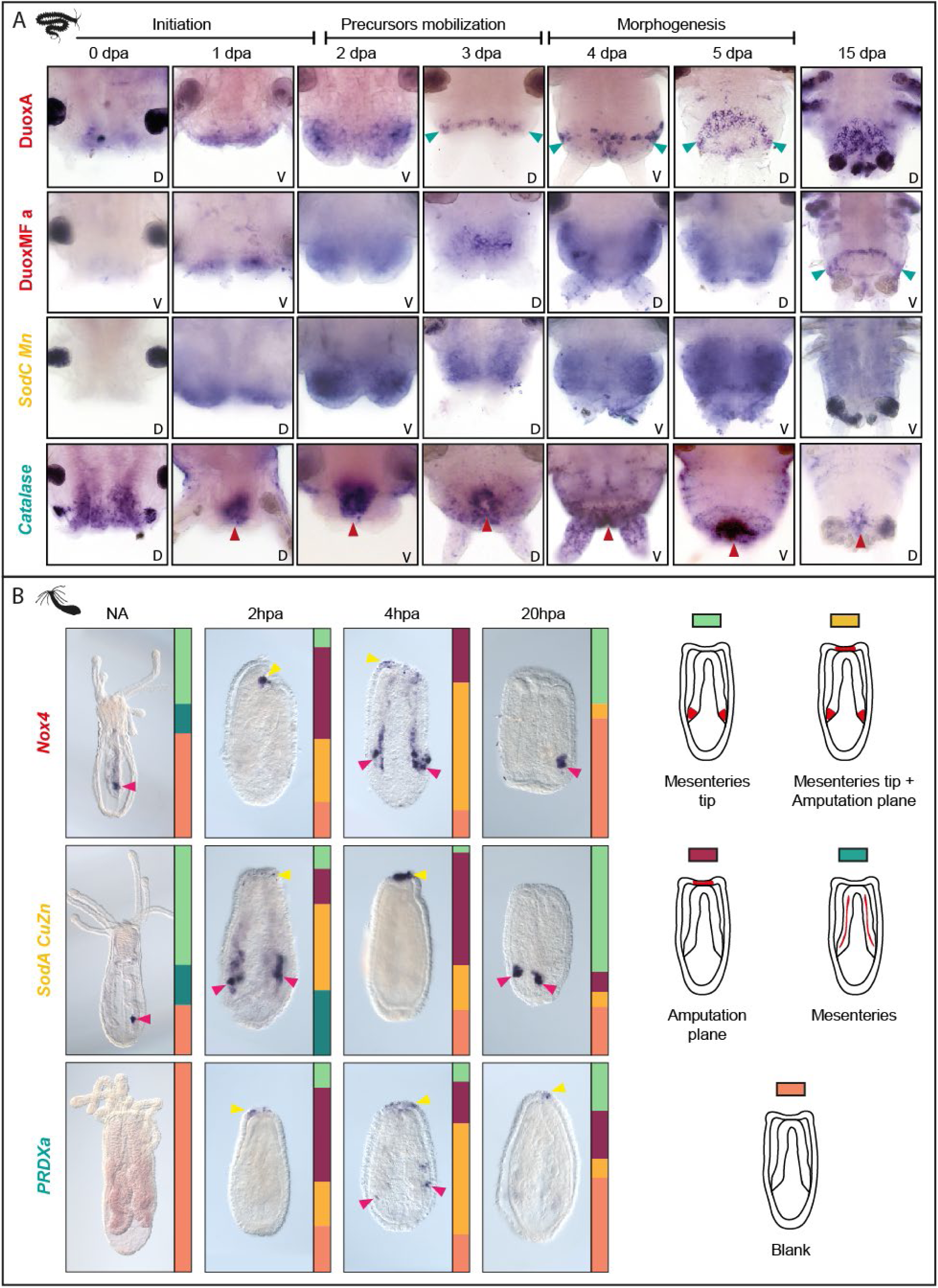
Dynamic spatial expression patterns of ROS metabolism genes during regeneration. *Whole-mount in situ hybridizations of selected genes at different stages of regeneration. (A) Posterior regeneration in* Platynereis. *Blue arrowhead:* DuoxA *and* DuoxMFa *expression in the growth zone. Red arrowhead:* catalase *expression in the anus. V: ventral view. D: dorsal view. Anterior is up. (B) Oral regeneration in* Nematostella. *The main expression pattern is shown. Proportions of alternative patterns (in reference to the cartoons to the right) are indicated below each picture. Yellow arrowhead: expression at the amputation plane. Pink arrowhead: expression in the aboral tips of the mesenteries. Oral is up*.

In *Platynereis*, most genes are expressed close to the amputation plane at 0dpa and 1dpa then more widely in the blastema and regenerated structures at later stages (*SodC Mn*; see other genes on Sup Fig 11), but some display a more specific pattern (Figure 7A). *Catalase* is expressed in the posterior-most part of the gut and the reformed anus, as well as segmentation stripes at 5dpa. *DuoxA* is expressed in the newly formed growth zone as of 3dpa. Interestingly, the gene encoding the associated maturation factor, *DuoxMF a*, is more widely expressed in the blastema during regeneration, but is also expressed in the growth zone at 15dpa. Besides, both genes display similar levels of expression over the course of regeneration (Figure 6D). Altogether, ROS metabolism genes are dynamically and widely expressed in regenerating structures in *Platynereis*.

In *Nematostella* uncut polyps, ROS metabolism genes are either not detected (*PRDXa*), or expressed in the aboral tips of the mesenteries (MTs) in uncut polyps (*Nox4*, *SodA CuZn*) (Figure 7B; Sup Fig 11). Following amputation, *Nox4* is expressed at the amputation plane (AP) at 2hpa, and in both the AP and the MTs at 4hpa. The opposite expression dynamic is observed for *SodA CuZn* and *PRDXa*, with dual expression at the AP and MTs at 2hpa, and a restricted AP expression at 4hpa in most polyps. By 20hpa, expression is lost at the AP, and the major expression pattern observed in non-amputated polyps is restored in most polyps (no expression for *PRDXa*, MTs for *Nox4* and *SodA CuZn*) (Figure 7B). Other genes we investigated display a similar dynamic: they are either not detected or expressed in the mesenteries in intact polyps, then expressed either at the AP or both at the AP and in the MTs following amputation, and the initial expression profile is recovered in most polyps by 20hpa. The expression of *SodB Mn* (ROS conversion) and *GLRXc* (ROS detoxification) are retained longer at the AP (36hpa and 24hpa, respectively) (Sup Fig 11).

These data suggest that most ROS metabolism genes are expressed in the aboral tip of the mesenteries in physiological conditions, and that their expression is induced early at the amputation plane in response to head removal. Contrary to the wide expression patterns observed in *Platynereis*, ROS metabolism genes expression during regeneration in *Nematostella* is more restricted, both spatially (amputation plane and/or aboral mesenteries tips) and temporally, with most regeneration-specific gene expression detectable within the first 24 hours following amputation.

#### Dynamic ROS production during Platynereis and Nematostella regeneration

We then assessed whether, and if present where and when, ROS production takes place during posterior regeneration in *Platynereis*, thanks to the use of ROS-sensitive dye CellROX Green that labels ROS-producing cells. ROS production was detected as early as 1hpa and disappeared by 72hpa (Figure 8A). Quantification shows that the proportion of CellROX+ cells increases shortly after amputation, going from nearly 0% in non-amputated individuals to 9% at 1hpa, and reaching a peak at 24hpa with 25% of ROS-producing cells. It then significantly decreases (8,3% at 36hpa and 4% at 48hpa) and returns to basal levels at 72hpa (Figure 8B; Sup Fig 12 for detailed counts). ROS-producing cells were localized close to the amputation plane, in the three outermost cell layers (Figure 8A, sagittal sections). Double labelling for ROS-producing cells (CellROX) and cell proliferation (EdU) revealed no colocalization and indicates that most of this ROS production precedes cell proliferation burst (Sup Fig 13), which takes place at 48hpa (Planques et al. 2019). Thus, the two processes appear to be activated sequentially.

**Figure 8:**
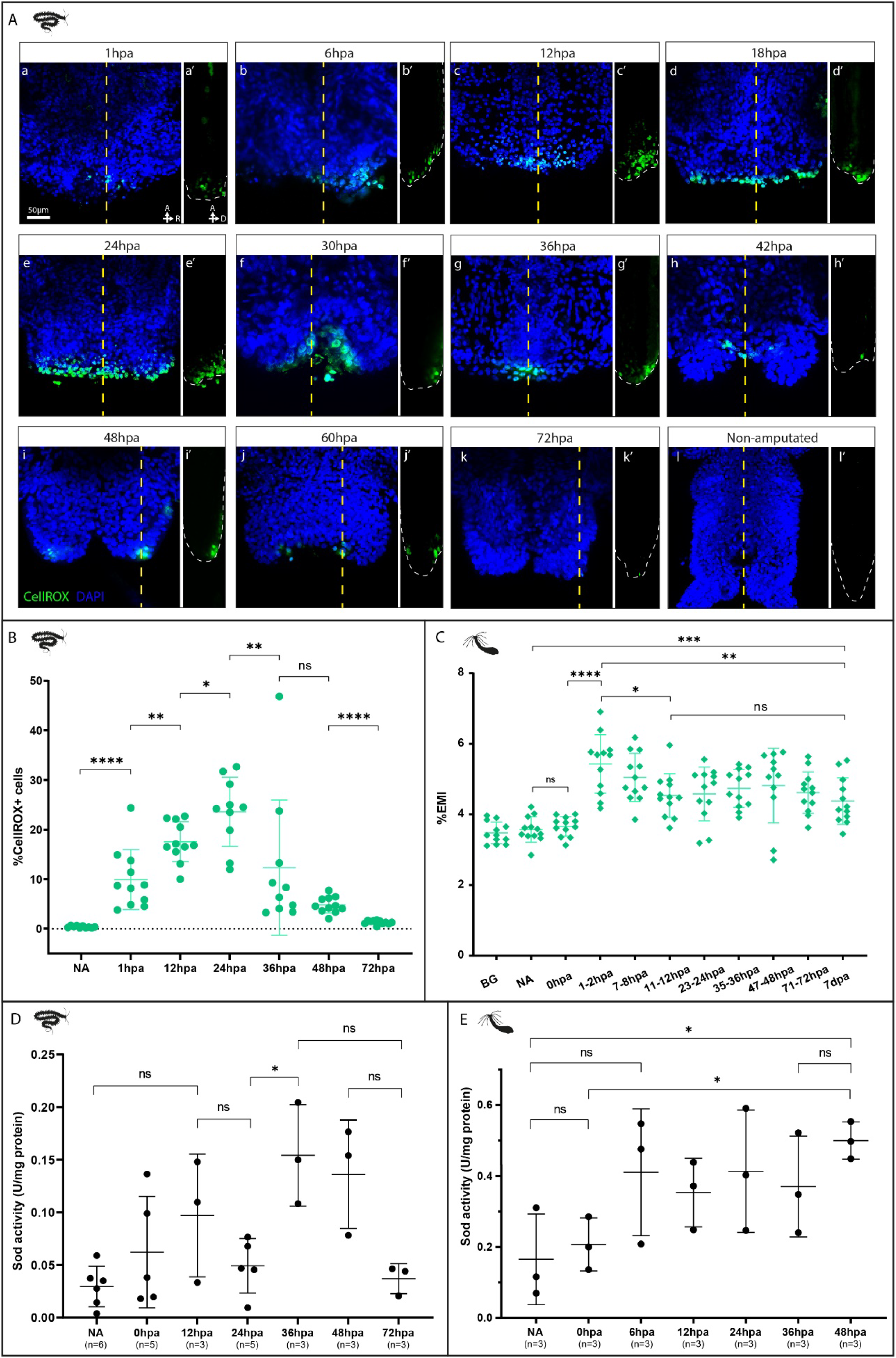
Characterisation of ROS production during regeneration in Platynereis and Nematostella. (Aa-Al) Confocal Z-stacks projections showing ROS-producing cells (nuclei-located CellROX labelling in green; nuclei labelling in blue) in the regenerating posterior part of Platynereis *from 0 to 72 hours post-amputation (hpa, Aa-Ak) and in non-amputated (NA) control (Al). Ventral views, anterior is up. Virtual sagittal sections along the dotted line are shown right of each stack (Aa’-Al’). (B) Quantification of the proportion of CellROX positive cells in the regenerating posterior part (7 timepoints, n=10). Each bracket corresponds to a Mann-Whitney U test (ns p>0.05; * p≤0.05; ** p≤0.01; **** p≤ 0.0001). Mean + SD are shown. For count details see Sup* Fig 12. *(C) Fluorescence readout of Amplex UltraRed for H2O2 production in non-amputated (NA) or in regenerating (hpa)* Nematostella *polyps over 1h, and immediately following amputation (0hpa) (n=12 for each timepoint). BG: background value in 1/3ASW. Each bracket corresponds to a Mann-Whitney U test (ns p>0.05; * p≤0.05; ** p≤0.01; *** p≤0.001; **** p≤ 0.0001). Mean + SD are shown. (D-E) SOD enzymatic activity detected in* Platynereis *(D) and* Nematostella *(E) during regeneration and in non-amputated controls, showing at least three replicates* per *time-point. Each bracket corresponds to a Mann-Whitney U test (ns p>0.05; * p≤0.05). Mean + SD are shown*.

In *Nematostella*, we used the Amplex UltraRed reagent to establish whether and when H_2_O_2_ production takes place following amputation (Figure 8C). A basal level of H_2_O_2_ was observed in non-amputated (NA) and 0hpa polyps, when compared to background (BG) levels. By 1hpa, H_2_O_2_ levels significantly increase (with reporter fluorescence rising by 25% compared to non-amputated individuals), before decreasing at 12hpa. Lower levels of H_2_O_2_ production then persisted over throughout regeneration and were still measured by 7dpa (Figure 8C).

As CellROX does not discriminate between different ROS types produced, we studied the activity of SOD enzymes in regenerating parts to assess whether the burst of ROS in *Platynereis* correlates with an increase of H_2_O_2_ production. SOD activity did not significantly vary in the early stages of regeneration, but increased at 36hpa, namely after the ROS burst, suggesting that most of the observed labelling might not be due to H_2_O_2_ (Figure 8D). In *Nematostella*, SOD activity appeared to increase upon amputation, though not significantly until rather late (48 hpa) (Figure 8E), suggesting that other potential sources might be involved in the early increase of H_2_O_2_ production, such as Nox4 which has been proposed to generate it directly (without the involvement of SODs) (Nisimoto et al. 2014).

To gain additional insights about the activity of enzymes that may be responsible for the decrease of ROS production, we measured catalase enzymatic activity for both animals. We found that catalase activity does not vary significantly over the course of regeneration (Figure 9A) in *Platynereis*, while it increases by 48hpa in *Nematostella* (Figure 9B). As *PeroxA* and *PeroxC* are more widely expressed than *catalase* in *Platynereis* over the course of regeneration (Sup Fig 11), we measured glutathione peroxidase (GPX) activity in the regenerating posterior part of *Platynereis* to establish if GPX could be responsible for the decrease of ROS production. We find that GPX activity is increased at 48hpa, coincident with ROS dropping (Figure 9C). Hence, it appears that catalase activity might underlie ROS detoxification during regeneration in *Nematostella*, while GPX, not catalase, are responsible for ROS decrease in *Platynereis*.

**Figure 9:**
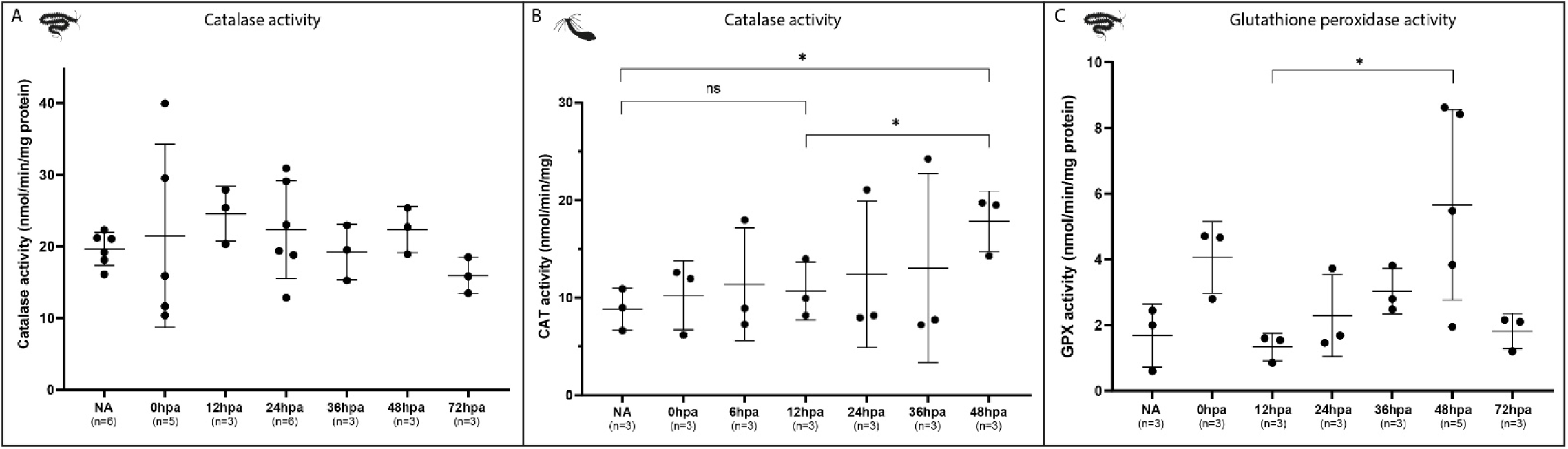
*Enzymatic activity of ROS detoxification actors during* Platynereis *and* Nematostella *regeneration*. Catalase (A: Platynereis*; B:* Nematostella*) and GPX (glutathione peroxidase) (C:* Platynereis*) enzymatic activities detected during regeneration and in non-amputated controls, showing at least three replicates* per *time-point. Each bracket corresponds to a Mann-Whitney U test (ns p>0.05; * p≤0.05). Mean + SD are shown*.

In sum, these analyses show that upon *Platynereis* posterior amputation, a production of non-H_2_O_2_ ROS is very rapidly triggered in superficial cells at the amputation plane, reaches its maximum when the wound epithelium is completely reformed and returns to basal level thanks to the action of GPX. During oral regeneration of *Nematostella*, a burst of H_2_O_2_ occurs in the early hours after bisection, initially in a SOD-independent manner but then relayed by SOD activity at later time points. In contrast to *Platynereis*, ROS detoxification in *Nematostella* occurs *via* catalase activity. The three ROS metabolism modules (production, conversion, detoxification) seem to be sequentially activated during regeneration in both species.

#### ROS production inhibition impairs regeneration success in *Platynereis* and *Nematostella*

##### Morphological effects of ROS production inhibition

To assess the role of ROS during regeneration, we abolished their production using two widely used pharmacological inhibitors: apocynin (APO), an inhibitor of Nox/Duox assembly (Stolk et al. 1994), and diphenylene iodonium (DPI), a general flavoprotein (electron transporters) inhibitor (O’Donnell et al. 1993). Concentration assays determined that continuous treatment over the course of regeneration with APO 500µM or DPI 10nM efficiently blocked regeneration at stage 2 without causing lethality or worm autotomy (indicative of an important stress) in *Platynereis* (Sup Fig 14A, B). In *Nematostella*, continuous treatment with APO (100µM and 500µM) only affected regeneration in a minority of individuals (Sup Fig 14C), while with DPI 100nM regeneration was efficiently blocked at an early stage (stage 1, Amiel et al, 2015) without causing tissue damages or lethality (Sup Fig 14D). We decided to pursue our experiments using APO in *Platynereis*, as it is supposedly more specific than DPI (Altenhöfer et al. 2015), and to this end we confirmed that APO treatment did significantly decrease ROS production using CellROX labelling and quantification on treated animals (from 22% to 4% CellROX+ cells at 24hpa, Sup Fig 14E-F). Conversely, since the effects of APO treatment were not as consistent in *Nematostella*, we pursued our experiments using DPI, which has been used successfully to inhibit bisection-induced H_2_O_2_ production in another cnidarian, *Hydra* (Suknovic et al, 2022).

We then performed three independent inhibition treatments in *Platynereis* for 5 days (duration of regenerative process) with daily scoring of control (DMSO 0,5%) *versus* treated (APO 500µM) individuals, and confirmed that in APO-treated animals regeneration is significantly delayed as early as 2dpa and then stalled, with very few individuals that regenerate beyond stage 2 (Figure 10A). A two-lobed blastema was observed in 5dpa APO-treated subjects that presented no noticeable morphological difference with 2dpa control ones (Figure 10B). In *Nematostella*, inhibition treatments were similarly conducted on control (DMSO 1%) *versus* treated (DPI 100nM) individuals. Scoring at 7dpa (end of regenerative process) confirmed that the majority of treated individuals were significantly blocked at stage 1 (Figure 10C), *i.e.* they underwent wound-healing, and the oral tips of their mesenteries are fused together and in contact with the epithelia at the amputation plane (Figure 10D). F-actin staining by phalloidin further indicated that DPI treatments in *Nematostella* do not affect muscle organization in the mesenteries and body wall epithelia (Figure 10D).

**Figure 10:**
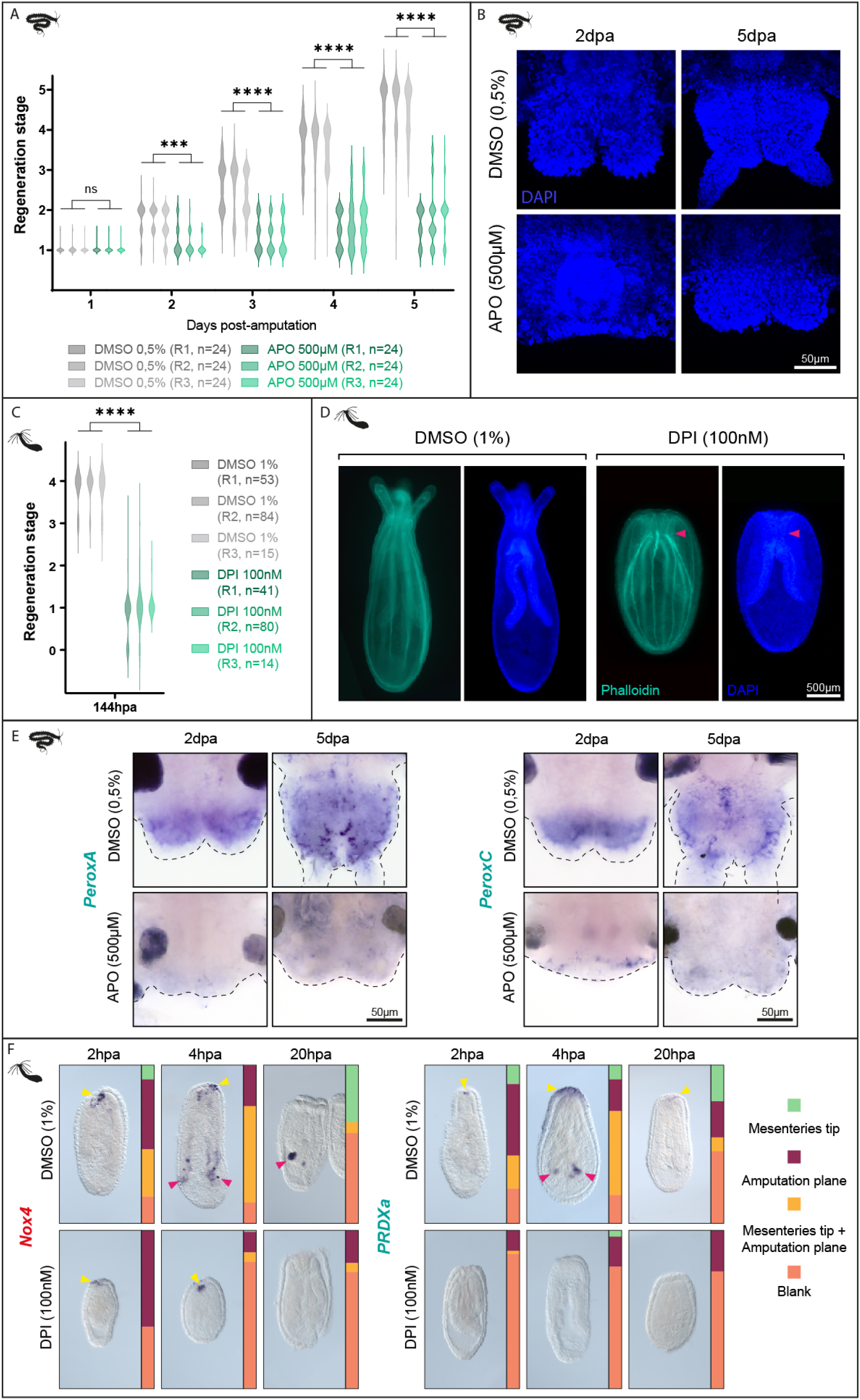
Morphological and transcriptional effects of ROS production inhibition. (A) Daily scoring for the regeneration stage reached by Platynereis *worms during 5dpa (days post-amputation), under APO 500µM treatment versus DMSO 0,5% controls. Three replicates (R1, R2, R3) were performed independently with 24 worms for each condition. Each bracket corresponds to multiple Mann-Whitney tests (ns p>0.05; *** p≤0.001; **** p≤0.0001). (B) Morphology of* Platynereis *regenerating posterior parts at 2 and 5dpa under APO 500µM treatment versus DMSO 0.5% controls, shown with nuclei DAPI labelling. (C) Regeneration stage reached by* Nematostella *polyps at 7dpa, under DPI 100nM treatment versus DMSO 0,1% controls. Three replicates (R1, R2, R3) were performed independently with 14 to 84 polyps* per *condition. Each bracket corresponds to Mann-Whitney test (**** p≤0.0001). (D) Morphology of regenerating* Nematostella *polyps at 7dpa under DPI 100nM treatment versus DMSO 0.1% controls, shown with actin labelling (phalloidin) and nuclei labelling (DAPI). Red arrowhead: contact between the mesenteries and the amputation plane. (E) Whole mount in situ hybridization for two putative target genes of the antioxidative response transcription factor Nrf, peroxA and peroxC, at 2 and 5dpa under APO 500µM treatment and DMSO 0.5% control in* Platynereis. *Dorsal views, anterior is up. (F) Whole mount in situ hybridization for a putative target gene of the antioxidative response transcription factor Nrf, PRDXa, and a potential ROS producer during regeneration, Nox4, at 2, 4 and 20hpa under DPI 100nM and DMSO 1% control in* Nematostella. *Oral is oriented toward the top*.

We evaluated the effect of inhibiting ROS production on putative target genes of the antioxidative response transcription factor Nrf in both species, as well as *Nox4* (not a putative target of Nrf, but a potential ROS producer during regeneration) in *Nematostella*. In *Platynereis*, *PeroxA* and *PeroxC*, were widely expressed in the blastema and regenerated structures (Sup Fig 11). This expression disappeared under APO treatment at 2 and 5dpa (Figure 10E). In *Nematostella*, *PRDX4a* and *Nox4* were expressed in the amputation plane and/or the mesenteries tip at 2, 4 and 20hpa. This expression was lost in most polyps under DPI treatment at all stages (proportion of ‘blank’ phenotype increased) (Figure 10F). This strongly suggests that expression of these putative Nrf target genes is triggered by ROS production during regeneration in both species.

Thus, ROS production is crucial for regeneration success in *Platynereis* and *Nematostella*, and the transcriptional antioxidant response possibly relies on Nrf in a way similar to vertebrates.

##### ROS production inhibition differently affects apoptosis and proliferation

Inhibition of ROS production impaired regeneration and affected antioxidative response-associated gene expression, demonstrating the requirement of ROS for successful regeneration. As in a number of regeneration models (axolotl, *Drosophila*, *Hydra*…) ROS production is responsible for apoptosis induction after injury (Fogarty et al. 2016; Carbonell et al. 2021; Tursch et al. 2022), we investigated the cell death profile following ROS inhibition using TUNEL assay.

We found that although their morphologies differed, apoptosis kinetics appeared similar in control and APO-treated *Platynereis* during regeneration (Figure 11A). Indeed, quantification of TUNEL+ cells revealed no significant difference at 2 dpa, and a higher proportion of apoptotic cells in 5dpa treated worms than in control ones, possibly due to the fact that cells remain in a 2dpa-like apoptotic state, where cell death is important (Figure 11B). In *Nematostella*, while DPI treatment did not affect the early wave of apoptosis close to the amputation plane at 2hpa (Johnston et al, 2021), it drastically reduced the second wave at 24hpa, when dying cells are more widely distributed in the body wall epithelia of control polyps (Figure 11C). Quantification of TUNEL+ cells confirmed these observations, with a similar number of apoptotic cells found in both conditions at 2hpa, but decreased five-fold at 24hpa in DPI-treated polyps (Figure 11D).

**Figure 11:**
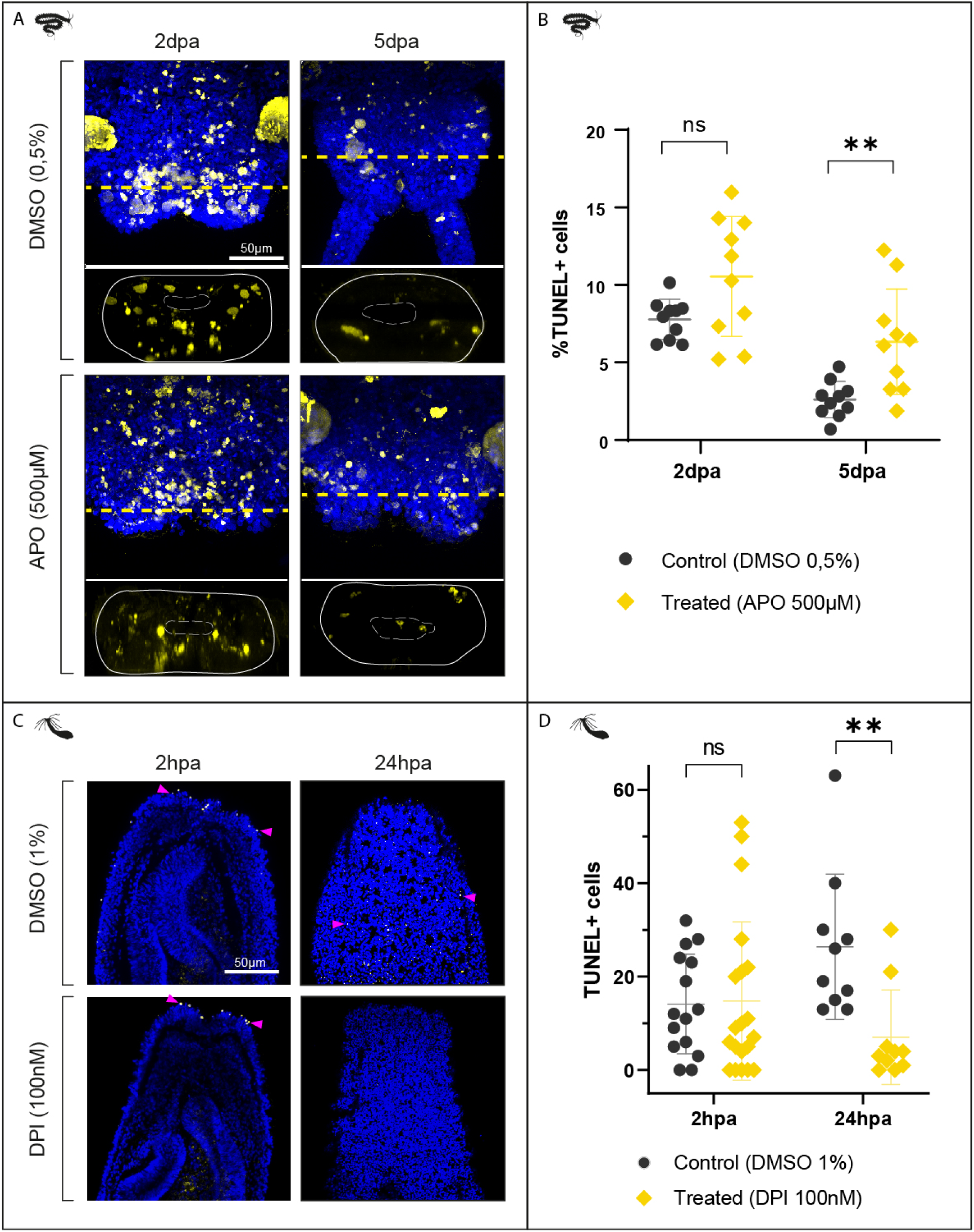
Effects of ROS inhibition on apoptosis during Platynereis and Nematostella regeneration. (A) Cell death labelling (TUNEL assay, yellow) at 1, 2, 3 and 5dpa under APO 500µM treatment and DMSO 0.5% control in Platynereis. *Ventral views, anterior is up. (B) Proportion of TUNEL positive cells in the regenerating posterior part at 2 and 5dpa under APO 500µM treatment and DMSO 0.5% control. Each bracket corresponds to a Mann-Whitney U test (ns p >0.05; ** p≤0.01). (C) Cell death labelling (TUNEL assay, yellow) at 2 and 24hpa under DPI 100nM and DMSO 0.1% control in* Nematostella. *Pink arrowheads: TUNEL+ cells. Oral pole is up. (D) Amount of TUNEL+ cells in the oral part at 2 and proportion of TUNEL+ cells at 24hpa under DPI 100nM and DMSO 0.1% control. Each bracket corresponds to a Mann-Whitney U test (ns p >0.05; ** p≤0.01)*.

A mitogenic role of ROS is established during regeneration in various models such as *Xenopus*, zebrafish, axolotl and planarian (Gauron et al. 2013; Love et al. 2013; Al Haj Baddar et al. 2019; Van Huizen et al. 2019). We therefore tested if ROS is required for cell proliferation during regeneration in *Platynereis* and *Nematostella* (Figure 12).

**Figure 12:**
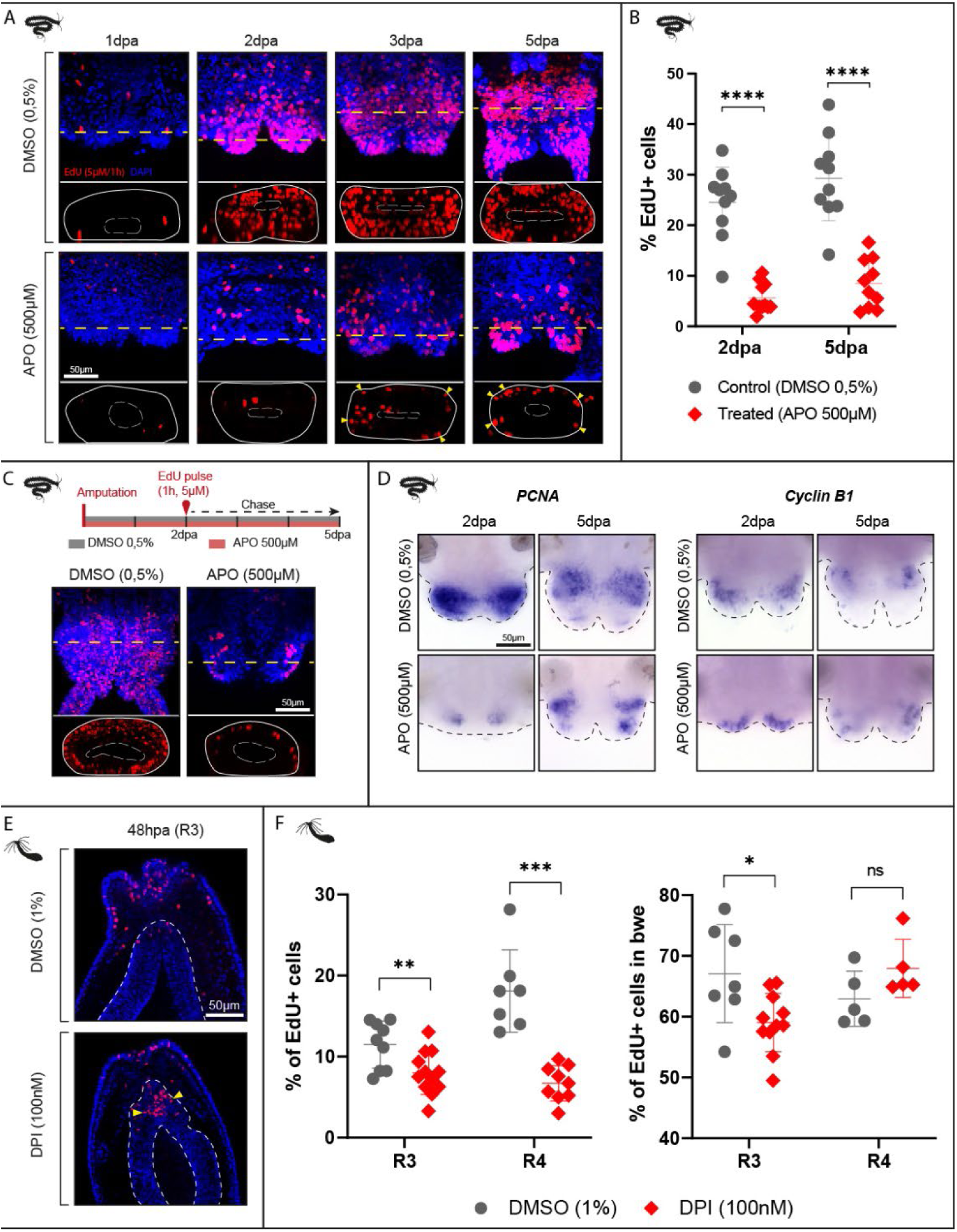
Effects of ROS inhibition on cell proliferation during Platynereis and Nematostella regeneration. (A) EdU labelling (red) at 1, 2, 3 and 5dpa under APO 500µM treatment and DMSO 0.5% control in Platynereis. *Virtual transversal sections along the dotted line are shown below. Ventral views, anterior is up. Yellow arrowhead: ectodermal proliferation. (B) Quantification of EdU positive cells in the regenerating posterior parts at 2 and 5dpa under APO 500µM treatment and DMSO 0.5% control. Each bracket corresponds to a Mann-Whitney U test (**** p≤0.0001). Mean + SD are shown. Ventral views, anterior is up. (C) Schematic for the EdU pulse and chase experiment under APO treatment or control conditions (DMSO 0,5%), and associated confocal Z-stack projections at 5dpa in* Platynereis. *(D) Whole mount in situ hybridization for two cell cycle genes,* pcna *and* cyclinB1*, at 2 and 5dpa under APO 500µM treatment and DMSO 0.5% control in* Platynereis. *Ventral views, anterior is up. (E) EdU labelling (red) at 48hpa under DPI 100nM treatment and DMSO 0.1% control in* Nematostella *(R3). Oral pole is up. (F) Proportion of EdU positive cells in the regenerating oral part (left) and proportion of EdU+ cells found in the body wall epithelium (bwe) (right) at 48hpa under DPI 100nM treatment and DMSO 0.1% control (R3 and R4). Each bracket corresponds to a Mann-Whitney U test (ns p >0.05; * p≤0.05; ** p≤0.01; *** p≤0.001)*.

In *Platynereis*, we observed that APO-treated worms were able to initiate a bi-lobated blastema (stage 2), a process that is proliferation-independent, but did not progress into the further proliferation-dependent stages of posterior regeneration (Planques et al. 2019). To evaluate the effect of ROS production inhibition on cell proliferation, we labelled S-phase cells with a 1h EdU incorporation pulse in DMSO control and APO-treated *Platynereis*. We observed that proliferation was drastically reduced in APO-treated worms compared to controls at all investigated stages of regeneration (Figure 12A), confirmed by quantification of the proportion of EdU+ cells at 2 and 5dpa (decreasing from 25% to 5% and 31% to 9% respectively) (Figure 12B). In addition, most remaining EdU-labelled cells were found localized essentially in outer cell layers at 5dpa in APO-treated worms (Figure 12A, virtual transversal sections, yellow arrowheads).

To assess if EdU-incorporating cells in APO-treated subjects progress into the cell cycle and divide, we performed an EdU pulse-and-chase experiment by incubating APO-treated and control worms 1h in EdU at 2dpa and chasing for 3 days before observing them. In 5dpa control worms, as expected, we observed stippled patterns of EdU labelling, due to its dilution over successive cell divisions (Figure 12C). In APO-treated worms however, EdU+ nuclei were rather uniformly labelled, indicating that not only fewer cells are incorporating EdU, but most of them are stuck in S-phase and do not divide. This was confirmed by the expression patterns of two previously characterized cell cycle genes, *pcna* and *cycB1* (Demilly et al. 2013; Gazave et al. 2013), both being widely expressed in the blastema of control worms at 2 and 5dpa, while their expression territory is reduced in APO-treated 5dpa worms (even if the structure is itself reduced) (Figure 12D). The remaining labelling appeared restricted to the blastema periphery, consistent with the location of EdU+ cells in 5dpa APO-treated worms (Figure 12A).

Similar labelling of S-phase cells using EdU was performed in DMSO control and DPI-treated *Nematostella* polyps in “early” (R1 & R2, Sup Fig 15) and “late” 48hpa polyps (R3 & R4, Figure 12 E,F), defined by the EdU / DAPI ratio (higher in “late” vs “early” - Sup Fig 15). We observed that in “late” 48hpa (R3 + R4) polyps the overall proliferation significantly decreased between DMSO and DPI-treated animals (Fig 12F), mainly driven by a stronger reduction of cell proliferation in the body-wall epithelia (*bwe*) than in the mesenteries (*mes*) as observed in one of the replicates (R3, Figure 12 E, F). A similar significant decrease of cell proliferation in the *bwe* compared to the *mes* was also observed in one of the “early” replicates (R2), while overall proliferation remained unaffected in “early” 48hpa (R1 + R2) polyps (Sup Fig 15). Interestingly, by assessing the EdU/DAPI ratios in DMSO *vs* DPI (R1 + R2 + R3 + R4), it seems that cell proliferation is globally stalled in “early” 48hpa stage state (Sup Fig 15). Taken together these data suggest a role of ROS in pursuing the burst of cell proliferation at 48hpa and distinct effects of ROS in the mesenteries *versus* epithelium of the body wall at the amputation plane during regeneration in *Nematostella*.

Our data show that ROS inhibition during regeneration does not affect apoptosis in *Platynereis*, while in *Nematostella* it inhibits the widespread apoptotic wave taking place at 24hpa, but does not affect the early apoptotic events at the amputation plane at 2hpa. As for proliferation following ROS inhibition, it is drastically reduced in *Platynereis* internal tissues, while it is globally affected in *Nematostella* with a predominant reduction of EdU positive cells in the epithelium of the body wall.

##### Distinct effects of the inhibition of ROS production in *Platynereis* ectodermal and meso-endodermal tissues

We then wondered whether regeneration was stalled strictly by a default of proliferation or if other defects were involved in its failure. To answer this question, we further characterized the resulting phenotype of APO-treated *Platynereis* worms using molecular markers involved in tissue patterning and differentiation.

We first assessed the expression pattern of genes associated with the growth zone (*Hox3*, Sup Fig 16A1), stem cells and progenitors (*PiwiB*, Sup Fig 16A2), and segmentation (*Engrailed*, Sup Fig 16A3). For all three genes, the expression pattern observed in 5dpa APO-treated worms was similar to the one observed in 2dpa control worms. This result indicates that a growth zone is reformed but not very active, as very few tissues are produced and segmentation is not initiated. *PiwiB* expression seemed more restricted under APO treatment, which might be caused by reduced cell proliferation (Figure 12A, B). We next focused on meso-endodermal tissues specification and organisation. Phalloidin labelling (actin staining) revealed that the pygidial muscle ring was formed in 5dpa APO-treated worms, suggesting *de novo* formation of muscle fibres during regeneration is not dependent on ROS production (Sup Fig 16B1). Markers of mesodermal derivatives (*Twist*, Sup Fig 16B2) and the gut (*FoxA*, Sup Fig 16B3) displayed similar patterns in 5dpa APO-treated worms and 2dpa controls, indicating that the specification of the mesoderm and gut was initiated but stalled. Overall, it seems neither global morphology nor meso-endodermal specification can progress beyond a 2dpa-like stage under inhibition of ROS production.

We then investigated genes known to be expressed in the ectoderm at 2dpa: *Wnt1* (expressed in the wound epithelium Figure 13A), *Caudal* (expressed in ectodermal cells and the gut Figure 13B), *Myc* and *Pl10* (expressed largely in the blastema including the superficial cells, Figure 13C, D). Ectodermal expression of all four markers (red arrowheads in control worms) was either lost or reduced in 5dpa APO-treated worms. *Caudal* is the most striking example, its expression being retained in the gut but lost in the ectodermal cells of the regenerating parts (Figure 13B).

**Figure 13:**
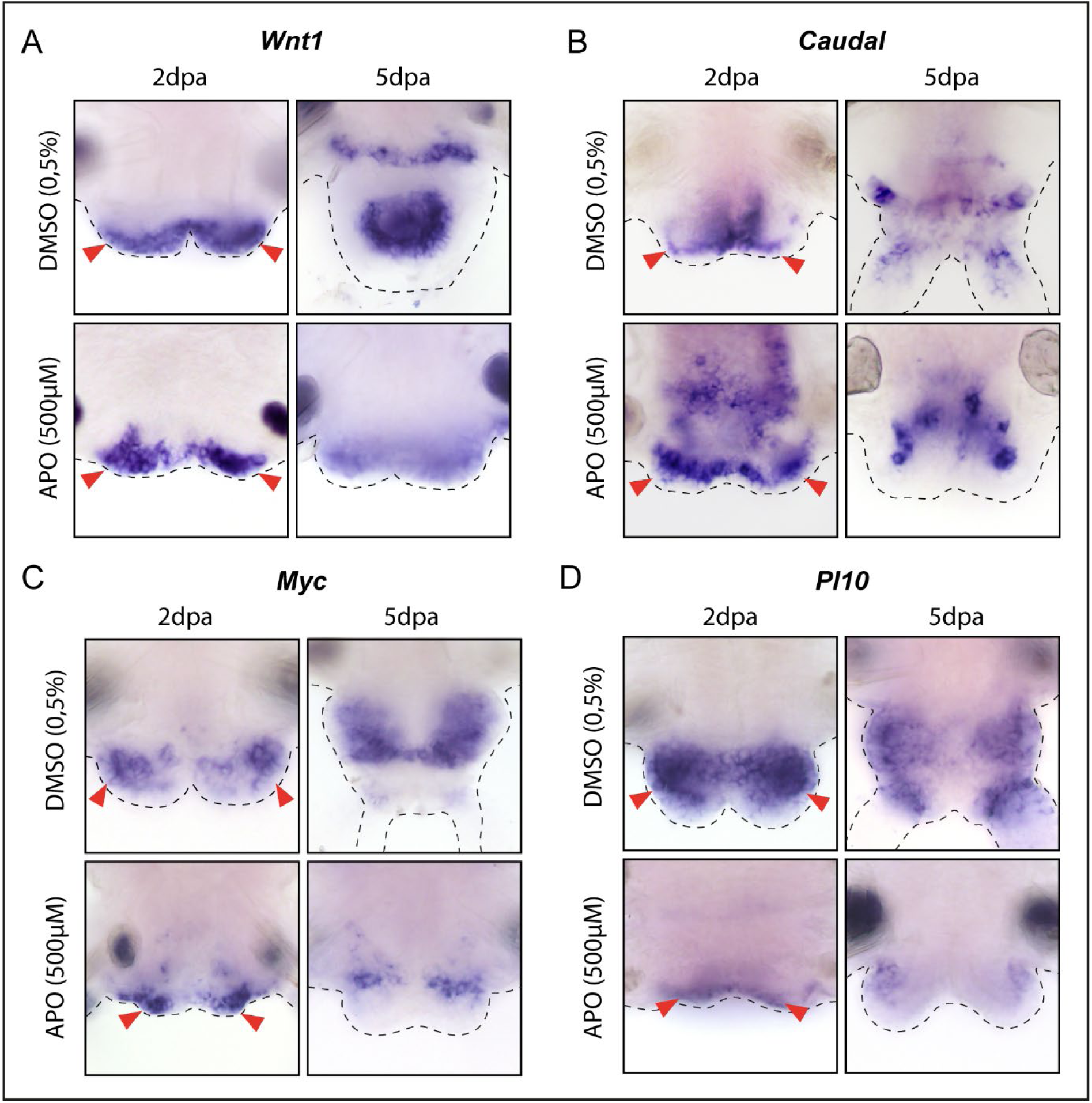
Effects of ROS inhibition on ectodermal gene expression during *Platynereis* regeneration.

*Whole-mount* in situ *hybridizations for ectodermal markers at 2 and 5dpa in APO-treated worms and controls:* Wnt1 *(A),* Caudal *(B),* Myc *(C) and* Pl10 *(D). Red arrowheads: ectodermal labelling in control worms. Anterior is up. Ventral views*.

Taken together, these results suggest that under ROS inhibition meso-endodermal specification is initiated but blocked at a 2dpa-like stage, consistent with the outer morphology of APO-treated worms, while ectodermal identity is altered. This distinct behaviour of meso-endodermal *versus* ectodermal tissues can be put into perspective with the proliferation profile observed in APO-treated worms, with most of the residual proliferation taking place in the ectoderm (Figure 12A).

## Discussion

In the present study we have assessed 89 genomes (85 metazoans and 4 non-metazoan species) for the absence/presence of orthologous genes associated with ROS metabolism. We highlighted phylum-or species-specific gene losses and duplications as well as a rich evolutionary history of ROS-associated genes, providing insight into the complexity of ROS metabolism evolution. Thus, we have assessed the expression of the ROS-metabolisms genes during the process of regeneration in various species. We highlighted that during evolution each species seems to have retained a specific combinatorial set of genes regulating the ROS metabolism. In addition, we observed complex spatio-temporal expression dynamics that in some organisms, reflect the sequential deployment of the ROS metabolism modules during regeneration. Using the annelid *Platynereis dumerilii* and the anthozoan cnidarian *Nematostella vectensis* to perform a comparative study, we have shown that ROS is required in both species for a successful regeneration of lost body parts. We further show that ROS metabolism in both species does not involve the same enzymes and affect apoptosis and cell proliferation differently. Last, we showed a tissue differential response to the ROS inhibition in both species suggesting tissue-specific signaling. Overall this work highlights strong variations in the previously proposed conserved ROS, apoptosis and cell proliferation relationship, revealing the mechanistic plasticity of ROS metabolism required for metazoan regeneration.

### A dynamic metabolism arsenal ensures ROS signaling functions among metazoan species

#### The evolutionary history of ROS metabolism actors is shaped by multiple gene losses and duplications and key co-evolution event

The ROS metabolism actors represent major, often ancient, enzymatic families (Inupakutika et al. 2016) that are supposed to be widespread across Metazoa (Hewitt and Degnan 2022; Hewitt and Degnan 2023). Our extensive exploration of genomic data from 85 species representing the major metazoan lineages reveal that the numbers of genes encoding most of the redox arsenal vary strikingly among lineages and within a specific lineage from one species to another. This highlights the fact that a multitude of gene losses and duplications occurred in such gene families in all main lineages of animals (Figure 3).

Duplication events have resulted in an unexpected redundancy between the ROS metabolism actors. Such redundancy for enzymes of a same gene family in particular the GPX, the Sod Cu/Zn, and the Nox and Duox, suggest sub-functionalization and potential new gene functions, associated with specific pattern of subcellular localization and tissue distribution (Zhang 2003; Magadum et al. 2013). This was notably demonstrated for *Nox/Duox* genes in vertebrates which play different roles in various tissues (hormone synthesis in thyroid, sperm capacitation in testicles etc) (Moreno et al. 2002; Geiszt et al. 2003). Redundancy can also be observed across gene families at the level of the functions carried out by those enzymes. For example, Sod Cu/Zn and Sod Mn (proposed to be unrelated gene families, Fink and Scandalios 2002) involved in ROS conversion and GPX and catalase, in charge of ROS detoxification, seem functionally redundant. Upon loss of one or the other enzyme, an alternative element could play the required function, making the system extremely robust, even if the fine catalytic reaction is not similar. Gene losses are also a major driving force leading to genome evolution (Albalat and Cañestro 2016) and biological processes plasticity, in this specific case the evolution of ROS metabolism.

One noticeable example is the coevolving Duox and DuoxMF gene families. A complex evolutionary history sculpted by various gene duplications and losses resulted in two gene copies in mammalians that are not homologous to the ones present in the other bilaterian species. Following the mammalian-specific duplication, the mammalian sets of genes have specialized, with Duox2 carrying out specific antimicrobial functions in the gut and hormone synthesis functions in the thyroid (Miot and De Deken 2023). While the coevolution of Duox and Duox MF suggests a high selective pressure retains their association, both genes are lost in some phyla (cnidarians, platyhelminths), in which their function might still be carried out by other Nox genes.

Hence, despite this eventful evolutionary history, the immense majority of investigated species have at least a ROS-producing enzyme (Nox or Duox plus Duox MF), a ROS-converting enzyme (Sod Cu/Zn and Sod Mn plus the chaperone CCS), a ROS detoxifying enzyme (catalase or GPX) and to a lesser extent (see below) both actors of the antioxidative response regulation (Nrf and Keap1). Altogether, this suggests that all species have a core ROS metabolism toolkit that is not necessarily the same. This in turn supports the idea that the selective pressure on the genes/proteins level is low, while at the level of the functions of a specific ROS module it is probably much more constrained.

#### Additional complexity of the ROS regulatory module evolution

Nrf2 and its inhibitor Keap1 are known key actors of the antioxidative response in mammalians, regulating the transcription of ROS metabolisms genes according to the cellular redox balance. Upon oxidative stress, the inhibitory association of Keap1 with Nrf2 is disrupted, and Nrf2 migrates to the nucleus to activate the transcription of detoxification genes (Bellezza et al. 2018). Based on studies in *Drosophila* and the identification of NRF and Keap1 orthologs in other species, it has been proposed that this key role might be evolutionarily conserved (Kobayashi et al. 2002; Sykiotis and Bohmann 2008; Choe et al. 2009) (our own results). However, our study shows that while this may be true for the majority of bilaterian species, the situation for non-bilaterians is radically different and challenges this view. Indeed, Porifera, Placozoa and several Cnidaria species lack Keap1, while Ctenophora have neither Keap1 nor Nrf. This is especially surprising as many of poriferan and cnidarian species are exposed to high levels of ROS, when they harbor phototrophic symbionts (Richier et al. 2005), or are themselves hotspot ROS producers (Taenzer et al. 2023).

The case of cnidarians with only some species possessing both Nrf and Keap1 orthologs is even more puzzling and one can wonder if Keap1 and Nrf really act as regulators of the antioxidative response in those species. Indeed, in the absence of diagnostic residues in both Nrf (DLG/ETGE) and Keap1 (key cysteine), interaction and regulation are uncertain. A constitutive activity of Nrf, that would therefore not require the ROS-induced inactivation of Keap1, could be envisaged for these poriferan and cnidarian species. However, it is known that sustained Nrf2 activation is lethal during mice development (Wakabayashi et al. 2003).

Another possibility could be the presence of alternate regulations of Nrf activity, as demonstrated for nematodes. In *C. elegans,* Nrf ortholog SKN1 (a divergent gene, also lacking DLG and ETGE motifs) operates in association with its inhibitor WDR-23 (a WD40 repeat protein, unrelated to Keap1), thanks to a specific motif interaction, meaning that an alternate regulatory system for the antioxidative response might have emerged in this phylum (Choe et al. 2009; Lo et al. 2017). In addition, it is well documented that within the Nrf superfamily, Nrf2 is involved in the antioxidative response (Zhang et al. 2009; Zhao et al. 2011). This is not the case for Nrf1 and Nrf3, two transcription factors that are involved in protein and lipid homeostasis (Radhakrishnan et al. 2010; Lee et al. 2013; Waku and Kobayashi 2021). As Nrf1, 2 and 3 result from a relatively recent duplication in gnathosomes, the ancestral function of Nrf in metazoans remains unknown. Functional and mechanistic studies on key model species, in a comparative approach, may help to further understand these points and to further unravel the evolutionary history of the antioxidative response in animals.

### ROS as a universal trigger of animal regeneration?

#### Injury-induced ROS production, albeit through different mechanisms, are required for regeneration in *Platynereis* and *Nematostella*

ROS play a myriad of functions in animals ranging from wound healing to development (Cordeiro and Jacinto 2013; Rampon et al. 2018). They have been also shown to be involved in the initiation of restorative regeneration in several species such as zebrafish, *Xenopus*, *Drosophila*, *Hydra* etc (Gauron et al. 2013; Love et al. 2013; Fogarty et al. 2016; Suknovic et al. 2022). With the aim to extend and deepen these observations on the role of ROS during the initiation of regeneration, and to test if conserved or distinct ROS metabolism arsenals modulate such biological functions in metazoans, we investigated ROS production during regeneration in two complementary models, *Platynereis dumerilii* and *Nematostella vectensis*. The first one is an annelid, a group for which almost no data existed so far regarding ROS involvement in the initiation of regeneration, and it possesses a complete ROS arsenal (Sup Table 4). The second one is an anthozoan cnidarian. This sea anemone possesses members of all major modules, but lacks *Duox* and *Duox Maturation Factor* from the ROS-producing module as well as *Keap1*, from the antioxidant regulatory module (Sup Table 4). Thus, it provides a crucial point of comparison among metazoans and within cnidarians, especially with *Hydra* which possesses a Keap1 ortholog, and in which ROS signaling has been shown to be important for apical regeneration.

We showed that in both species, ROS are produced very early upon amputation (Figure 8), while the type of ROS (H_2_O_2_ *versus* non-H_2_O_2_ for *Nematostella* and *Platynereis* respectively) (Figure 8), as well as the enzymes in charge of their detoxification (GPX for *Platynereis* and catalase for *Nematostella*) differ between the two models (Figure 9). Data from the literature show that early ROS production is widespread following amputation (zebrafish, Gauron et al. 2013; *Xenopus*, Love et al. 2013; *Hydra*, Suknovic et al. 2022; axolotl, Al Haj Baddar et al. 2019; planarian, Pirotte et al. 2015). However, the ROS subtypes involved and the enzymatic activity underlying this production and detoxification are rarely established, except in *Danio* and *Hydra*. In the zebrafish, H_2_O_2_ was detected during caudal fin regeneration using the genetic probe HyPer (Meda et al. 2016), enzymatic assays indicate an increase of Sod activity at 15hpa, consistent with the peak H_2_O_2_ level, while catalase activity increases later (72hpa) and might contribute to H_2_O_2_ detoxification (Thauvin et al. 2021). In *Hydra*, both H_2_O_2_ and mitochondrial O_2_ ^-^ are produced during apical regeneration (detected using MitoSox and Amplex UltraRed respectively), and similarly to zebrafish, an increase of Sod activity is observed (10 minutes post-amputation), followed by an increase of catalase activity (30 minutes post-amputation) (Suknovic et al. 2022).

We also showed that ROS signalization is mandatory for regeneration in both species, as the process is blocked at early stages in both species (*i.e.*, prior to the onset of cell proliferation) following the inhibition of ROS production (Figure 10). This is reminiscent of observations in other models, in which ROS inhibition also impairs regeneration (zebrafish, Gauron et al. 2013; *Xenopus*, Love et al. 2013; axolotl, Carbonell et al. 2021; *Hydra*, Suknovic et al. 2022; planarian, Bijnens et al. 2021). We also showed that the expression of several antioxidative genes depends on ROS production, suggesting that a transcriptional antioxidative response takes place during regeneration in *Platynereis* and *Nematostella*, although we do not know if this transcriptional regulation relies on Nrf orthologs. In other models, amputation induces an up-regulation of several putative ARE genes encoding actors of ROS metabolism: *trxn* in gecko (Zhang et al. 2016), *gsr* in planarian (Bijnens et al. 2021), *prdx* and *gpx* in zebrafish heart (Han et al. 2014), *txn* and *prdx* in axolotl (Zhulyn et al. 2023). Whether their transcription is regulated by ROS *via* Nrf orthologs is unknown.

#### Distinct downstream effects of ROS on injury-induced apoptosis and cell proliferation in ***Nematostella* and *Platynereis***

The combined action of ROS metabolism modules (production, detoxification and regulatory module) allows a fine tuning of ROS production within tissues, which is required for their signaling functions during regeneration. Indeed, in many different species, ROS act as upstream signals by tightly orchestrating major cellular events such as apoptosis (zebrafish Gauron et al. 2013; *Hydra* Suknovic et al. 2022; *Drosophila* Fogarty et al. 2016) and proliferation (zebrafish Gauron et al. 2013, Han et al. 2014; *Xenopus* Love et al. 2013; planarian Van Huizen et al. 2019), and triggering molecular pathways to induce a regenerative answer (Vriz et al. 2014; Vullien et al. 2021).

In contrast with other models (zebrafish, *Drosophila, Hydra*), in *Platynereis*, ROS inhibition does not affect injury-induced apoptosis (Figure 11A, B). However, it is important for injury-induced cell proliferation taking place during posterior regeneration. In the presence of ROS, cell proliferation rate is low at 1dpa, increases at 2dpa, and is at its highest from 3dpa to 5dpa, with about 50% of S-phase cells in the blastema and regenerating structures (Planques et al. 2019; Bideau et al. 2024). Proliferation is known to be required for regeneration to proceed beyond the blastemal stage; however, blastema formation itself is proliferation-independent (Planques et al. 2019). Under ROS inhibition, injury-induced proliferation is significantly reduced as early as 2dpa. This reduction is mostly observed in the meso-endoderm, while proliferation still takes place in the ectoderm (Figure 12A, B). Hence, ROS in *Platynereis* are required for regeneration independently of apoptosis, by inducing cell proliferation in a tissue-specific manner.

In *Nematostella*, three waves of apoptosis have been reported (Johnston et al. 2021): the first shortly after injury, coinciding with the wound healing process (2-6hpa), the second at the onset of cell proliferation initiation (∼24hpa), and the third when the pharynx is reformed (∼60hpa). Inhibition of apoptosis (all three waves combined) blocks cell proliferation and regeneration (Johnston et al. 2021). While inhibition of ROS has no effect on the first wave of apoptosis, it strongly reduces / blocks the second wave of apoptosis at 24hpa (Figure 11C, D). Interestingly, specific inhibition of the second wave of apoptosis does not block regeneration (Johnston et al. 2021), suggesting that ROS also affect other mechanisms that, combined with the second wave of apoptosis, are crucial for regeneration in *Nematostella*. Injury-induced cell proliferation in *Nematostella* is observed as early as 16-24hpa (Passamaneck and Martindale 2012; Amiel et al. 2015), leading to a burst of proliferative cells at 48hpa at the amputation plane in the body wall epithelia and in the oral part of the mesenteries. While the first step of regeneration (up to 24hpa) is cell proliferation independent, the completion of regeneration is stacked when cell proliferation is blocked (Amiel et al. 2015). Unexpectedly, despite the inhibition of the second apoptotic wave and of the regeneration process, cell proliferation is not totally blocked but stalled at the level usually observed in 48hpa polyps with strong indications that this reduction is mainly restricted to the body wall epithelial at the amputation plane (Figure 12E, F). Hence, ROS in *Nematostella* are required for regeneration by affecting a second wave of apoptosis and cell proliferation in a tissue-specific manner.

#### ROS involved in inter-tissue communication during regeneration

Our results highlight a potential involvement of ROS in tissue-specific responses and/or inter-tissue communication in *Platynereis* and *Nematostella*.

In *Platynereis*, the distinct effects of ROS inhibition on the ectoderm and the meso-endoderm affect their molecular identity. The near absence of proliferation in the meso-endoderm might be responsible for “freezing” its molecular identity under APO treatment at a 2dpa-like stage (Sup Fig 16), while the residual proliferation drives an identity change in the ectoderm (Figure 13). Whether the tissue identity progresses further into regeneration as a result or is merely altered cannot be determined in the absence of ectodermal markers for the later stages of regeneration. These specificities of the ectoderm and meso-endoderm can also be analyzed with regard to ROS production itself, which mostly takes place in epithelial cells between 1 and 48hpa (Figure 8A), suggesting an inter-tissue communication between two compartments during regeneration: (1) the ectoderm, which produces ROS early following amputation, and in which proliferation is partially ROS-independent, allowing tissue identity to evolve in the absence of ROS, and (2) the meso-endoderm, which perceives a ROS-induced mitogenic signal from the neighbouring ectodermal cells, and in which proliferation is ROS-dependent, causing tissue identity to stall completely in the absence of ROS.

We know from a previous study in *Nematostella* that a crosstalk between the epithelia of the body wall and the oral part of the mesenteries is crucial at the amputation plane to induce cell proliferation, and for the successful outcome of regeneration (Amiel et al. 2019). Although ROS inhibition does not prevent this contact (Figure 10), its tissue-specific impact on cell proliferation suggests that ROS might be involved in the molecular crosstalk following the initial tissue contact. More precisely, ROS might be involved in the attraction of cells within the mesenteries towards the body wall epithelium. In fact, between 24-48hpa mesenteries cells migrate towards the amputation plane body wall epithelium, a population of these cells seem to accumulate at the oral tip of the mesenteries without being able to pursue their migration into the body wall epithelium (Amiel et al. 2019). The tissue crosstalk *per se* and the involvement of ROS in this process and in particular in the cell migration, remains to be determined.

Molecular crosstalks in regenerative contexts are mostly documented at the cellular level, for example between immune cells and resident cells (including stem cells) of the injured tissues involved in bidirectional paracrine signalling (Harrell et al. 2021; Guenin-Mace et al. 2023; Manneken and Currie 2023). Interestingly, ROS have been shown to contribute to such inter-cell types communication. In zebrafish, ROS act as a chemo-attractant after caudal fin injury to recruit leukocytes, to the wound site (Niethammer et al. 2009; Yoo et al. 2011). In *Drosophila* imaginal disc, injury-induced ROS production activates haemocytes, that in turn trigger apoptosis in the damaged tissues via TNF secretion and JNK signalling (Fogarty et al. 2016). However, we have little information as to how ROS might contribute to communication between different tissues during regeneration, mostly because the cells responsible for ROS production or ROS-sensing are rarely identified.

#### Evolutionary variations of the ROS, apoptosis, proliferation relationship highlight independent ROS-related mechanisms driving metazoan regeneration

Results from zebrafish caudal fin regeneration (Gauron et al. 2013), *Drosophila* imaginal disc regeneration (Fogarty et al. 2016) and *Hydra* oral regeneration (Chera et al. 2009; Chera et al. 2011; Suknovic et al. 2022) have shown that injury-induced ROS production induces apoptosis that in turn leads to compensatory cell proliferation. Such a sequence of signalling has been proposed as a conserved strategy at the onset of animal regeneration (Diwanji and Bergmann 2018). Additional data support a link between ROS and cell proliferation during zebrafish heart (Han et al. 2014), *Xenopus* tail (Love et al. 2013), *Drosophila* gut (Buchon et al. 2009) and planarian whole body (Van Huizen et al. 2019) regeneration. However, in these studies the link between ROS and apoptosis is not evidenced or explored.

Our results indicate that a “ROS -> apoptosis -> proliferation” cellular cascade triggering the onset of regeneration in animals is not as conserved as proposed. In fact, ROS in *Platynereis* are crucial for cell proliferation in the endo-mesoderm (to a lesser extent in the ectoderm) and do not seem to be involved in triggering apoptosis following injury. Injury-induced ROS in *Nematostella* seem crucial for the second wave of apoptosis and cell proliferation balance between the body wall epithelia and the mesenteries. These results highlight a more fine-tuned, and tissue-specific mechanism. In addition, ROS signaling in *Nematostella* differs from *Hydra*, for which an entire ROS > apoptosis > proliferation cascade is reported. During apical regeneration in *Hydra*, ROS production induces the MAPK/ERK pathway, causing the phosphorylation and nuclearization of CREB, which in turn triggers apoptosis (Chera et al. 2011; Suknovic et al. 2022). The release of Wnt3 ligand by apoptotic cells subsequently induces proliferation (Chera et al. 2009). In contrast, MAPK/ERK signaling is not required for apoptosis during *Nematostella* oral regeneration (Johnston et al. 2021), suggesting the induction of the second wave of apoptosis by ROS might rely on another signaling pathway. Indeed, various signaling pathways (Wnt/βcatenin, JNK, MAPK/ERK …) are induced by ROS during regeneration in different species (Chera et al. 2011; Gauron et al. 2013; Love et al. 2013; Santabárbara-Ruiz et al. 2015; Jaenen et al. 2021), also supporting the idea of a non-similar cascade and indicating a loose selective pressure on the diverse molecular and cellular elements downstream of ROS.

Hence, ROS appear to be positively selected, either as an ancestral or convergently acquired trait, to initiate regeneration at the metazoan scale, but the fine molecular and cellular signalling to achieve this role is different and probably less constrained. This could be correlated with the strong selective pressure acting at the functional level of the specific ROS modules allowing a tight regulation of the ROS levels, which are important for triggering regeneration.

## Material & Methods

### Comparative genomics and phylogenetic analyses

Putative members of the ROS metabolism machinery gene families (n=24, Figure 1) were identified by reciprocal best BLAST hit approach on available transcriptomic and genomic data for 85 species of interest spanning all major metazoan lineages, plus 4 non-metazoan species, using *H. sapiens* sequences as query on the BioEdit software (v7.2.5) (Hall 1999). A list of the databases used to retrieve these sequences, of the corresponding species, their abbreviations as well as their taxonomic classifications are available in Sup Table 1. After manual checking, redundant sequences were discarded, and in some cases incomplete overlapping sequences were concatenated. All identified sequences are available in supplementary documents (Sup Files 1-8).

Sequences were then aligned using MUSCLE (Gabler et al. 2020) (available on the MPI Bioinformatics Toolkit platform, Zimmermann et al. 2018) and alignments were curated either manually using SeaView (v5.0.5) (Gouy et al. 2010), or automatically using the BMGE software (v1.12.1) (Criscuolo and Gribaldo 2010). Maximum likelihood phylogenetic analyses were performed using the PhyML 3.0 software (Guindon et al. 2010). The amino acid substitution model was selected by SMS (Smart Model Selection) (Lefort et al. 2017) with Bayesian Information Criterion. Starting tree was generated by BioNJ (Gascuel 1997), and statistical branch support was assessed by aBayes fast likelihood method (Anisimova et al. 2011). Resulting phylogenetic trees were visualized and designed with FigTree 1.4.4 and Adobe Illustrator. Species names were abbreviated using the first letter of the genus name followed by the three first letters of the species name.

Conserved domains were identified in Nox/Duox, SOD, Nrf and Keap1 sequences using InterProScan (Paysan-Lafosse et al. 2023) and NCBI CD-search (Wang et al. 2023).

NCBI Genome Data Viewer (https://www.ncbi.nlm.nih.gov/gdv) was used to perform microsynteny analysis by searching available genomes of species from our dataset for *Duox* and *DuoxMF* genes.

Nrf and Keap1 association was investigated using PSOPIA (Murakami and Mizuguchi 2014) to obtain interaction scores between pairs of proteins, and AlphaFold 3 (Abramson et al. 2024) to generate predicted structures for Nrf and Keap1 interaction.

Ancestral state reconstruction was performed using Mesquite (version 3.81). Character matrices and species trees were constructed manually, and ancestral state reconstructions were performed using the “Trace all characters” tool, using both parsimony and Maximum-Likelihood methods. A few unlikely reconstructions (suggesting successive loss and reacquisition of a single sequence) were resolved manually using parsimony principles.

### Gene expression levels

Genes involved in ROS metabolism were identified in the *Platynereis* reference transcriptome (Paré et al. 2023). Their expression levels at different regeneration stages (*i.e.* 0, 1, 2, 3, 5 days post amputation as well as non-amputated control) were retrieved from RNA-seq datasets previously produced in the lab (Paré et al. 2023). Similarly, ROS metabolism genes in *Nematostella vectensis* were identified and their associated levels of expression at different regeneration stages (*i.e.* 0, 2, 4, 8, 12, 16, 20, 24, 36, 48, 60, 72, 96, 120, 144 hours post amputation as well as non-amputated control) obtained using the NvERTx database (https://nvertx.ircan.org/ER/ER_plotter/home) (Warner et al. 2018). Expression data for other model organisms were retrieved from Sinigaglia et al. 2022 (*Paryhale hawaiensis,* 0, 12, 24, 36, 48, 60, 72, 84, 96, 108, 120hpa and non-amputated control), Chang et al. 2017 (*Xenopus tropicalis*, 0, 6, 15, 24, 72hpa and non-amputated control), Nauroy et al. 2019 (*Danio rerio*, 0, 2, 3, 10dpa) and Wenger et al. 2019 (*Hydra vulgaris*, 0, 0.5, 1, 2, 4, 8, 16, 24, 36, 48hpa and non-amputated control). For all species, the list of genes, accession numbers and expression level data at different time points during the course of regeneration are provided in [Sup Table 3].

Expression level dynamics were visualized and heatmaps produced using Heatmapper (Babicki et al. 2016) and Adobe Illustrator.

#### *Platynereis* breeding culture, amputation procedure and biological material fixation

*Platynereis* juvenile worms were obtained from a culture established in the Institut Jacques Monod following general guidelines by Dorresteijn 1990 and Vervoort and Gazave 2022. Worms aged 3 to 4 months and spanning 30-40 segments were selected for the experiments. Amputations were performed after anaesthesia in MgCl2 7.5% and sea water solution (1:1), using a microknife to remove the pygidium and up to 5 posterior segments as detailed in Planques et al. 2019 and Vervoort and Gazave 2022.

For Whole-Mount *In Situ* Hybridization (WMISH), TUNEL and antibody labelling experiments, regenerating parts of the worms were collected and fixed at specific timepoints following amputation in 4% PFA diluted in PBS Tween-20 0.1% (PBT) during 2 hours at RT, then rinsed in PBT, gradually dehydrated in 100% MeOH and stored at -20°C. For EdU proliferation labelling, regenerating worms were incubated with 5µM EdU in Natural Filtered Sea Water (NFSW) for 1h at RT before sample collection and fixation (see Results for specific EdU incubation and amputation times). For phalloidin labelling experiments, samples were similarly produced while not dehydrated and stored directly in PBT at 4°C. For fluorescent ROS labelling, regenerating worms were incubated with 30µM CellROX (C10444, Invitrogen) in NFSW for 1h at RT. Regenerating parts were then collected and subjected to a light fixation (3.7% formaldehyde, 15min, RT), and readily treated for imaging.

#### *Nematostella* breeding culture, bisection procedure and biological material fixation

*Nematostella* subjects were obtained from a culture established at the Institute for Research on Cancer and Aging, Nice (IRCAN), following guidelines by Hand and Uhlinger for maintenance, feeding and spawning (Hand and Uhlinger 1992). Juveniles or adults (depending on experiment) were relaxed on a light table for 10-15 minutes before adding 7.14% MgCl_2_ (1:5 in 1/3 Artificial Sea Water; ASW) for anaesthesia. Polyps were then cut using a microsurgery scalpel n°15A (Swann-Morton, Sheffield, UK), perpendicular to the oral–aboral axis of the body column.

For Whole-Mount *In Situ* Hybridization (WMISH), amputated or uncut polyps were fixed with 4% PFA, 0.1% glutaraldehyde in 1/3 ASW (2min, on ice) then with 4% PFA, 0.5% DMSO in 1/3 ASW (1h, on ice, in the dark). Fixed samples were rinsed with PBT (5 times) before transferring to MeOH 100%.

For sample fixation prior to staging, polyps were relaxed on the light table and using MgCl_2_ before fixation in 4% PFA in 1/3 ASW for 1 h at RT. Fixed animals were then washed three times in PBT and counterstained with Hoechst (1: 5000) and BODIPY FL PhallAcidin 488 (1: 200) to visualize global morphology. For EdU proliferation labelling, regenerating polyps were incubated with 100μM EdU in 1/3 ASW for 30min at RT before collection (see Results for specific EdU incubation and amputation times). Samples were then fixed in 4% PFA, 0.2% glutaraldehyde in PBS 0.1% Triton X-100 (PBTx 0.1%) for 2min on ice, then in 4% PFA in PBTx 0.1% for 1h on ice and rinsed in PBTx 0.1%. For TUNEL labelling, regenerating polyps were fixed in 4% Formaldehyde in 1/3 ASW for 1h at RT or overnight at 4°C, then washed in PBS 0.2% Triton X-100 (PBTx 0.2%).

#### Gene cloning, probe synthesis and Whole-mount *in situ* hybridization (WMISH)

Fragments of ROS metabolism genes of interest were amplified by PCR using specific primers on cDNA obtained from mixed stages of *Platynereis* and *Nematostella* embryos, larvae and regenerated structures, then cloned into pMiniT 2.0 (NEB) and pGEM-T Easy (Promega) vectors respectively and sequenced, as described in Gazave et al. 2013 and Amiel et al. 2017 (Sup File 9; Sup File 10). Validated plasmids were used as template to produce RNA antisense probes labelled with digoxigenin (DIG) for WMISH.

WMISH in *Platynereis* were performed as described in Demilly et al. 2013 and Vervoort and Gazave 2022. Fixed samples were rehydrated, washed in PBT then treated with 40 µg/ml proteinase K PBT for 10min, 2mg/ml glycine PBT for 1min, 4% PFA PBT for 20min and finally washed in PBT. The samples were then incubated overnight at 65°C with DIG-labelled RNA antisense probes in hybridization buffer (Vervoort and Gazave 2022) then washed in SSC buffers and formamide at 65 °C. Blocking was then performed during 1h with PBT and sheep serum, before incubating the samples with anti-digoxygenin-alkaline phosphatase antibody (Roche, 1:4000) for 1h. After washing in PBT, samples were incubated in staining buffer (Tris-NaCl pH9.5 + MgCl_2_) for 10 min, then in staining buffer complemented with NBT/BCIP until full staining. Stained samples were washed in Tris-NaCl buffer pH7.5 and PBT then transferred in glycerol 87% before imaging.

WMISH protocol in *Nematostella* was adapted from Genikhovich and Technau 2009. Fixed samples were rehydrated, washed in PBT then treated with 10 µg/ml proteinase K PBT for 20min, twice 2mg/ml glycine PBT for 5min, 1% triethanolamine PBT with increased concentration of acetic anhydride (0, 1.5 and 3µL *per* 500µL), 3.7% PFA PBT for 1h and finally washed in PBT. The samples were incubated in hybridization buffer overnight at 65 °C, then with DIG-labelled RNA antisense probes in hybridization buffer overnight at 65 °C. Washing in solutions of hybridization buffer with increasing proportion of SSC buffer, then solutions of SSC buffer with increasing proportion of PBT, were performed prior to blocking during 1h in blocking solution (10% Boehringer-Mannheim Blocking reagent in maleic acid buffer). Samples were incubated with an anti-digoxygenin-alkaline phosphatase antibody (Roche, 1:5000) in blocking solution for 1h, then in staining buffer (Tris-NaCl pH9.5 + MgCl_2_) complemented with NBT/BCIP until full staining. Stained samples were washed in PBT then transferred in glycerol 87% before imaging.

### Antibody staining, EdU cell proliferation assay, TUNEL cell death assay, phalloidin and CellROX labelling in *Platynereis*

For immunostaining, EdU and TUNEL assays, samples were rehydrated, PK-digested and post-fixed (as described in the previous section). Then, for EdU and TUNEL labelling, samples were labelled using respectively the Click-It EdU Imaging Kit (488 or 555nm, ThermoFisher, as in Bideau et al. 2024) or the Click-iT TUNEL kit (647nm, ThermoFisher, as in Demilly et al. 2013). For immunostaining, samples were then incubated 1h in PBT and sheep serum, then with anti-acetylated tubulin (Sigma T7451, 1:500) antibodies which label the neurites. After several washes in PBT and another hour of blocking with sheep serum, samples are incubated with fluorescent secondary anti-mouse IgG Alexa Fluor 488 or 555 conjugate (Invitrogen, 1:500). For phalloidin labelling, samples were incubated in phalloidin-Alexa 555 (Molecular Probes, 1:100) overnight at 4°C. Following those labelling, samples were counterstained overnight at 4°C using DAPI (1:1000) before transferring the samples in glycerol/DABCO.

ROS-producing cells were labelled by incubating regenerating worms at different time points with 30µM CellROX Green (ThermoFisher) in NFSW for 1h at RT, then regenerating structures were collected and subjected to a light fixation (procedure described above), PBT washes and nuclei counter-staining (DAPI 1:1000, 1h, RT), before rapid observation.

### EdU cell proliferation assay and TUNEL cell death assay in *Nematostella*

For cell proliferation labelling, samples were permeabilized in PBS 0.5% Triton X-100 (PBTx 0.5%) for 20min, then stained using a Click-It EdU Imaging Kit (Invitrogen #C10337, as in Passamaneck and Martindale 2012). For apoptosis labelling, samples were permeabilized with Proteinase K 10 µg/ml in PBS for 20min at RT, rinsed in PBS, re-fixed with 4% Formaldehyde in PBS1x, rinsed in PBS and treated using a *In Situ* Cell Death AP kit (Roche 11684809910). Following those labelling, samples were rinsed in PBS and counterstained overnight at 4°C using DAPI (1:5000, 1h, RT) before transferring the samples in 87% glycerol and observation.

### H2O2 detection in *Nematostella*

Regenerating or control 5mm-long subadults were left to incubate in 80µL 1/3 ASW each for one hour at RT. Following incubation, 50µL of supernatant were harvested for each polyp and transferred into a black 96-well fluorescence microplate. Twelve polyps were used *per* timepoint. 50µL of reaction solution (0.2 U/mL HorseRadish Peroxidase [ThermoFisher #31491] + 100µM Amplex UltraRed [Invitrogen # A36006] in 1/3 ASW) was added to each well, and fluorescence reading (ex 520nm / em 580nm) performed readily using a plate reader (SAFAS Xenius XM). The background value was obtained by adding 50µL of reaction solution to 50µL of 1/3 ASW.

### Detection of catalase, superoxide dismutase and peroxidase enzymatic activity

Following amputation, regenerated parts (plus the posterior-most segment) of *Platynereis* or 1/8^th^ of the body abutting the amputation plane in *Nematostella* were collected at different regeneration timepoints or stages (as defined in Planques et al. 2019 and Amiel et al. 2015). For Non-Amputated control samples, the five posterior-most segments (*Platynereis*) or 1/8^th^ of the body comprising the head (*Nematostella*) were collected. Protein extractions were immediately performed on fresh tissues. A hundred worms or ten subadult polyps were used *per* replicate *per* time point. Fresh tissues were homogenized in 20mM HEPES buffer pH 7.2 (1mM EGTA, 210mM mannitol, 70mM sucrose) and centrifugated at 1.500g for 5 minutes at 4°C. The supernatant was stored at -80°C. At least three biological replicates were generated for each time points. Enzymatic activity assays and calculations were performed following instructions from kits #706002 (SOD), #707002 (catalase) and #703102 (GPX) by Cayman Chemicals. Results were normalized by sample concentration.

### ROS inhibition treatments and scoring

Pharmacological inhibition of ROS production was performed using two inhibitors: APO (178385-1GM, Sigma) and DPI (D2926-10MG Sigma). 100mM and 10mM stock solutions respectively were prepared in 100% DMSO. Efficient concentrations leading to a robust and reproducible phenotype (APO 500µM and DPI 10nM in *Platynereis*, DPI 100nM in *Nematostella*) were determined experimentally (see Sup Fig 14).

Regenerating worms and polyps were incubated during a specific time window in NFSW and 1/3 ASW respectively, containing either APO or DPI at the efficient concentration (see results for details about the time windows and concentrations used). Control worms and polyps were incubated in NFSW or ⅓ASW containing similar concentrations of DMSO. All solutions were changed every 24 hours to maintain proper inhibitor activity for the whole duration of the experiments. Worms and polyps were then fixed, as described above, before subsequent experiments. For scoring experiments, individual *Platynereis* worms were incubated in 2 ml of inhibitor solution or DMSO control solution on 12-wells plates, and scored daily to assess their regeneration stage, as described in Vervoort and Gazave 2022, while *Nematostella* juvenile polyps were incubated by groups of up to 50 individuals in 1 ml of inhibitor solution or DMSO control solution on 24-wells plates, and scored at the end of the regenerative process (7dpa), as precise staging requires fixation and DAPI staining of individuals (staging system from Amiel et al. 2015).

### Images acquisition and analyses

A Leica DM 5000 B microscope equipped with a CTR 5000 control unit was used for *Platynereis* colorimetric WMISH (brightfield) acquisition. A Zeiss Imager Z1 microscope was used for *Nematostella* colorimetric WMISH (brightfield) and DAPI/phalloidin labelling acquisitions. Fluorescent confocal images of samples were acquired with Zeiss LSM780 (*Platynereis*) and LSM800 (*Nematostella*) confocal microscopes. Image processing (contrast and brightness, z-projection, virtual transversal views etc) was performed using FIJI and Abode Photoshop. Figures were designed with Adobe Illustrator. Cell counting (for ROS+ cells, EdU+ cells and TUNEL+ cells) were performed using the IMARIS 9.5.0 software (Oxford Instruments) following an automatic cell counting procedure as defined in Bideau et al. 2024.

### Statistical analyses

All statistical tests and associated graphic representations were performed using GraphPad Prism 9. Mann–Whitney U tests were used to compare proportions of CellROX+ cells, enzymatic activities and H_2_O_2_-related Amplex UltraRed fluorescence rate between different timepoints over the course of regeneration. Multiple Mann–Whitney U tests were used to compare regeneration progress in APO and DPI-treated *versus* control subjects, and to compare EdU+ and TUNEL+ cell proportions between different experimental conditions.

## Acknowledgments

We thank the Institute Jacques Monod facility staff for their help with the *Platynereis* culture. We thank Renaud Rebillard and the IRCAN Marine invertebrate facility for (anti)Aging research (ANTIAGE) for *Nematostella* husbandry and care. Equipment acquisition for the ANTIAGE facility was supported by Université Côte d’Azur - IDEX UCAJedi (ANR-15-IDEX-01), Région SUD, Canceropôle PACA, Conseil Départemental 06, CNRS Biologie, the ANR RENEW (ANR-20-CE13-014) and GIS FC3R (via Inserm). We acknowledge the ImagoSeine core facility of Institut Jacques Monod, member of France-BioImaging (ANR-10-INBS-04) and IBiSA, with the support of Labex “Who Am I”, Inserm Plan Cancer, Region Ile-de-France and Fondation Bettencourt Schueller. We acknowledge IRCAN’s Molecular and Cellular Core Imaging (PICMI) Facility. PICMI was supported financially by FEDER, Conseil régional Provence Alpes-Côte d’Azur, Conseil départemental 06, Cancéropôle PACA, Gis Ibisa and INSERM. We thank the Institut Jacques Monod for its support (6 months) for AV’s fourth year of PhD. We thank Gabriel Krasovec for his helpful suggestions regarding the structure of this manuscript. We thank Noria Izard (intern) for her contribution to the characterization of APO effects in *P. dumerilii*. We thank all past and present members of both teams for their stimulating exchanges surrounding this project.

## Authors contribution

EG, ER and MV designed the study with further contribution from AV and AA. AV performed the comparative genomic analysis part, led by MV and EG. AV performed the wet lab work on *Platynereis*, led by EG, and with contribution from LB and CR. AV and AA conducted the wet lab work on *Nematostella*, led by ER, and with contribution from DD. AV, AA, EG and ER analyzed the data. EG, ER and MV provided financial support. AV, EG and ER wrote the manuscript with contribution from AA. All authors read and approved the final version of the manuscript.

## Data Availability

All data needed to evaluate the conclusions in this study are present in the paper and the Supplementary Materials. Any requests can be addressed to the corresponding authors EG and ER.

## Supplementary tables

**Supplementary Table 1: Species of interest**

First sheet provides taxonomic information. Second sheet indicates the source (website and associated publications) of sequence datasets (predicted proteins or transcripts).

**Supplementary Table 2: Domain structure of Nrf / Keap1 sequences**

The table contains schematic linear representations of putative Nrf (first tab) and Keap1 (second tab) sequences found in all studied species, indicating their domain composition and the position of DLG / ETGE motifs (Nrf) or key cysteines (Keap1). Legends for colours and Interproscan references are found at the bottom of the sheet. bZIP domains were aligned to make up for the **varying** length of Nrf sequences.

**Supplementary Table 3: Expression data of ROS metabolism genes in metazoans**

List of genes, accession numbers and average expression level data for the heatmaps in Figure 6. Each tab corresponds to one species: *Paryhale hawaiensis* (Sinigaglia et al. 2022), *Xenopus tropicalis* (Chang et al. 2017), *Danio rerio* (Nauroy et al. 2019), *Hydra vulgaris* (Wenger et al. 2019), *Nematostella vectensis* (Warner et al. 2018), *Platynereis dumerilii* (Paré et al. 2023).

**Supplementary Table 4: ROS metabolism genes in Platynereis and Nematostella**

List of ROS metabolism genes identified in *Platynereis* and *Nematostella* with identifiers from Paré et al. 2023 and the NvERTx database (Warner et al. 2018) respectively.

## Supplementary figures

**Supplementary Figure 1:**
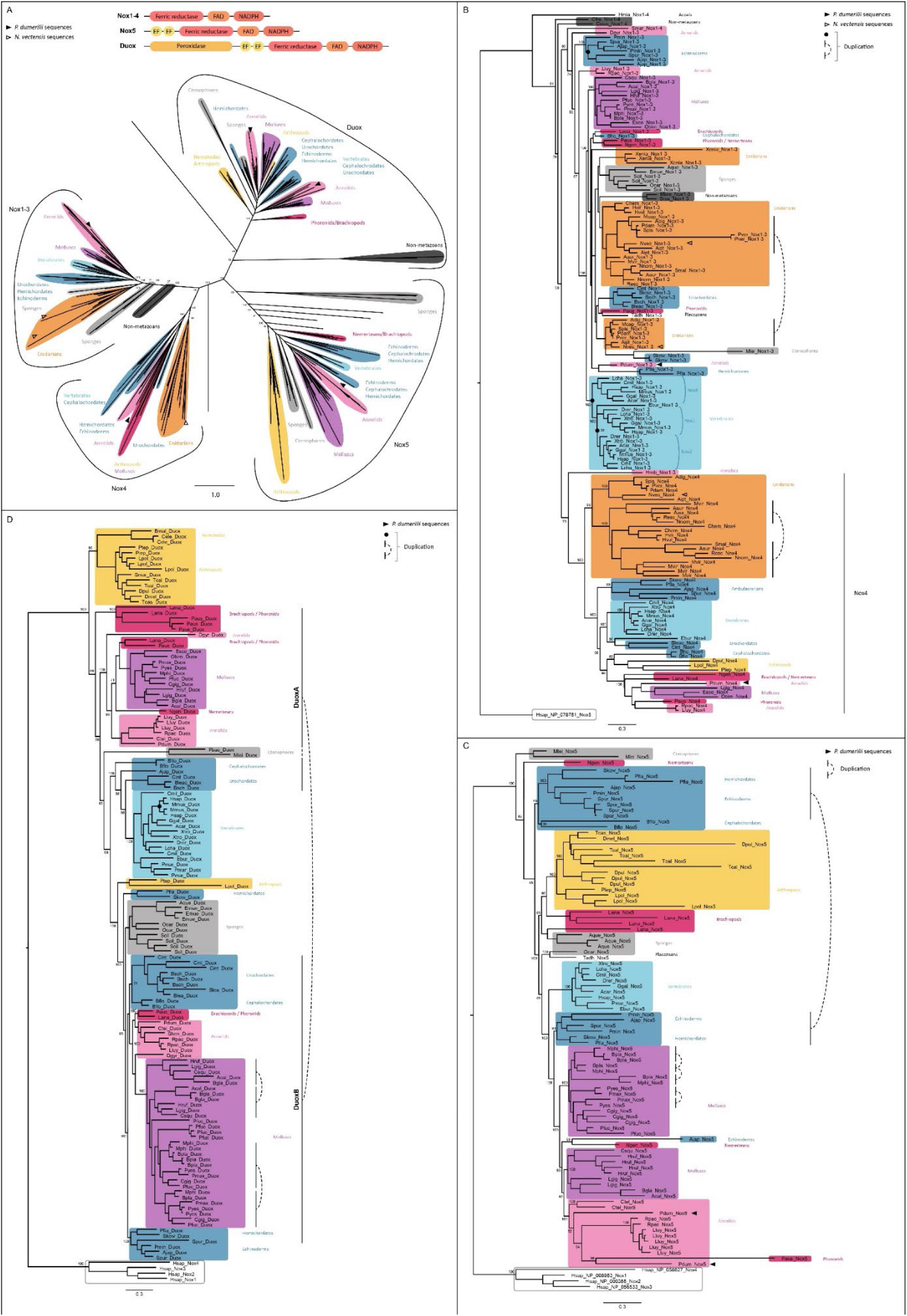
Phylogenetic analysis of the *Nox/Duox* gene families. For all gene families, Maximum likelihood (ML) trees were reconstructed with PhyML, statistical support (aBayes values) is indicated for the deeper nodes and *Platynereis* and *Nematostella* sequences are indicated by plain and empty arrowheads respectively. (A) Global phylogeny of Nox and Duox sequences (unrooted tree). Schematics for the domain composition of Nox1-4, Nox5 and Duox proteins are shown (top). A phylogenetic analysis using only the Nox domain of each sequence yielded a similar topology (data not shown). Since the Nox/Duox is a multigenic family and no non-metazoan sequence could be used as a suitable outgroup, the tree was left unrooted. Amino acids substitution model by SMS (Q.pfam +R+F), Bayesian Information Criterion. (B) Phylogenetic analysis of Nox1-3 and Nox4 sequences. A human Nox5 sequence was used for rooting. Nox4 sequences are highlighted. Duplications in cnidarians, vertebrates and echinoderms are indicated either by a black dot at the node or by a dotted bracket when a single node could not be pinpointed. Amino acids substitution model by SMS (LG +G+I+F), Bayesian Information Criterion. (C) Phylogenetic analysis of Nox5 sequences. Human Nox1-4 sequences were used for rooting. Duplications in molluscs are indicated by a dotted bracket. Amino acids substitution model by SMS (LG +G), Bayesian Information Criterion. (D) Phylogenetic analysis of Duox sequences. Human Nox1-4 sequences were used for rooting. Duplications are indicated either by a black dot at the node or by a dotted bracket when a single node could not be pinpointed. Amino acids substitution model by SMS (LG +G+F), Bayesian Information Criterion.

**Supplementary Figure 2:**
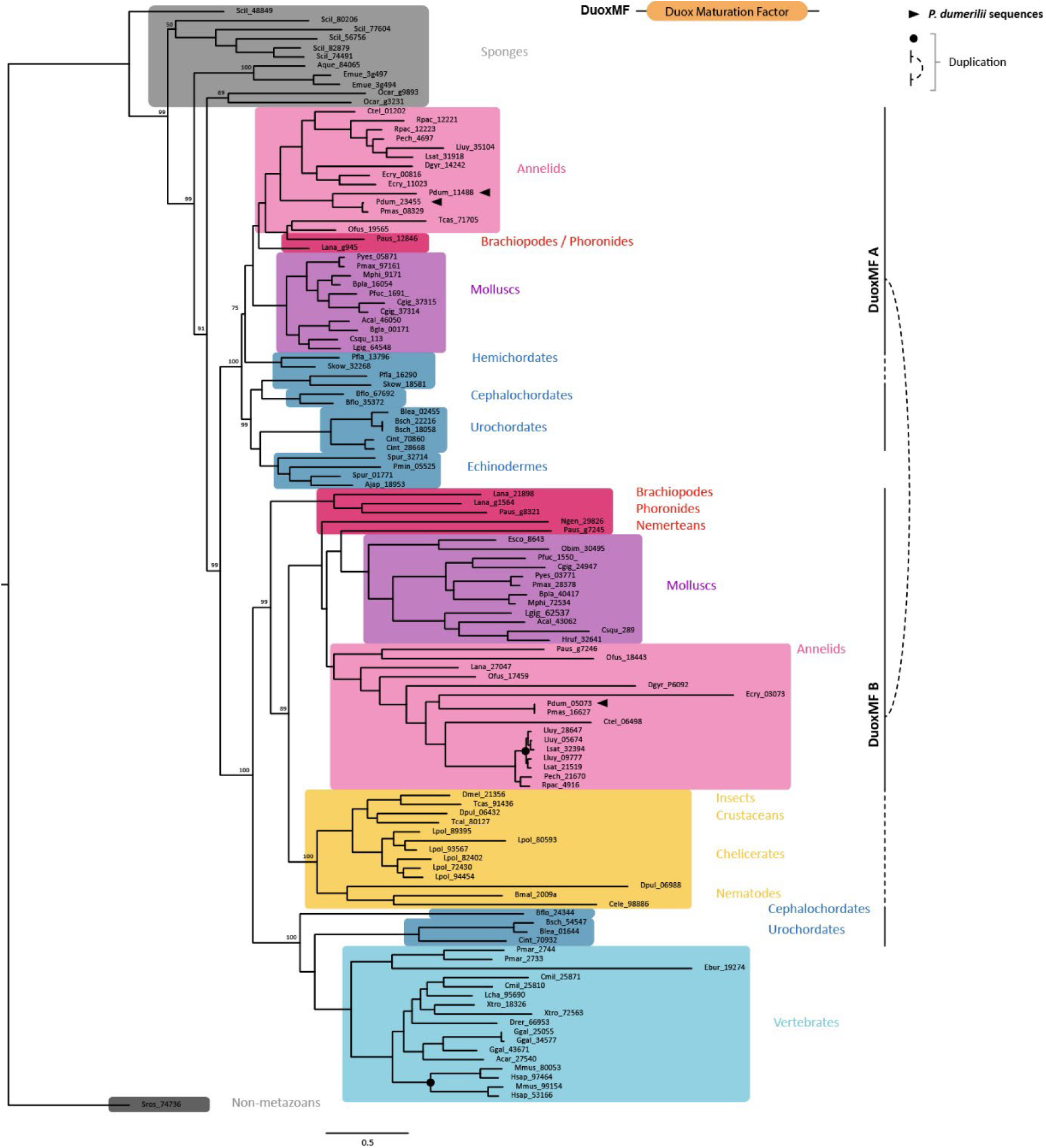
Phylogenetic analysis of the *Duox Maturation Factor* gene family. Maximum likelihood (ML) tree constructed with PhyML is shown. A non-metazoan DuoxMF sequence was used for rooting. Statistical support (aBayes values) is indicated for the deeper nodes. Duplications are indicated either by a black dot at the node or by a dotted bracket when a single nod could not be pinpointed. *Platynereis* sequences are indicated by black arrowheads. Amino acids substitution model by SMS (LG +R+F), Bayesian Information Criterion.

**Supplementary Figure 3:**
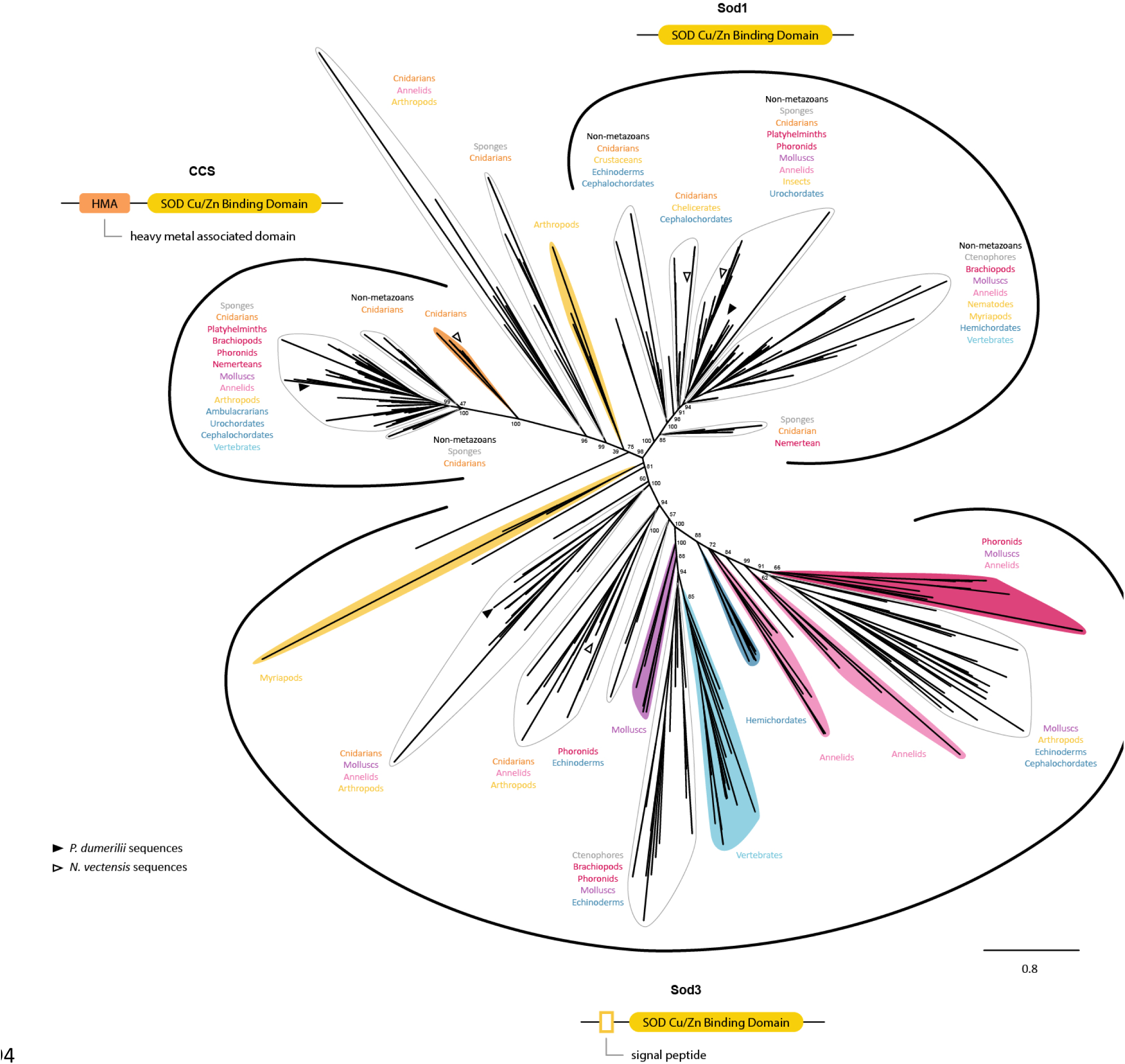
Phylogenetic analysis of the *Sod Cu/Zn* and *CCS* gene families. Maximum likelihood (ML) tree constructed with PhyML is shown. Since the Sod Cu/Zn appear to be a multigenic family, and no single group of non-metazoan sequences could be used as a suitable outgroup, the tree was left unrooted, with three main groups displayed. Statistical support (aBayes values) is indicated for the deeper nodes. Ancestral duplications are indicated either by a dot at the node or by a dotted bracket when a single node could not be pinpointed. *Platynereis* and *Nematostella* sequences are indicated by plain and empty arrowheads respectively. Amino acids substitution model by SMS (WAG +R), Bayesian Information Criterion.

**Supplementary Figure 4:**
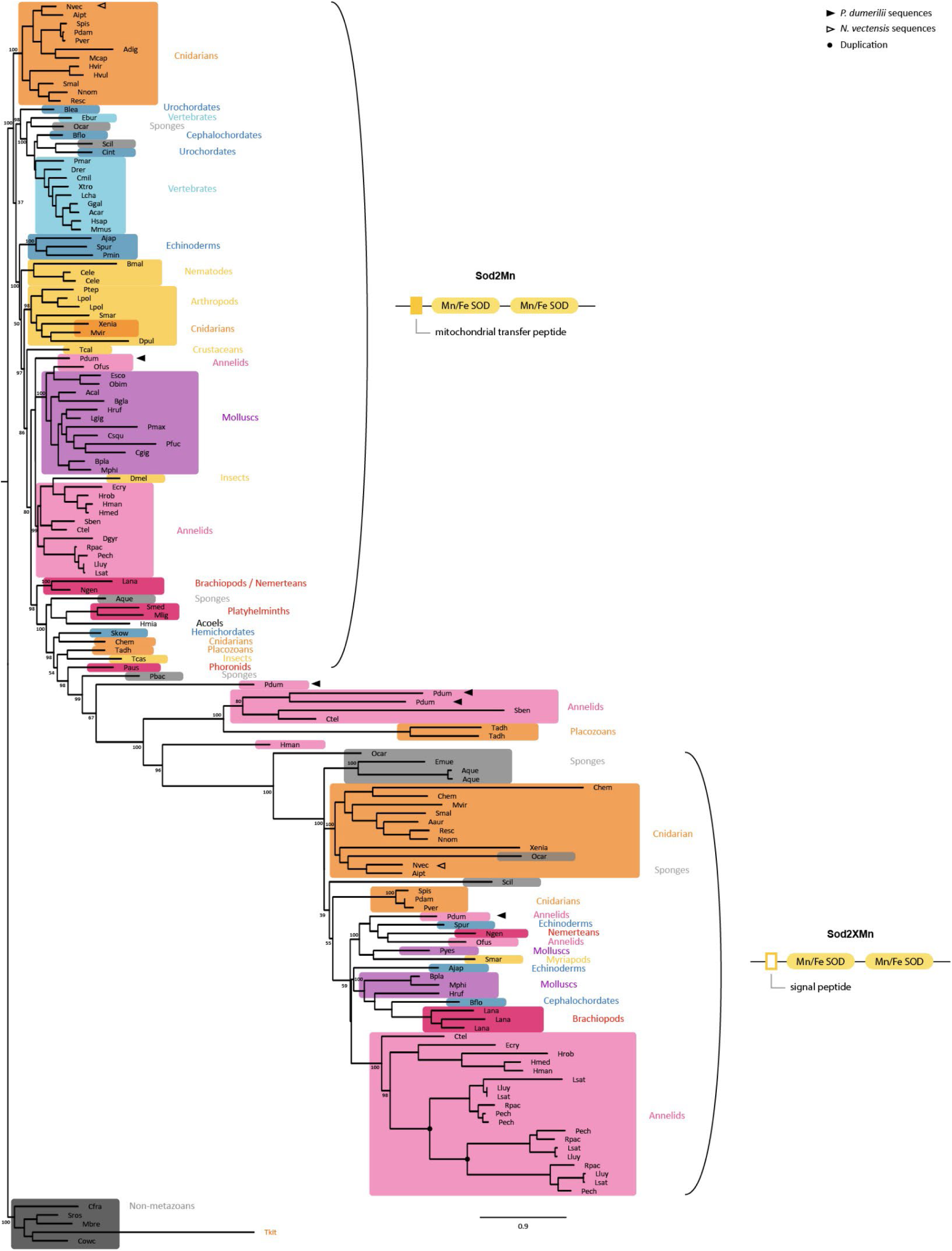
Phylogenetic analysis of the *SodMn* gene family. Maximum likelihood (ML) tree constructed with PhyML is shown. Non-metazoan sequences were used for rooting. Statistical support (aBayes values) is indicated for the deeper nodes. Ancestral duplications are indicated by a black dot at the nodes. *Platynereis* and *Nematostella* sequences are indicated by plain and empty arrowheads respectively. Amino acids substitution model by SMS (Q.pfam +G+I), Bayesian Information Criterion.

**Supplementary Figure 5:**
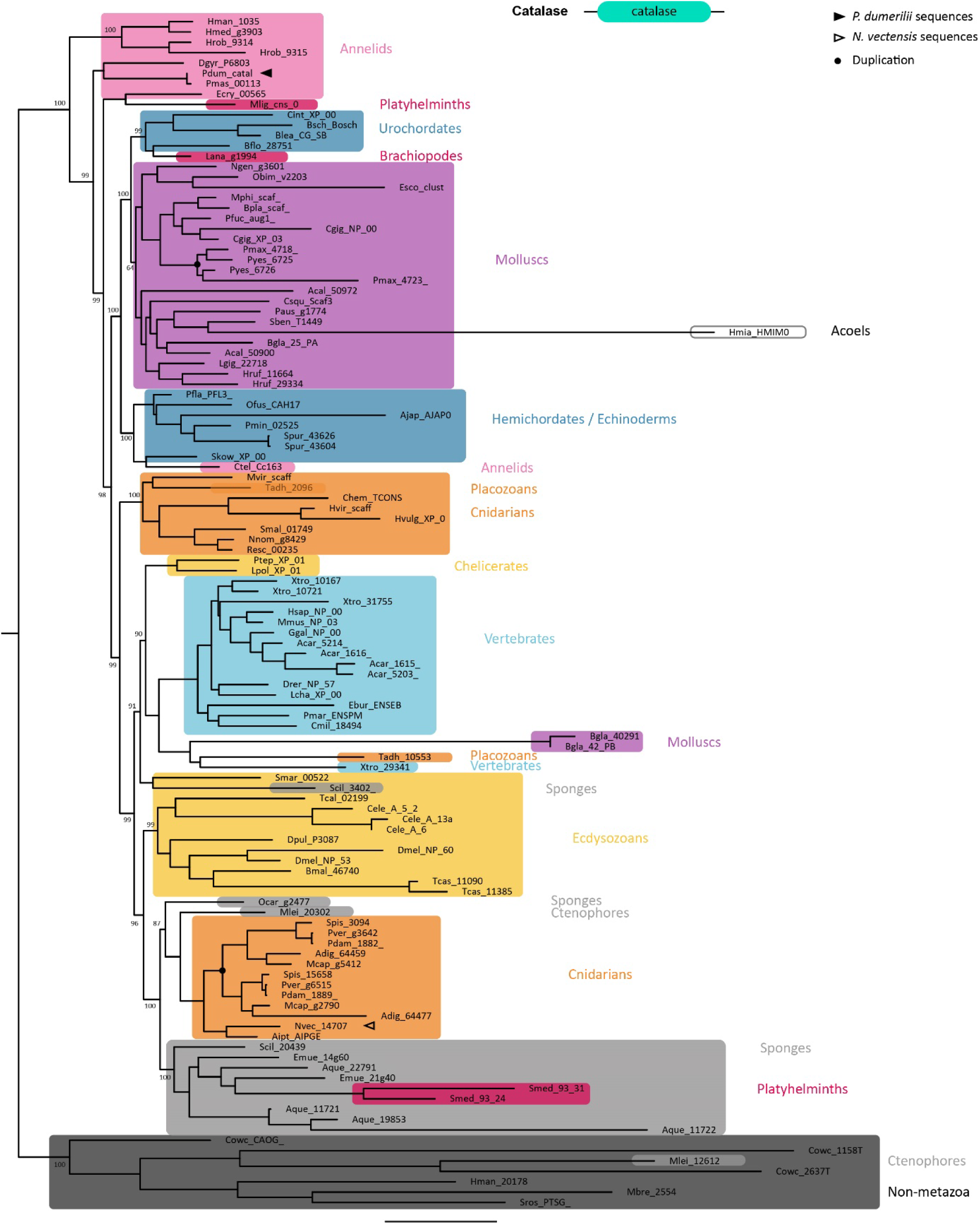
Phylogenetic analysis of the *catalase* gene family. A Maximum likelihood (ML) tree, rooted on non-metazoan sequences is shown. Statistical supports (aBayes values) are indicated for the deeper nodes. Duplication in scallops and Scleractinia are indicated by a black dot at the nodes. *Platynereis* and *Nematostella* sequences are indicated by plain and empty arrowheads respectively. Amino acids substitution model by SMS (LG +R), Bayesian Information Criterion.

**Supplementary Figure 6:**
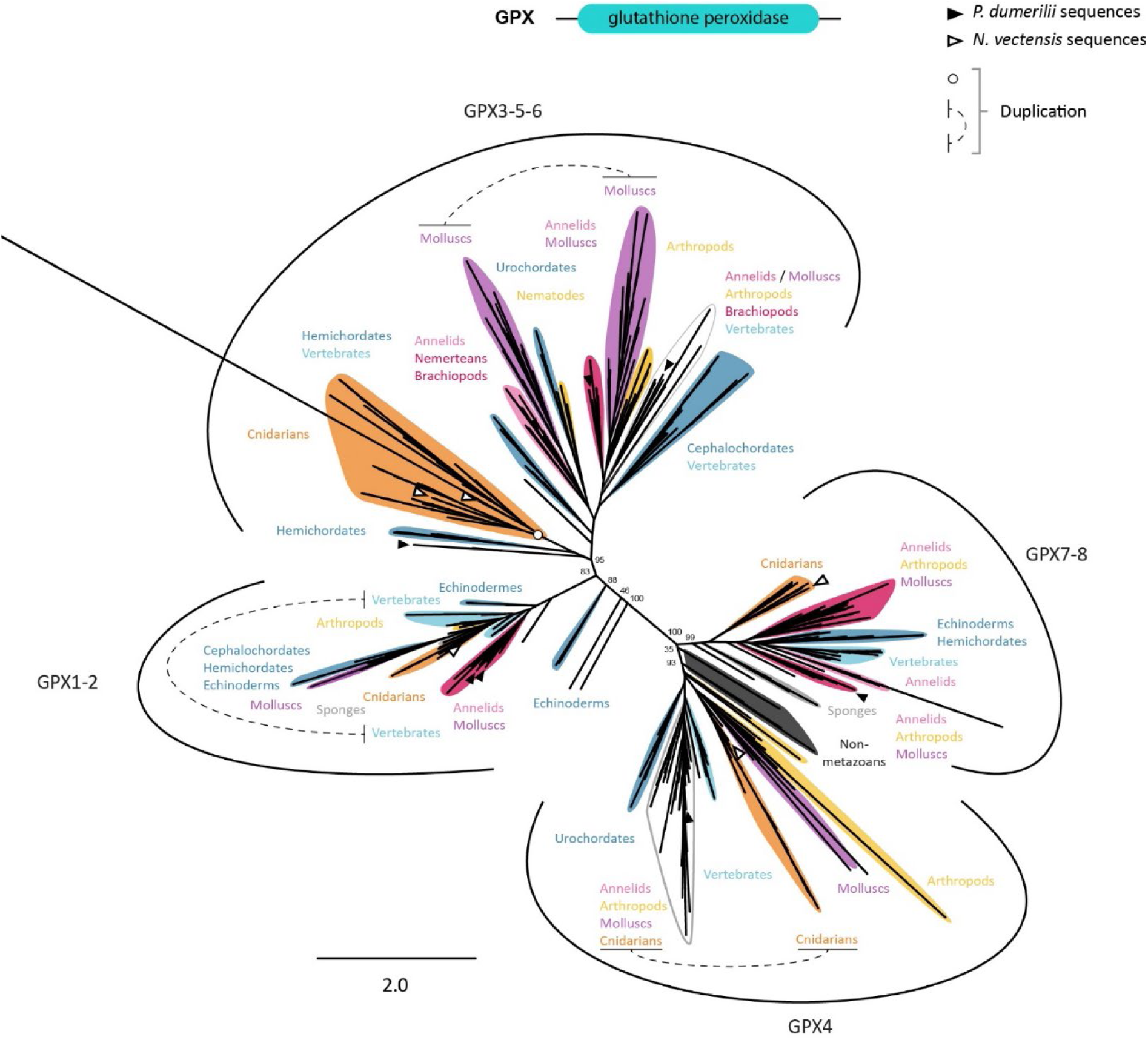
Phylogenetic analysis of the *GPX* gene family. A Maximum likelihood (ML) tree is shown. Since the GPX are a multigenic family and no group of non-metazoan sequences could be used as a suitable outgroup, the tree was left unrooted, with four main groups displayed. Statistical support (aBayes values) is indicated for the deeper nodes. *Platynereis* and *Nematostella* sequences are indicated by plain and empty arrowheads respectively. Amino acids substitution model by SMS (WAG +R), Bayesian Information Criterion.

**Supplementary Figure 7:**
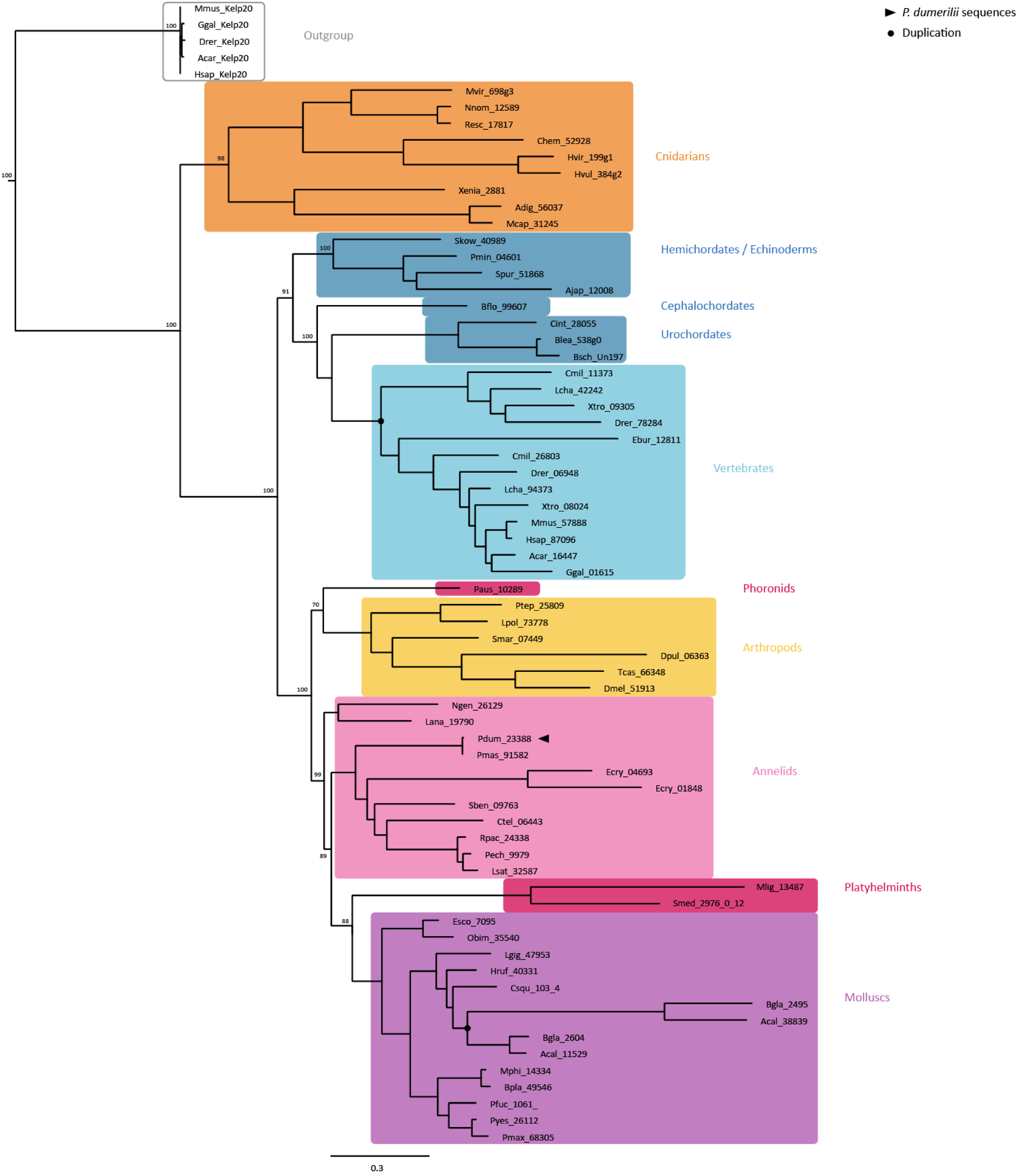
Phylogenetic analysis of the *Keap1* gene family. A Maximum likelihood (ML) tree constructed with PhyML is shown. Vertebrate Kelp20 sequences were used for rooting. Statistical support (aBayes values) is indicated for the deeper nodes. Ancestral duplications are indicated by a black dot at the node. *Platynereis* sequences are indicated by black arrowheads. Amino acids substitution model by SMS (LG +R+F), Bayesian Information Criterion.

**Supplementary Figure 8:**
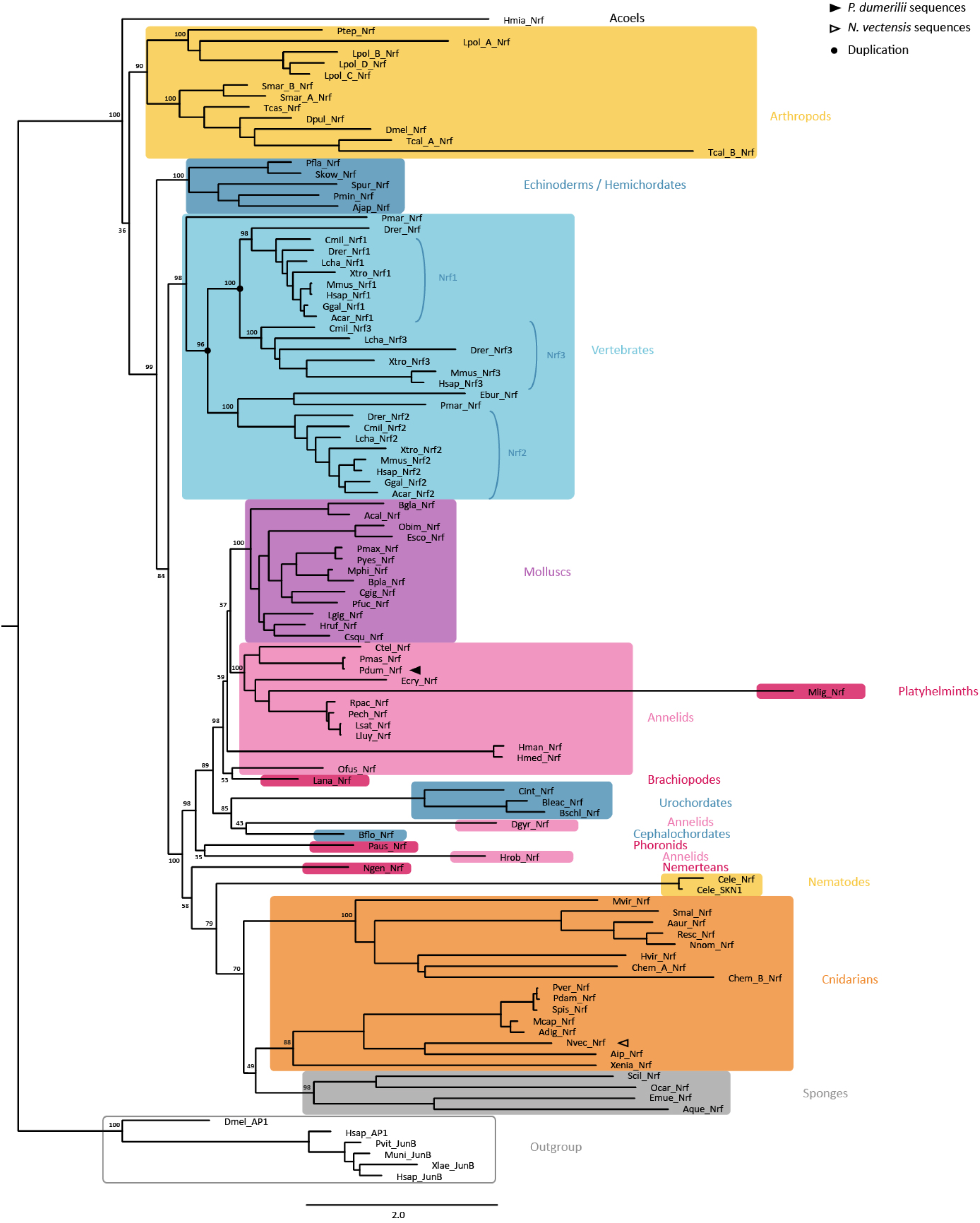
Phylogenetic analysis of the *Nrf* gene family. A Maximum likelihood (ML) tree constructed with PhyML is shown. bZIP protein sequences (AP1 and JunB) were used for rooting. Statistical support (aBayes values) is indicated for the deeper nodes. Ancestral duplications are indicated by a black dot at the node. *Platynereis* and *Nematostella* sequences are indicated by plain and empty arrowheads respectively. Amino acids substitution model by SMS (Q.insect +R+F), Bayesian Information Criterion.

**Supplementary Figure 9:**
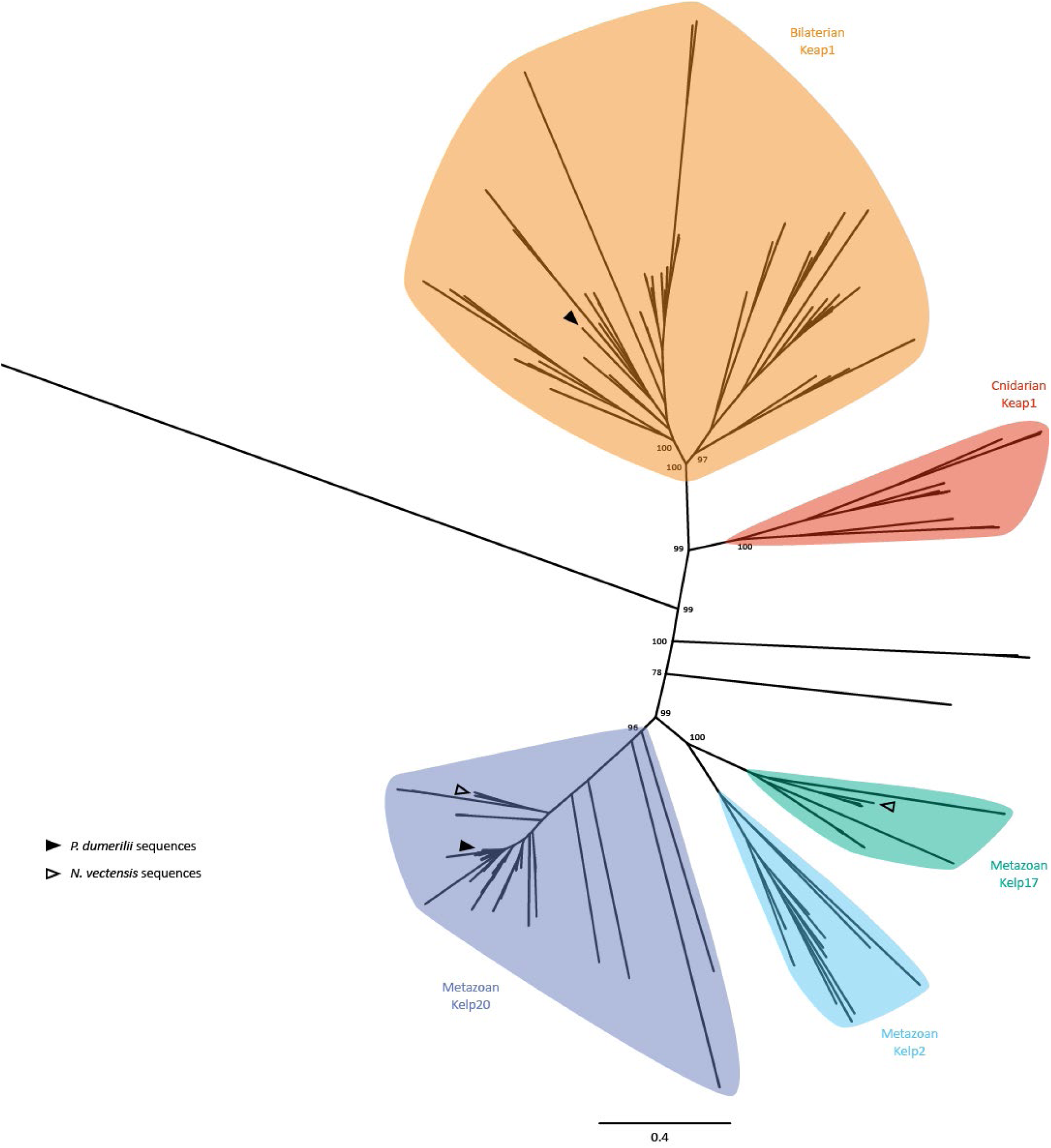
Phylogenetic analysis of the *Keap1* and *Kelp* gene families. A Maximum likelihood (ML) tree constructed with PhyML is shown. Keap1 sequences were aligned with a subset of Kelp20, Kelp2 and Kelp17 sequences from all main metazoan lineages to clarify the position of cnidarian putative Keap1 sequences. Since the tree displays multiple gene families it was left unrooted. Statistical support (aBayes values) is indicated for the deeper nodes. *Platynereis* and *Nematostella* sequences are indicated by plain and empty arrowheads respectively. Amino acids substitution model by SMS (LG +G+I+F), Bayesian Information Criterion.

**Supplementary Figure 10:**
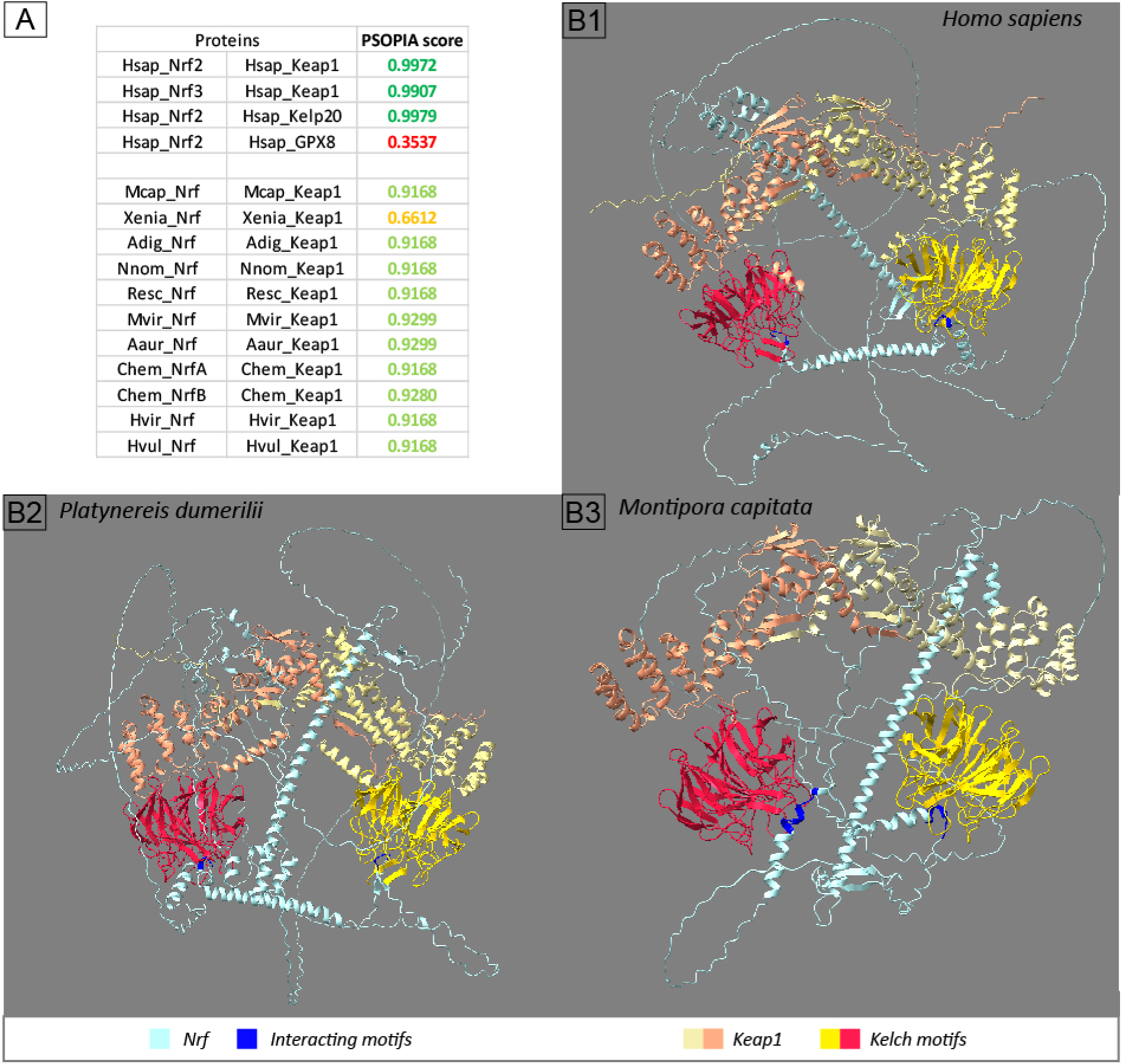
Structure prediction of Nrf/Keap1 interactions. (A) Protein interaction scores by PSOPIA based on Nrf and Keap1 protein sequences. Hsap: Homo sapiens. Mcap: *Montipora capitata*. Xenia: *Xenia sp*. Adig: *Acropora digitifera*. Nnom: *Nemopilema nomurai*. Resc: *Rhopilema esculentum*. Mvir: *Morbakka virulenta*. Aaur: *Aurelia aurita*. Chem: *Clytia hemisphaerica*. Hvir: *Hydra viridissima*. Hvul: *Hydra vulgaris*. (B) Predicted conformations of Nrf / Keap1 association by AlphaFold in *Homo sapiens* (B1), *Platynereis dumerilii* (B2) and *Montipora capitata* (B3).

**Supplementary Figure 11:**
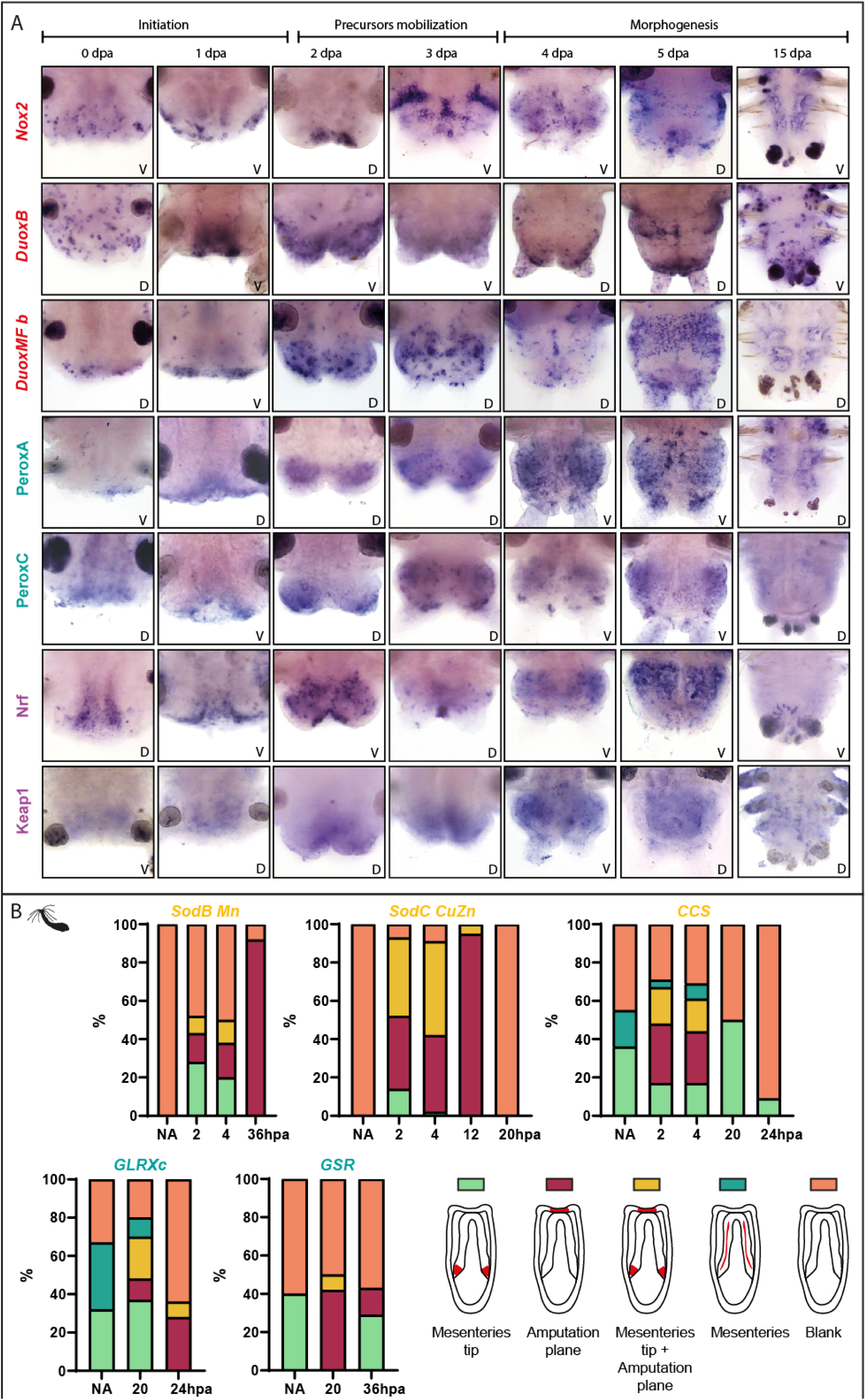
Additional expression patterns of ROS metabolism genes during regeneration. (A) Whole-mount *in situ* hybridizations of chosen genes at different stages of *Platynereis* posterior regeneration. V: ventral view. D: dorsal view. (B) Proportions of different expression patterns for 5 ROS metabolism genes over the course of oral regeneration in *Nematostella*.

**Supplementary Figure 12:**
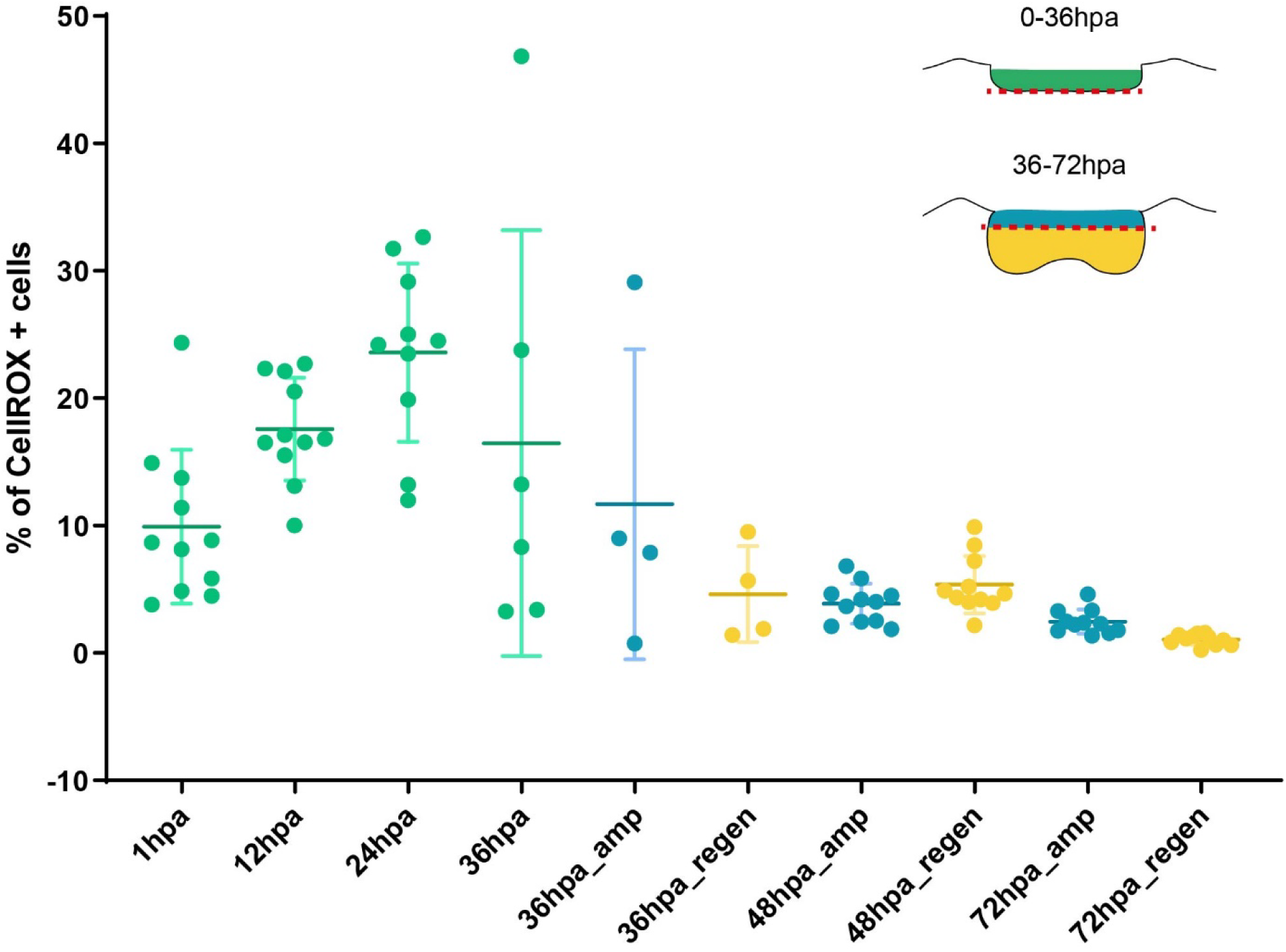
Detailed CellROX+ counting in *Platynereis* posterior regeneration. Detail of CellROX+ cells quantification method. For 0-24hpa samples, and 36hpa samples displaying little regenerated tissue, counting was performed in a 30µm wide volume abutting the amputation plane (green). For 48-72hpa samples, and 36hpa samples displaying a small blastema, counting was performed in a 30µm wide volume abutting the amputation plane (blue, _amp) and in the regenerated structures (yellow, _regen). In a spirit of simplification, _regen and _amp counts of each 36, 48 and 72hpa samples were pooled together in Figure 8B, as this did not alter the overall kinetic of ROS production.

**Supplementary Figure 13:**
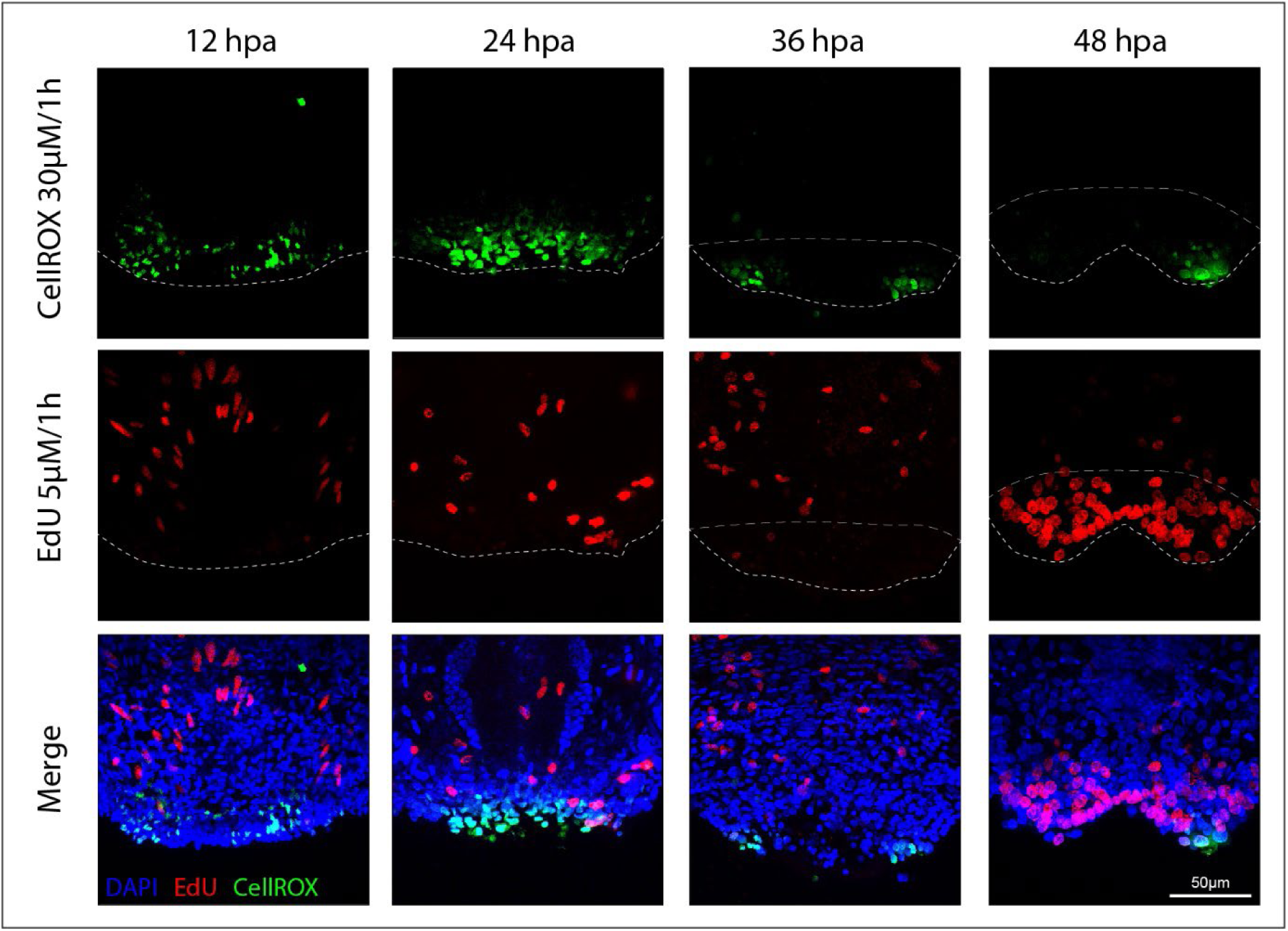
ROS production and proliferation double labelling in *Platynereis.* Confocal Z-stacks projections of ROS production (CellROX, green) and proliferation (EdU, red) during the first 48 hours of posterior regeneration in *Platynereis*. Ventral views, anterior is up, nuclei labelling in blue.

**Supplementary Figure 14:**
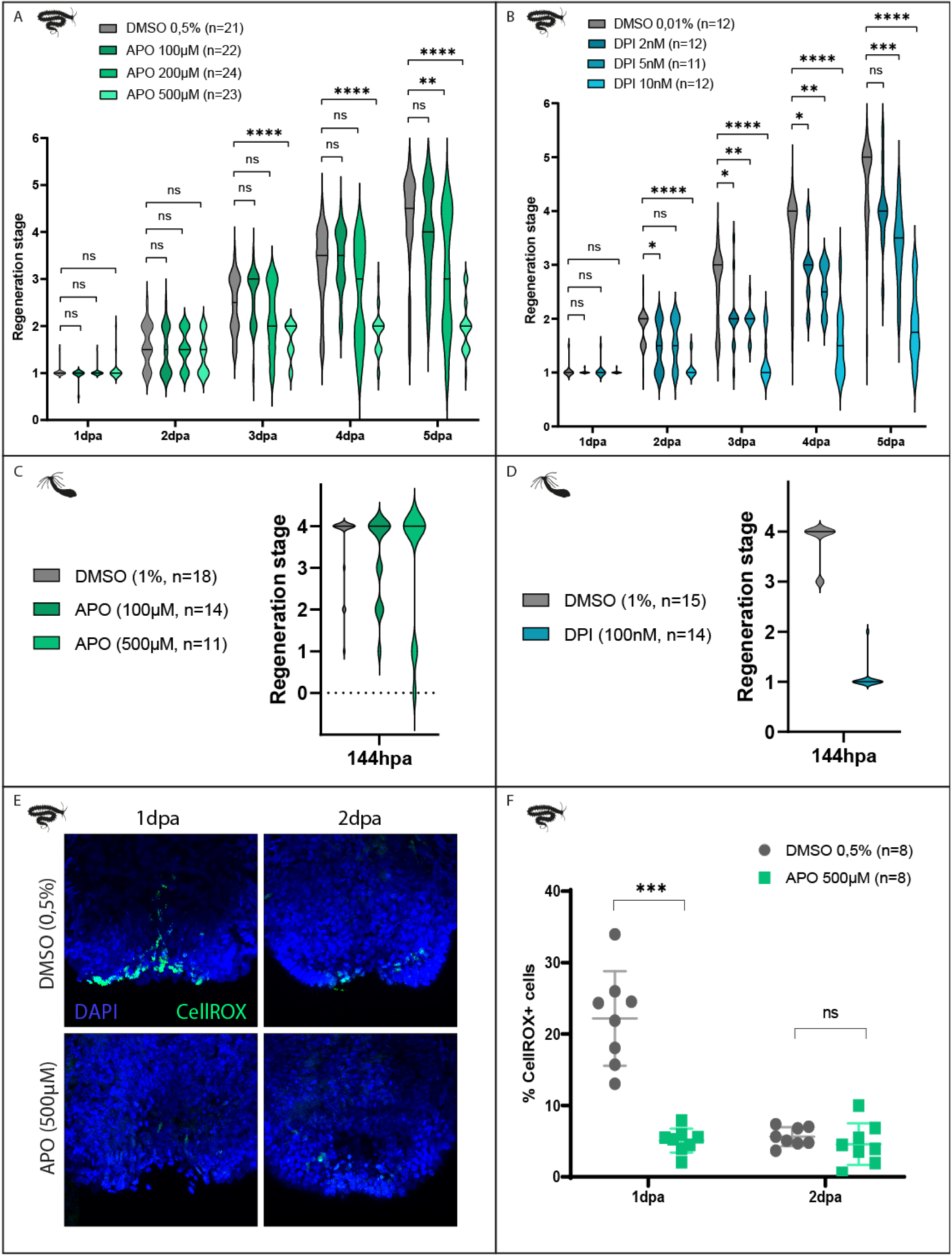
Efficient concentration and validation of APO and DPI. (A, B) Scoring of regenerating worms treated with different APO (A) and DPI (B) concentrations over 5 days post-amputation in order to identify efficient concentrations. Brackets correspond to multiple Mann-Whitney tests (ns p>0.05; * p≤0.05; ** p≤0.01; *** p≤ 0.001; **** p≤ 0.0001). (C) ROS production labelling (CellROX, green) at 1 and 2dpa under APO 500µM or control treatment. Ventral views, anterior is up, nuclei labelling in blue. (D) Quantification of the proportion of CellROX positive cells in the regenerating posterior part at 1 and 2dpa upon APO 500µM or DMSO 0,5% treatment. Each bracket corresponds to a Mann-Whitney U test (ns p >0.05; *** p≤0.001). Mean + SD are shown.

**Supplementary Figure 15:**
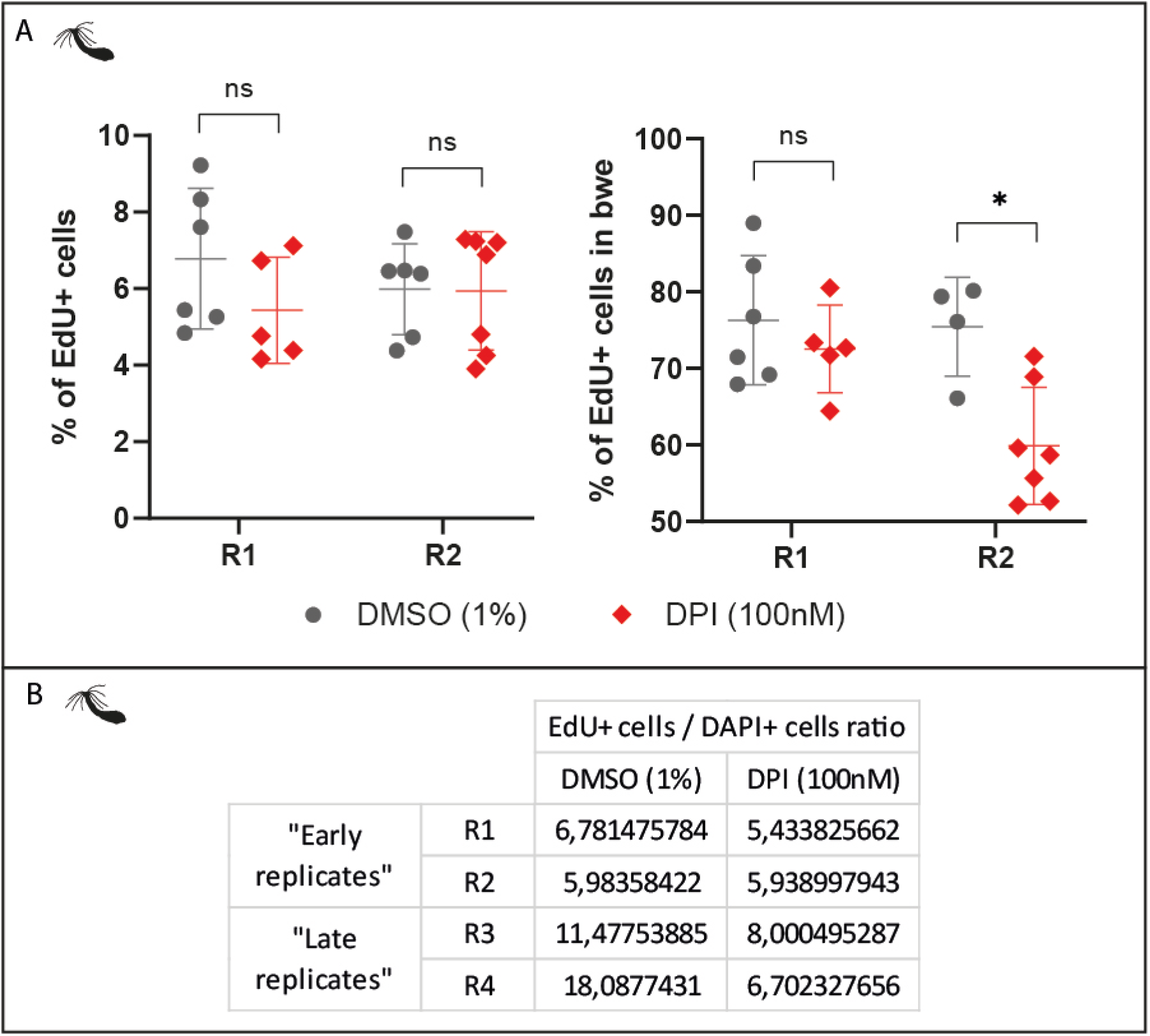
(A) Proportion of EdU positive cells in the *Nematostella* regenerating oral part (left) and proportion of EdU+ cells found in the body wall epithelium (bwe) (right) at 48hpa under DPI 100nM treatment and DMSO 0.1% control (R1 and R2). Each bracket corresponds to a Mann-Whitney U test (ns p >0.05; * p≤0.05). (B) EdU+/DAPI+ cells ratios in control and DPI-treated conditions for all four replicates.

**Supplementary Figure 16:**
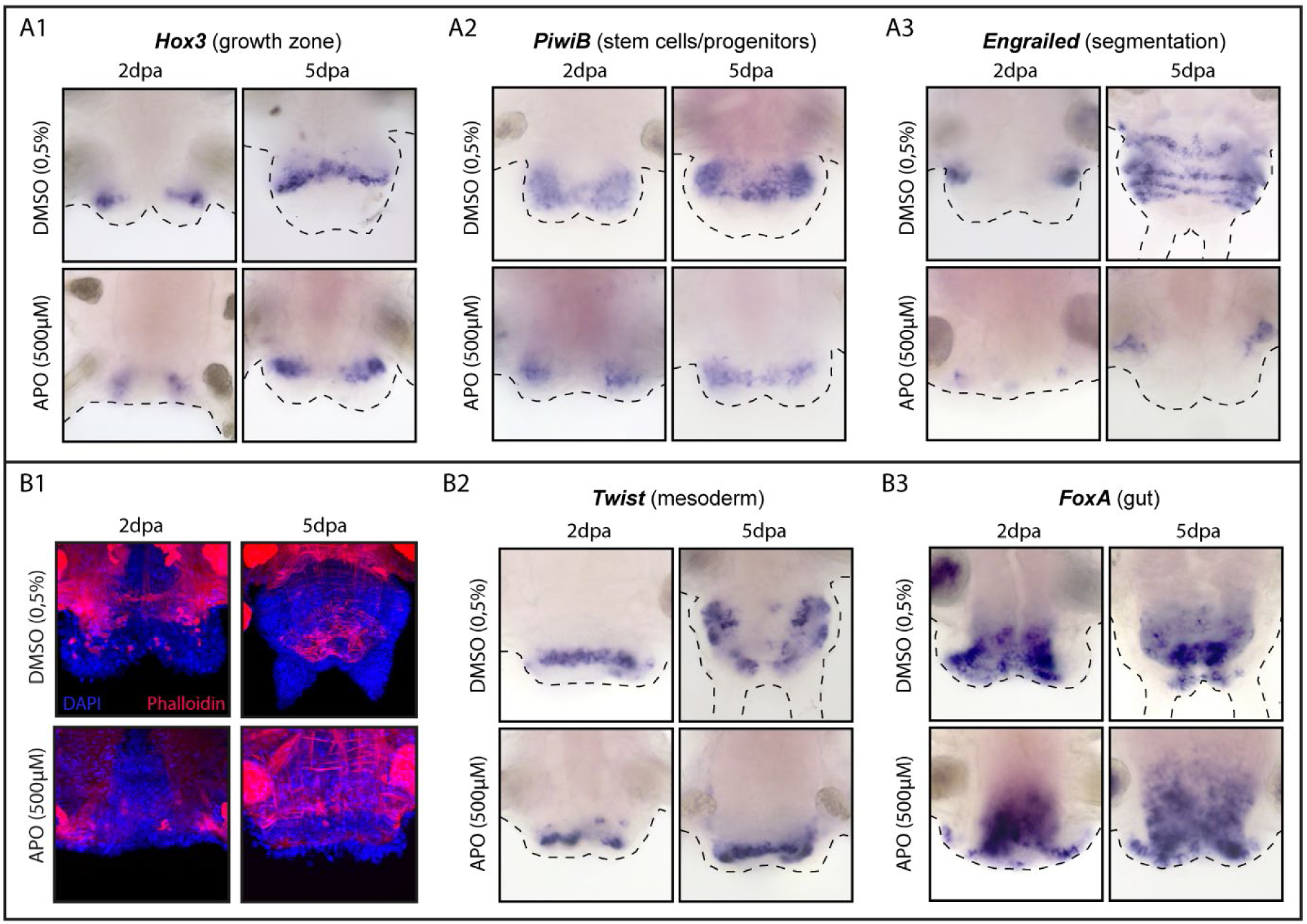
Phenotypic characterization of ROS inhibition during Platynereis regeneration. (A) Growth zone activity and segmentation: Whole-mount in situ *hybridizations for markers of the growth zone (*Hox3*, A1), progenitors (*PiwiB*, A2) and segmentation (*Engrailed*, A3) at 2 and 5 dpa for APO-treated worms and controls. (B) Meso-endoderm identity: Phalloidin labelling (B1) and whole-mount* in situ *hybridizations for markers of the mesoderm (*Twist*, B2) and gut (*FoxA*, B3) at 2 and 5 dpa for APO-treated worms and controls. Ventral views. Anterior is up*.

